# Structure and mechanism of a novel cytomegaloviral DCAF mediating interferon antagonism

**DOI:** 10.1101/2022.05.05.490734

**Authors:** Vu Thuy Khanh Le-Trilling, Sofia Banchenko, Darius Paydar, Pia Madeleine Leipe, Lukas Binting, Simon Lauer, Andrea Graziadei, Christine Gotthold, Jörg Bürger, Thilo Bracht, Barbara Sitek, Robert Jan Lebbink, Anna Malyshkina, Thorsten Mielke, Juri Rappsilber, Christian M. T. Spahn, Sebastian Voigt, Mirko Trilling, David Schwefel

**Author notes:** These authors contributed equally (Vu Thuy Khanh Le-Trilling, Sofia Banchenko). Correspondence (S.V.), (M.T.), (D.S.).

## Abstract

Human cytomegalovirus (CMV) is a highly relevant and ubiquitously distributed human pathogen. Its rodent counterparts such as mouse and rat CMV serve as common infection models. Here, we conducted the first global proteome profiling of rat CMV-infected cells and uncovered a pronounced loss of the transcription factor STAT2, which is crucial for interferon signalling. Deletion mutagenesis documented that STAT2 is targeted by the viral protein E27. Cellular and *in vitro* analyses showed that E27 exploits host-derived Cullin4-RING ubiquitin ligases (CRL4) to induce poly-ubiquitylation and proteasomal degradation of STAT2. A cryo-electron microscopic structure determination revealed how E27 mimics molecular surface properties of cellular CRL4 substrate receptors called *DDB1- and Cullin4-associated factors* (DCAFs) to displace them from the catalytic core of CRL4. Moreover, structural analyses elucidated the mechanism of STAT2 recruitment and indicate that E27-binding additionally disturbs STAT2-dependent interferon signalling by occupying its IRF9 binding interface. For the first time, these data provide structural insights into cytomegalovirus-encoded interferon antagonism and establish an atomic model for STAT2 counteraction by CRL4 misappropriation with important implications for viral immune evasion.

## Introduction

Human cytomegalovirus (HCMV; Human herpesvirus 5) is the prototypic member of the *Betaherpesvirinae* and a ubiquitous human pathogen. More than three quarters of the global population are latently infected with HCMV (Virgin et al., 2009). HCMV infections usually progress subclinically in healthy adults, but cause morbidity and mortality in immunocompromised individuals such as newborns, AIDS patients, transplant recipients, and the elderly (Griffiths and Reeves, 2021). During host adaptation, cytomegaloviruses became strictly species-specific. Therefore, meaningful *in vivo* experiments cannot be performed with HCMV in unmodified small animal models. This roadblock for basic research and preclinical development is circumvented by studying closely related betaherpesviruses such as mouse cytomegalovirus (MCMV; Murid herpesvirus [MuHV] 1) and rat cytomegaloviruses (RCMVs; MuHV-2 and MuHV-8) (Brizic et al., 2018; Geyer et al., 2015; Voigt et al., 2007) in their corresponding host species. Since MuHV infections represent some of the few genuine virus infection models in which the animal host is infected with its natural pathogen (Becker et al., 2007), they became well-established and broadly applied models in virology and immunology, enabling insights regarding general phenomena of virus pathogenesis and host immunity (Reddehase and Lemmermann, 2018).

Upon encounter of pathogens such as cytomegaloviruses, cytokines called interferons (IFNs) are expressed (Ivashkiv and Donlin, 2014; Panne et al., 2007; Schneider et al., 2008) which elicit potent antiviral activity (Isaacs and Lindenmann, 1957) by changing the cellular transcriptome and proteome (Megger et al., 2017; Trilling et al., 2013). In addition to the antiviral activity executed within infected host cells, IFNs also orchestrate cellular immune responses (Baranek et al., 2012). IFNs bind to cell surface receptors that are associated with *janus kinases* (JAK), which then phosphorylate intracellular receptor chains, generating docking sites for the *signal transducer and activator of transcription* (STAT) (Darnell et al., 1994). After phosphorylation by JAKs, STATs homo- and heterodimerise and form transcription factor complexes that translocate into the nucleus where they induce the expression of IFN-stimulated genes (ISGs) (Levy and Darnell, 2002). Type I IFNs (IFN-I) and type III IFNs (IFN-III) signal through STAT1/STAT2 heterodimers which recruit *IFN regulatory factor 9* (IRF9) to generate the active *IFN-stimulated gene factor 3* (ISGF3) that binds IFN-stimulated response elements (ISRE) in the nucleus (Horvath et al., 1996; Kotenko et al., 2003; Qureshi et al., 1995). IFNγ, the only known type II IFN (IFN-II), mainly signals via phosphorylated STAT1 homodimers that recognise gamma-activated DNA sequences (GAS) and express a distinct set of ISGs. Accordingly, STAT2 is crucial for IFN-I and IFN-III signalling. Furthermore, STAT2 directly and indirectly affects IFN-II signalling, e.g., as part of IFNγ-induced ISGF3 complexes (Le-Trilling et al., 2018; Matsumoto et al., 1999; Trilling et al., 2013; Trilling et al., 2011; Zimmermann et al., 2005), as constituent of unphosphorylated IRF9-STAT2 heterodimers (Blaszczyk et al., 2015; Platanitis et al., 2019), and by regulating STAT1 expression levels (Hambleton et al., 2013; Ho et al., 2016; Park et al., 2000). Accordingly, animals and humans lacking STAT2 are severely immunosuppressed and show an exaggerated susceptibility to infections (Gowen et al., 2017; Hambleton et al., 2013; Le-Trilling et al., 2018; Park et al., 2000).

The pivotal role of STAT2 for IFN signalling and innate immunity shaped virus evolution. Several relevant pathogens escape from an IFN-induced antiviral state by targeting STAT2. For example, Zika virus (ZIKV) (Grant et al., 2016), Dengue virus (DENV) (Ashour et al., 2009; Jones et al., 2005), paramyxoviruses (Parisien et al., 2001; Parisien et al., 2002b), MCMV (Trilling et al., 2011; Zimmermann et al., 2005), and HCMV (Le-Trilling et al., 2020) induce STAT2 degradation along the ubiquitin proteasome pathway (UPS). Intriguingly, the responsible virus proteins act as adapters that place STAT2 into the sphere of influence of cellular E3 ubiquitin ligases (E3s) (Barik, 2022; Le-Trilling and Trilling, 2020; Ulane and Horvath, 2002).

The MCMV protein M27 was the first CMV STAT2 antagonist to be described (Zimmermann et al., 2005). MCMV mutants lacking *M27* are severely attenuated *in vivo* (Le-Trilling et al., 2018; Zimmermann et al., 2005) due to the inability to interfere with STAT2-dependent IFN signalling as evident from the restored replication in STAT2-deficient mice (Le-Trilling et al., 2018). Mechanistically, M27 binds DDB1, a component of host Cullin4-RING E3s (CRL4), to induce poly-ubiquitination of STAT2, resulting in its rapid proteasomal degradation (Landsberg et al., 2018; Trilling et al., 2011). While the aforementioned complex formation has been confirmed in several models and by different techniques, direct binding of M27 to DDB1 and/or STAT2 in absence of other proteins has to our knowledge not been documented yet. This is relevant since paramyxoviral IFN antagonists bind one STAT and induce degradation of another, e.g., STAT1 is needed by human parainfluenza virus 2 to target STAT2 while STAT2 is required for both simian virus 5 (SV5) and mumps V proteins to target STAT1 (Ulane et al., 2005). Structural details regarding the interactions between CRL4, STAT2, and CMV antagonists are currently not available.

CRL4, similar to other CRLs, are modular complexes, comprising a core formed by the cullin scaffold (CUL4A or CUL4B) and a catalytic RING domain subunit (RBX1). By the adapter protein DDB1, the core is connected to various exchangeable substrate receptors called DCAFs, forming various cellular CRL4 E3s with different specificities (Zimmerman et al., 2010). CRLs are regulated by cycles of activation through modification of the cullin C-terminus with NEDD8 (“neddylation”), followed by de-neddylation of substrate-free CRLs and substrate receptor exchange (Baek et al., 2020; Cavadini et al., 2016; Goldenberg et al., 2004; Harper and Schulman, 2021; Reichermeier et al., 2020). Among CRLs, CRL4 exhibit unique features like the presence of the co-adaptor DDA1, which stabilises at least certain substrate receptors on CRL4 by bridging them to DDB1, as in the case of DCAF15 (Bussiere et al., 2020; Du et al., 2019; Faust et al., 2020; Olma et al., 2009; Shabek et al., 2018), and the existence of a flexible hinge in the DDB1 adapter that allows for rotation of the catalytic subunit around the substrate receptor, thus creating a broad ubiquitylation zone to accommodate substrates of different size and shape (Banchenko et al., 2021; Fischer et al., 2014; Fischer et al., 2011). Most likely, the latter property makes CRL4 a frequent target for viral exploitation to steer antiviral host factors efficiently into the ubiquitin-proteasome pathway. In addition to CMV M27, there are further examples of viral CRL4-binders that induce host protein degradation such as CMV RL1 (Nightingale et al., 2022; Nobre et al., 2019), UL35 (Salsman et al., 2012), UL145 (Le-Trilling et al., 2020), hepatitis B virus X (HBx) (Decorsiere et al., 2016), paramyxovirus V - the first viral CRL4 hijacker to be structurally analysed (Li et al., 2006; Precious et al., 2005), and the accessory proteins Vpx and Vpr from immunodeficiency viruses (Greenwood et al., 2019; Schwefel et al., 2015; Schwefel et al., 2014). Mechanistic studies of these viral CRL4 subversion processes yield direct molecular insight into virus adaptation to host replication barriers, advancing the understanding of virus pathology and potentially paving the way for therapeutic opportunities (Becker et al., 2019).

Here, we pursued a proteomic approach to assess RCMV-dependent protein regulation and identified STAT2 as one of the most thoroughly down-regulated factors. The RCMV protein E27 was necessary for reduction of STAT2 protein levels by a process sensitive to the inhibition of the proteasome or CRLs. Moreover, biochemical reconstitution demonstrated that E27 was sufficient for recruitment of STAT2 to the DDB1 adapter protein of host CRL4 complexes. Cryogenic electron microscopy (cryo-EM) structural analyses revealed that E27 replaces endogenous CRL4 substrate receptors, resulting in proper positioning of STAT2 for poly-ubiquitylation and subsequent proteasomal destruction to inhibit host IFN signalling.

## Results

### RCMV-E E27 down-regulates host STAT2

To uncover RCMV-induced proteome changes, rat embryo fibroblasts (REF) were infected with RCMV-England (RCMV-E). The protein content was analysed after 4 h and 28 h by liquid chromatography coupled to mass spectrometry (LC-MS) and compared to mock-infected controls. In each condition, more than 2,300 proteins were quantified in six replicates (Fig. S1A). To our knowledge, this represents the first proteome profiling of RCMV-infected cells. Gene ontology (GO) analysis of the data revealed that only very few ISGs were expressed (Fig. 1A). This under-representation of ISGs among RCMV-induced host proteins strongly suggests the expression of potent RCMV-encoded IFN antagonists. In order to obtain mechanistic insights, we further analysed the relative abundance of IFN signalling proteins. The expression levels of JAK1 and TYK2 kinases were similar to mock-infected cells at both time points (Figs. 1A, S1B). Most strikingly, STAT2 protein levels already started to diminish significantly 4 h post-infection and became virtually undetectable after 28 h of infection (Figs. 1A, S1B), indicating a rapid and sustained RCMV-induced loss of STAT2.

**Figure 1:**
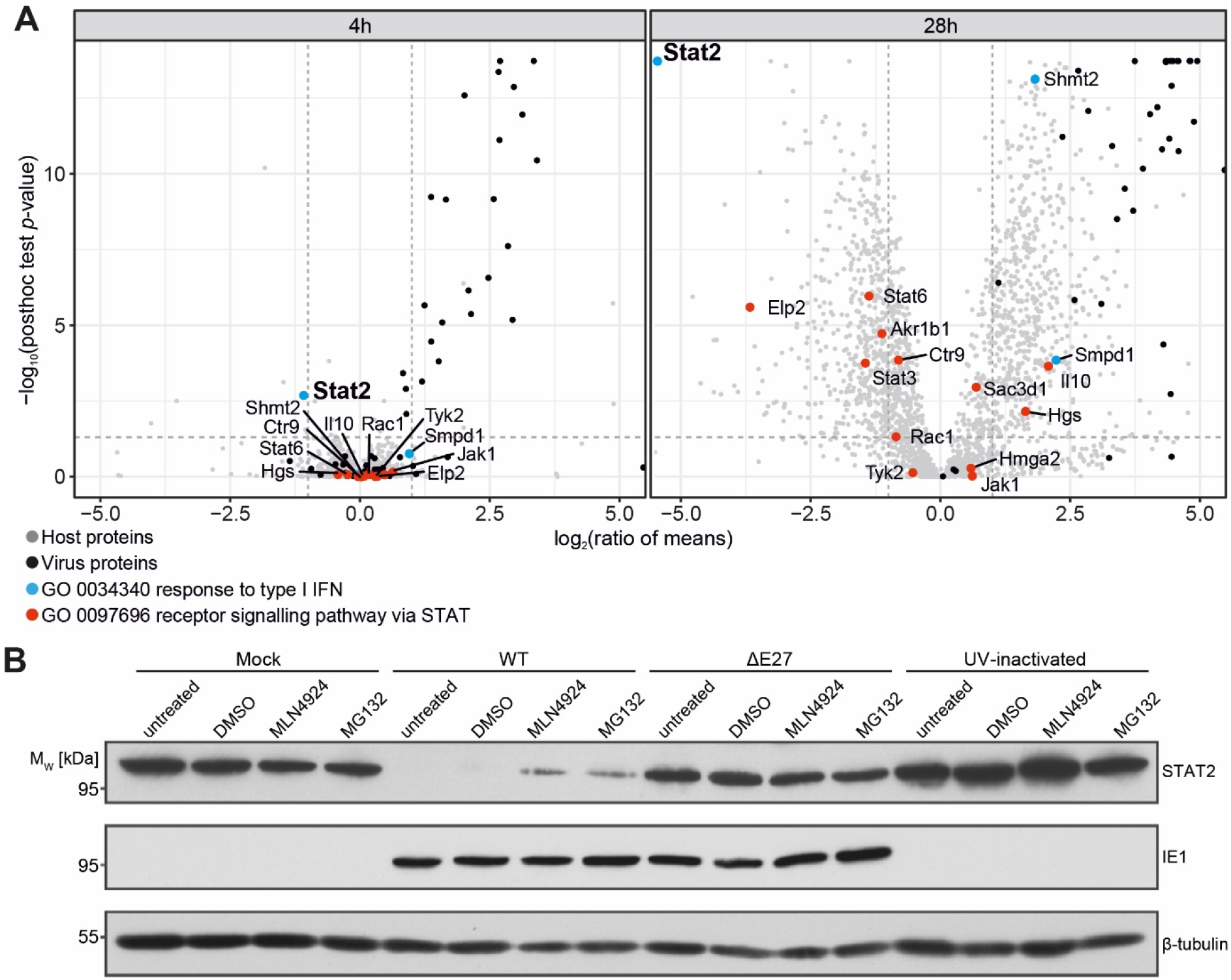
Host protein regulation during RCMV-E infection and the role of E27. **(A)** Mass spectrometric analyses of protein abundance changes upon RCMV-E infection, compared to mock-infected samples. Host and virus proteins are indicated as grey and black dots, respectively. Host proteins involved in the type I IFN response and the STAT signalling pathway are highlighted in blue and red. **(B)** Western blot analysis of STAT2 protein levels in mock-, WT-RCMV-E-, ΔE27-RCMV-E-or UV-inactivated RCMV-E-infected REF. 24 h p.i., cells were either left untreated or treated for 5.5 h with DMSO, 2.5 µM MLN4924 for blocking CRL activity or 10 µM MG132 for proteasome inhibition. IE1 and *β*-tubulin served as infection and loading controls.

Next, we sought to identify the RCMV-E-encoded factor responsible for the reduction of STAT2 levels. Inspection of the RCMV-E genome (Geyer et al., 2015) revealed 56% sequence identity of E27 to M27, the factor inducing STAT2 degradation in MCMV infection. To test if E27 accounts for STAT2 down-regulation, an RCMV-E mutant lacking *E27* was constructed by CRISPR/Cas9-mediated mutagenesis resulting in an E27-deficient RCMV-E (“ΔE27”). REF were infected with wild type (“WT”) and ΔE27 virus and STAT2 expression was assessed by immunoblot analysis 24 h post infection (p.i.). While WT infection confirmed the MS analysis and caused a complete loss of the STAT2 signal, the ΔE27 virus failed to reduce STAT2 protein concentrations (Fig. 1B). Moreover, the WT-mediated loss of STAT2 proved to be sensitive to treatment with MG132, a proteasome inhibitor, and MLN4924 (Fig. 1B), an inhibitor of the NEDD8-activating enzyme, which shuts off cellular neddylation processes that are essential for CRL activity. These findings strongly suggested an involvement of the ubiquitin proteasome system in E27-induced STAT2 down-regulation and furthermore indicated the involvement of host CRLs.

### E27 recruits STAT2 to CRL4 in vitro

These results prompted us to explore the physical association of E27 with STAT2 and DDB1. Most viral STAT2 antagonists are known to be species-specific (Ashour et al., 2010; Parisien et al., 2002a; Yoshikawa et al., 2019). Therefore, we cloned and purified *Rattus norvegicus* (rn) STAT2 for these studies. Conversely, hsDDB1 and rnDDB1 are almost identical (1128 of 1140 amino acids are conserved, and the few changes are located in the β-propeller domain B [BPB] of DDB1 or in disordered loops and regions not involved in intermolecular interactions). Accordingly, we applied hsDDB1, in analogy to various studies by us and others concerning DDB1 binders (Angers et al., 2006; Banchenko et al., 2021; Fischer et al., 2014; Fischer et al., 2011; Li et al., 2006). The incubation of DDB1 with equimolar amounts of STAT2, followed by analytical gel filtration (GF) chromatography, resulted in elution in separate peaks at the same elution volumes as the individual proteins in isolation (Fig. 2A, blue, magenta, and grey traces). This argues against an intrinsic association of DDB1 and STAT2. In contrast, E27 formed stable binary complexes with STAT2 (Fig. 2B, orange trace) and DDB1 (brown trace). Incubation of E27 together with DDB1 and STAT2 resulted in the elution of all three components in the same peak at an earlier elution volume than the isolated components (Fig. 2A, purple trace). This showed the formation of a stable ternary DDB1/E27/STAT2 protein complex, demonstrating that E27 is sufficient to bridge STAT2 to DDB1.

**Fig. 2:**
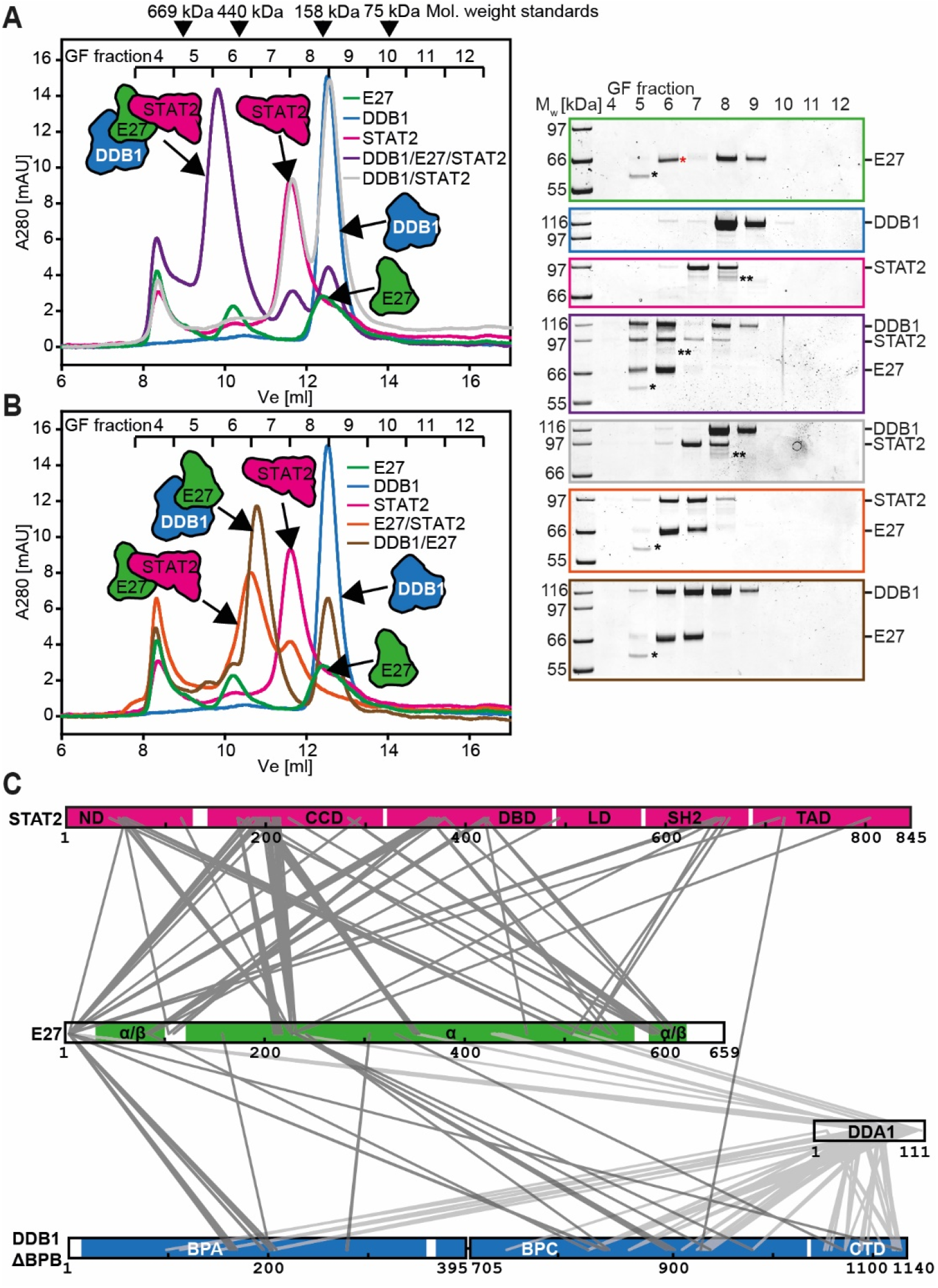
*In vitro* reconstitution and XL-MS analysis of DDB1(/DDA1)/E27/STAT2 protein complexes. (**A, B**) GF analysis of *in vitro* reconstitutions containing the indicated protein combinations. Coomassie blue-stained SDS-PAGE analyses of fractions collected during the GF runs are shown next to the chromatograms. The red * indicates a contaminant, which we identified by cryo-EM analysis as *E. coli* ArnA, a notorious contaminant in *E. coli* Ni-NTA protein purifications (Andersen et al., 2013) (Fig. S2, S3). Note that ArnA is present in GF fraction 6 (V_e_ ∼ 10.2 ml) of all runs containing E27, and migrates similarly to E27 on the SDS-PAGE. * - additional contaminant from E27 preparation; ** - contaminant from STAT2 preparation. **(C)** Schematic of proteins used for XL-MS analysis with domains annotated. Heteromeric cross-links obtained from XL-MS involving DDA1 are shown as light grey lines, otherwise as dark grey lines. ND – N-terminal domain, CCD – coiled-coil domain, DBD – DNA-binding domain, LD – linker domain, SH2 – Src homology 2, TAD – transactivation domain, BP – β-propeller, CTD – C-terminal domain.

To obtain complementary information on the topology of the DDB1/E27/STAT2 complex, and to determine a possible involvement of the CRL4 co-adapter DDA1 in E27 and/or STAT2-binding, cross-linking mass spectrometry (XL-MS) was performed. DDB1 lacking its β-propeller (BP) domain B (DDB1_ΔBPB_) was assembled with hsDDA1 (98% identical to rnDDA1), E27, and STAT2 *in vitro* and purified by GF. Subsequently, the complex was cross-linked using the UV-activated cross-linker sulfo-SDA, followed by trypsin digestion and analysis of cross-linked peptides by XL-MS. Twenty-eight and 96 heteromeric cross-links were identified between DDB1_ΔBPB_ and E27, and between E27 and STAT2, respectively, while only 1 cross-link between DDB1_ΔBPB_ and STAT2 was found (Fig. 2C). These data are in line with a model where E27 serves as connector between DDB1 and STAT2, and indicate that the DDB1 BPB is dispensable for the DDB1/E27/STAT2 complex formation. Furthermore, 69 and 21 cross-links were found between DDA1 and DDB1_ΔBPB_ or E27, respectively, but no cross-links extended from DDA1 to STAT2 (Fig. 2C). These observations argue for a role of DDA1 in the stabilisation of the DDB1/E27-interaction, but against its involvement in the STAT2 recruitment.

### *Cryo-EM structure of the DDB1* _ΔBPB_*/DDA1/E27/STAT2 complex*

Cryo-EM imaging of a cross-linked DDB1_ΔBPB_/DDA1/E27/STAT2 complex (Fig. S2) was pursued to gain structural insights. Single particle analyses yielded a density map at 3.8 Å resolution (map 1) (Table S1, Fig. S3). No experimental cryo-EM density corresponding to the expected position of DDA1 was observed (Fig. S4A). Based upon distance restraints obtained from XL-MS (Fig. 2C), the space accessible for interaction with the centre of mass of the DDA1 C-terminal helix was calculated using the DisVis algorithm (van Zundert and Bonvin, 2015). Visualisation of this space indicates that DDA1 preferably locates to a large, ill-defined DDB1 surface region around the edges of BPC and CTD. These data indicate that DDA1 is flexible and does not stably interact with E27, accounting for the lack of cryo-EM density (Fig. S4B). Conversely, clear cryo-EM density corresponding to DDB1_ΔBPB_ was present, allowing to unambiguously position both BP domains of the truncated DDB1 construct (Fig. 3A, S5A). Furthermore, assisted by XL-MS distance restraints and *in silico* structure prediction using AlphaFold2 (Jumper et al., 2021), a molecular model corresponding to E27 amino acid residues 105-570, folding into an elongated bundle containing 18 α-helices (the E27 α-domain), could be fitted into a density segment on top of DDB1_ΔBPB_ (Fig. 3A, S5B, see Methods for details). Prominent structural features were a protrusion from the edge of the α-domain, mainly formed by helix α8, inserting into a binding cleft formed by the DDB1 BPs, and a putative metal-binding motif composed of three cysteines and one histidine at the base of the protrusion. Taking into account the wealth of previous structural information (Ireland and Martin, 2019; Laitaoja et al., 2013), the tetragonal coordination geometry, the nature of the amino acid side chains coordinating the metal, as well as the coordination bond lengths strongly argue in favour of a structural zinc-binding site. Accordingly, we modelled a zinc ion in the corresponding position (Fig. S5C). Moreover, cryo-EM map 1 showed extra density features consistent with the presence of a 4-helix bundle on top of E27, opposite of the DDB1-binding interface (Fig. 3A). With the aid of an AlphaFold2 structure prediction of rnSTAT2, this density segment could be interpreted as a major portion of the STAT2 coiled-coil domain (CCD) encompassing residues 144-180 from helix αA, 196-231 (αB), 259-288 (αC) and 293-317 (αD) (Fig. S5D). 21 heteromeric cross-links identified by XL-MS could be mapped on the DDB1_ΔBPB_/E27/STAT2 CCD structure. In support of the cryo-EM model, 16 of these cross-links are within the 25 Å distance threshold enforced by the cross-linker chemistry, and the remainder are only slight outliers (distances between 25 Å to 35 Å) (Fig. S5E, S6, Table S2).

**Figure 3:**
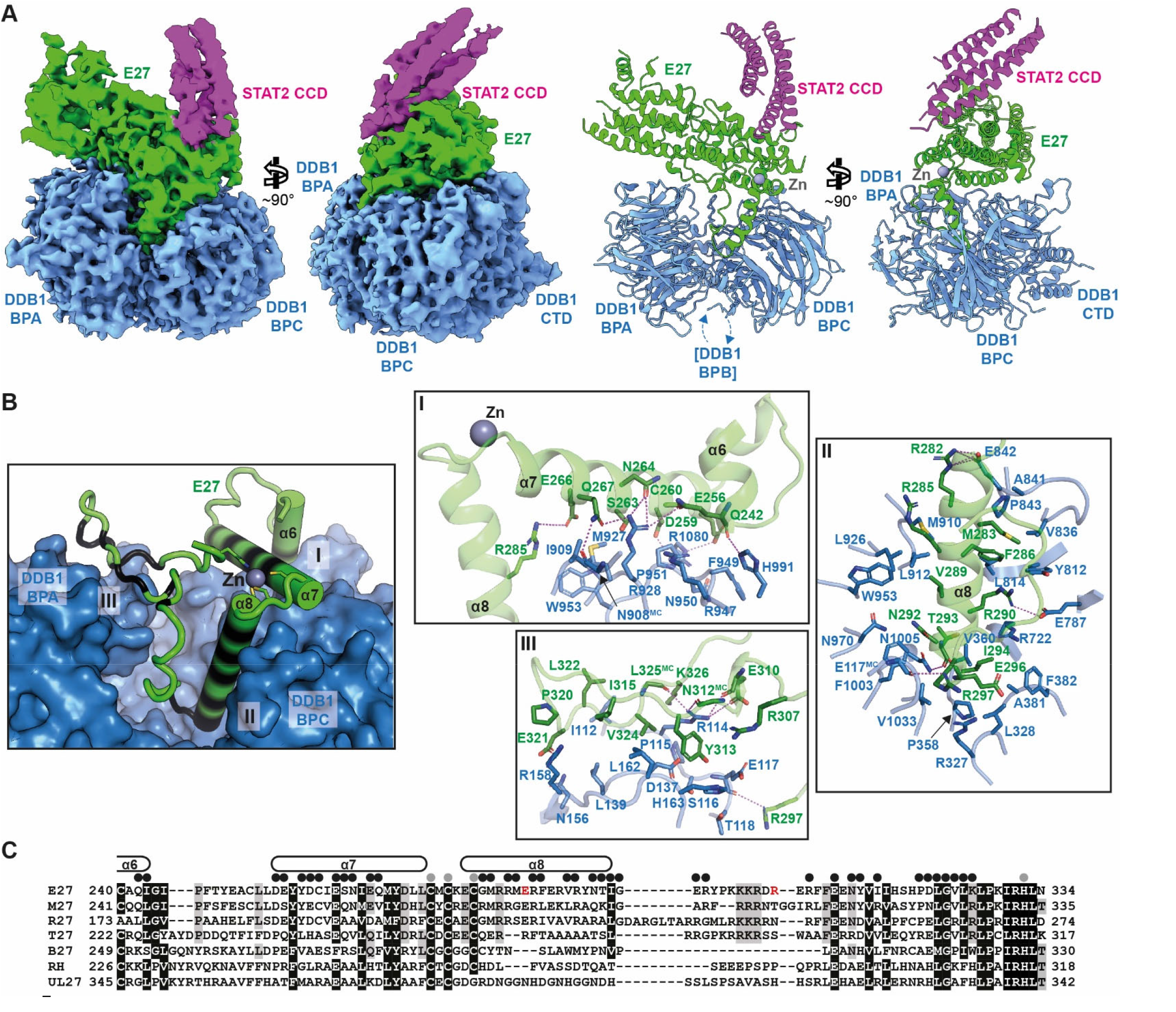
Cryo-EM structure of the DDB1ΔBPB/E27/STAT2 CCD protein complex. **(A) (Left panel)** Two views of cryo-EM map 1 contoured at threshold level 0.35 by the Chimerax software tool (Pettersen et al., 2021). Density segments corresponding to DDB1_ΔBPB_ are coloured blue, to E27 green and to the STAT2 CCD magenta. **(Right panel)** Cartoon representation of the molecular model fitted in cryo-EM map 1, in the same orientation as in the left panel. DDB1_ΔBPB_ is coloured blue and E27 green, the zinc ion is shown as grey sphere. **(B) (Left panel)** Overview of the DDB1_ΔBPB_/E27 interaction interface. DDB1_ΔBPB_ is shown in solvent-accessible surface representation and coloured blue, E27 is shown as green cartoon with helices represented as cylinders. E27 residues in contact with DDB1_ΔBPB_ are shaded black. The zinc ion is shown as black sphere and E27 side chains involved in zinc coordination are shown as sticks. **(Right panels)** Detailed views of interaction regions of interest I-III indicated in the left panel. Selected amino acid residues involved in intermolecular contacts are shown as sticks. Dashed lines represent hydrogen bonds or salt bridges. **(C)** Amino acid sequence alignment of E27 and homologues from other cytomegaloviruses. E27 residues contacting DDB1 are indicated by black dots and residues involved in zinc coordination as grey dots. E27 secondary structure elements are drawn above the alignment.

### *The DDB1*_ΔBPB_*/E27 protein complex*

Initially, we focused the structure analysis on the DDB1_ΔBPB_/E27 interaction. E27 binds to a cleft lined by the inner surfaces of DDB1 BPA and BPC (Fig. 3B). In this way, it is positioned on the opposite side of where DDB1 BPB would be located, which links to the CRL4 core. E27 helices α6, α7, α8, and the loop leading to α9 are in direct contact with DDB1 and helix α8 protrudes in the binding cleft (Fig. 3B). The DDB1/E27 binding interface, covering an area of 2,131 Å^2^, is divided in three regions (Fig. 3B, S7A). The first involves E27 helices α6 and α7, bordered by the zinc-coordinating cysteine triad, contacting residues located in DDB1 BPC WD40 repeats 6 and 7, and in the CTD. It is formed by a network of hydrogen bonds, additionally supported by salt bridges and hydrophobic interactions (Fig. 3B, box I). A second, central interaction region includes the protrusion formed by helix α8 and a subsequent polybasic loop, contacting residues lining the inner surface of DDB1 BPC and BPA WD40 repeats 3 and 7. This interface is mostly hydrophobic, with fewer polar and electrostatic interactions (Fig. 3B, box II). A third region comprises the extended loop leading back to the E27 zinc-binding motif, adjacent to DDB1 BPA WD40 repeats 3 and 4. Again, mainly hydrophobic interactions form this interface, assisted by hydrogen bonding (Fig. 3B, box III).

To evaluate the conservation of the DDB1/E27-binding surface, we mapped interface residues on a multiple sequence alignment of E27 homologues (Fig. 3C). Twenty-eight out of 38 interfacing residues are type-conserved between RCMV-E E27 and MCMV M27, while 26 are type-conserved between E27 and R27, the homologue encoded by RCMV-Maastricht (RCMV-M). Furthermore, 16 and 12 interface residues are type-conserved in E27 homologues from bat and tupaiid CMV, respectively, and only 8 and 9 residues in rhesus macaque CMV and HCMV, respectively. Additionally, in the case of tupaiid, bat, rhesus macaque, and human CMV, high sequence divergence of the central α8 binding region is apparent. Together, this suggests that DDB1-binding functionality is restricted to rodent CMV E27, M27, and R27.

To validate the structure, an E27 construct was prepared, from which a significant part of the DDB1-binding region (residues E284-R305, marked red in Fig. 3C) was deleted and replaced by a flexible linker (E27Δ). E27Δ was purified and employed for *in vitro* reconstitution with DDB1 or STAT2. Subsequent gel filtration analysis showed that E27Δ, in contrast to WT E27, was unable to bind DDB1 (Fig. S7B). However, like WT E27, E27Δ bound STAT2, demonstrating that the overall integrity of E27Δ was not compromised by mutagenesis (Fig. S7C). This suggests that the central binding region (Fig. 3B, box II) is critical for E27/DDB1 interaction, while the peripheral regions (Fig. 3B, boxes I and III) are not sufficient to sustain stable E27/DDB1 association on their own.

### E27 mimics endogenous CRL4 substrate receptors

Next, the DDB1_ΔBPB_/E27 complex structure was compared to the DDB1-binding modes of DCAF1 and of the previously characterised viral DCAFs (vDCAFs) SV5 V and HBx. Overall, they are positioned similarly relative to DDB1 (Fig. 4A). Analysis of their “molecular footprint” on DDB1 reveals that the E27 footprint is almost identical to the DCAF1-derived one, while SV5 V and HBx bind DDB1 in significantly smaller interaction interfaces, mostly restricted to DDB1 BPC (Fig. 4B). To engage DDB1 surface regions, E27 utilises other structural elements than DCAF1, i.e. a short and two long helices (α6-α8) and an extended loop, in contrast to three short helices and the bottom side of four WD40 repeats in DCAF1 (Fig. 4A, insets). Superposition of DDB1-bound E27 and cellular and viral DCAF structures showed that the only common secondary structure element is the C-terminal half of E27 α8, which aligns well with a short helical “H-box” motif in DCAF-type receptors (Fig. 4C). However, sequence conservation of this motif is low, with only four amino acid positions >60% type-conserved (Fig. 4D). Together, these data demonstrate how E27 emulates the DDB1-binding mode of endogenous DCAF substrate receptors using divergent structural principles. Furthermore, the analysis illustrates that all examined vDCAFs engage DDB1 via additional structural elements beyond the previously described “H-box” motif (Li et al., 2010). However, their structure and sequence are not conserved, and the only common characteristics are contact areas on the upper edge of DDB1 BPC. Accordingly, structural analyses of vDCAF/DDB1 interactions are necessary for a full mechanistic understanding of how they hijack CRL4.

**Figure 4:**
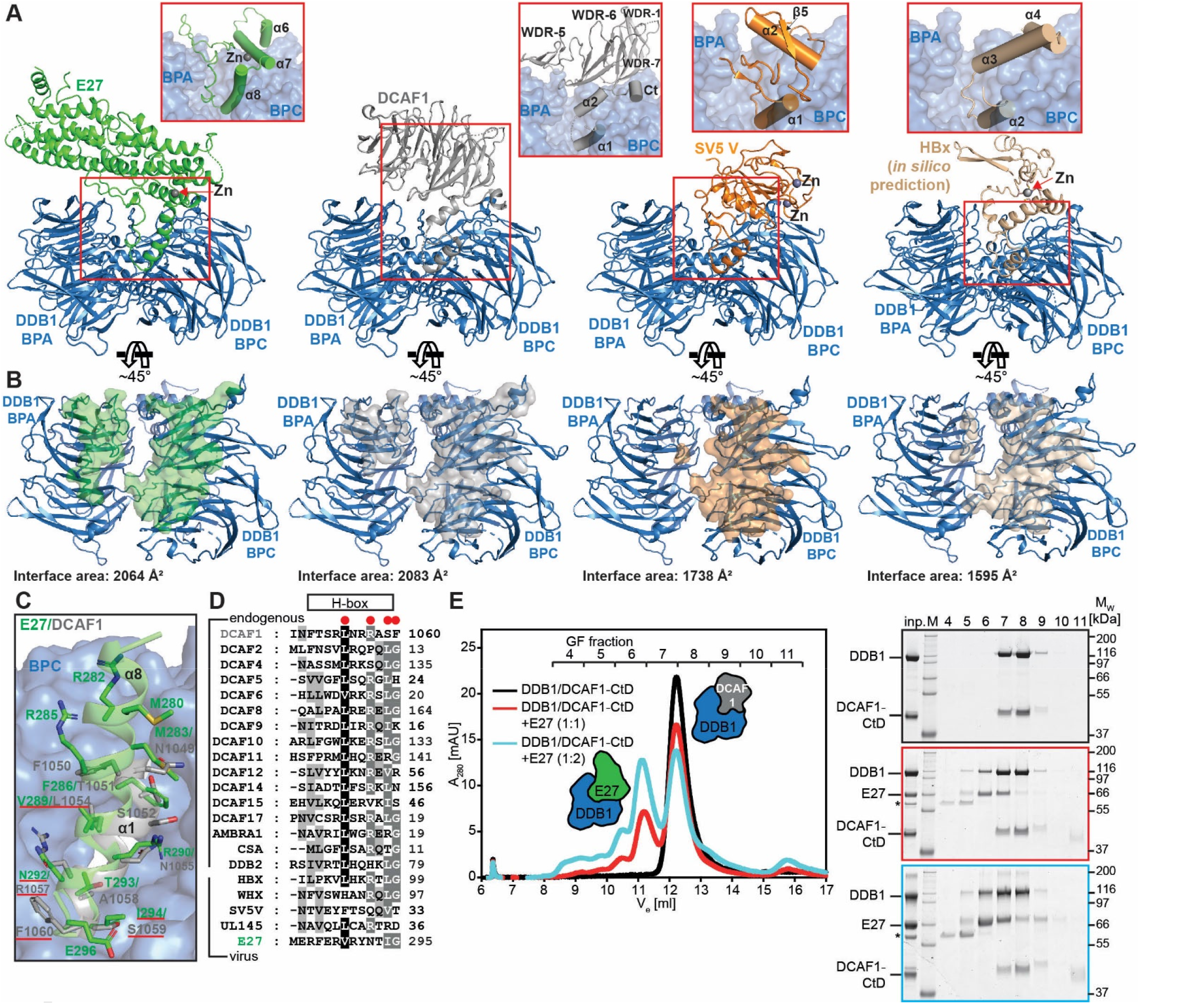
E27 replaces endogenous CRL4 substrate receptors. **(A)** Comparison of the DDB1/E27-binding mode with endogenous and viral CRL4 substrate receptors. Structures of DDB1 (blue) in complex with E27 (green), DCAF1 (grey, PDB 6zue (Banchenko et al., 2021)), SV5 V (orange, PDB 2hye (Angers et al., 2006)), and HBx (wheat, AlphaFold2 prediction, Fig. S8) are shown as cartoons. Regions of the AlphaFold2 HBx model with very low confidence have been removed. The insets display enlarged views of the boxed interaction areas, with DDB1 shown as semi-transparent blue surface. Zinc ions are shown as grey spheres. The position of the HBx zinc ion was modelled based on information from (Ramakrishnan et al., 2019). **(B)** “Molecular footprint” of substrate receptors from **A** on the DDB1 surface. DDB1 is shown in the same representation as in **A**, but slightly rotated to allow an unobstructed view of the receptor-binding cleft. The receptors themselves have been removed for clarity. Parts of the DDB1 surface in contact with the respective substrate receptor are coloured according to the scheme in **A**. **(C)** Detailed comparison of a helical DDB1-binding motif in E27 (green) and the DCAF1 “H-box” (grey). DDB1 is shown as semi-transparent blue surface, E27 and DCAF1 as semi-transparent cartoon. Selected amino acid residues involved in intermolecular contacts are shown as sticks. **(D)** Amino acid sequence alignment of the “H-box” DDB1-binding motif in endogenous substrate receptors and viral factors hijacking DDB1. Black boxes: >80% type-conserved; dark grey: >60%; light grey: >40%. >60% type-conserved residues are marked by red dots, and underlined in **C**. **(E)** Pre-formed DDB1/DCAF1 protein complex was incubated with indicated amounts of E27 and analysed by GF. SDS-PAGE analyses of GF fractions are shown next to the chromatogram. * - contaminant from E27 preparation.

Given the similar position and interface area of E27 and DCAF1 on DDB1, we hypothesised that E27 competes with endogenous DDB1/DCAF complexes for DDB1-binding. To gain biochemical insight, the pre-formed DDB1/DCAF1 complex was incubated with E27 and subjected to analytical GF. These analyses showed the formation of an elution peak containing DDB1 and E27, concomitant with reduction of the DDB1/DCAF1 complex peak at later elution volume, in an E27 concentration-dependent manner, indicating that E27 is able to displace DCAF1 from DDB1 (Fig. 4E).

### Mechanism of STAT2 recruitment

Based on cryo-EM map 1, the binding between E27 and the STAT2 CCD was analysed in detail (Fig. 5A, B, S9A). The interaction interface covers a surface area of 750 Å^2^. It comprises hydrophobic residues from STAT2 CCD helices αA, αB, and the C-terminal loop of αC. These residues project into a hydrophobic pocket lined by E27 amino acids from the loop downstream of α4 and side chains from a sequence stretch spanning α11 to α12. In addition, several peripheral hydrogen bonds and a salt-bridge network strengthen the interface.

**Figure 5:**
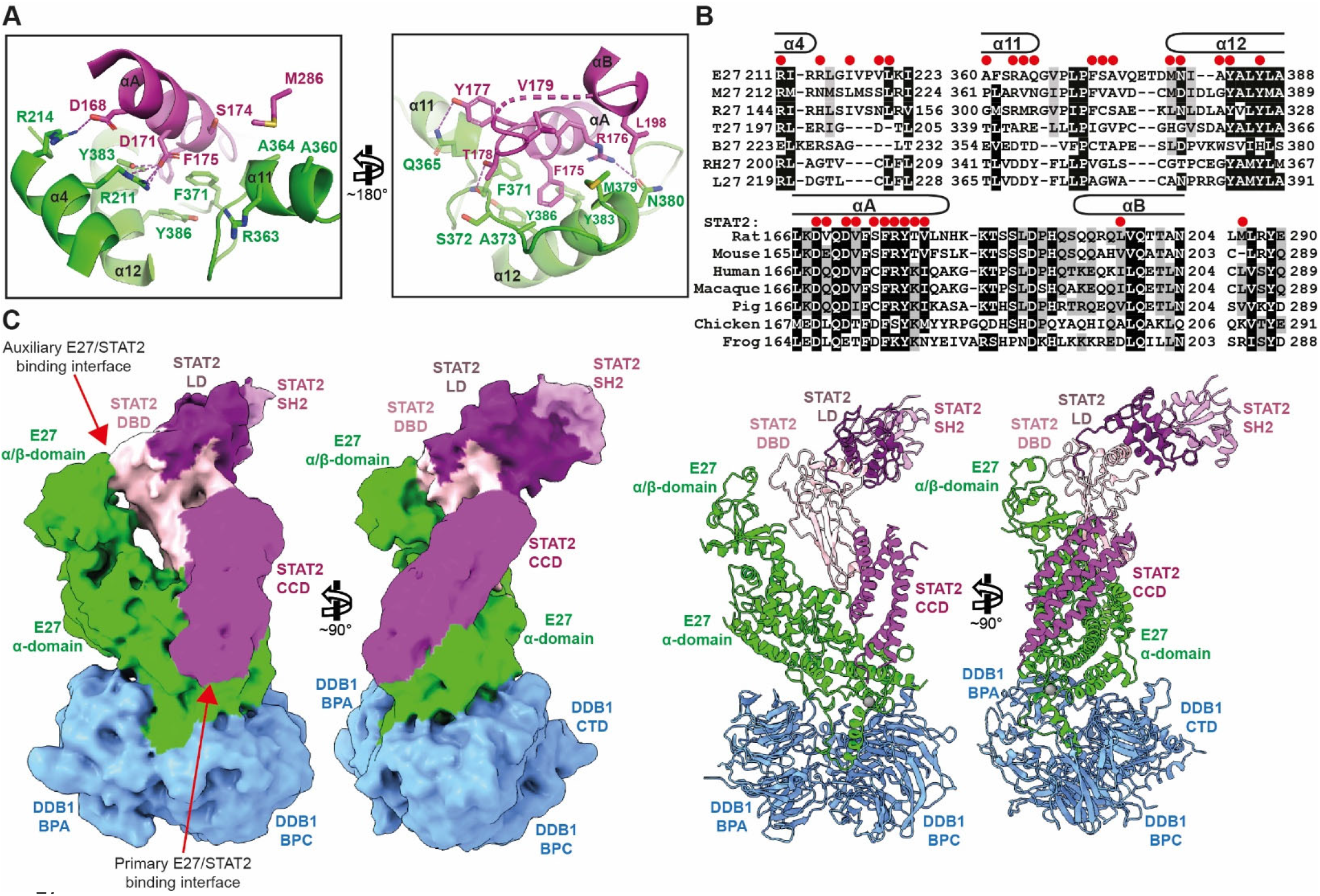
Mechanism of STAT2 recruitment. **(A)** Detailed view of the E27/STAT2 CCD interaction, derived from the cryo-EM map 1 molecular model. Proteins are represented as cartoons; E27 is coloured green and STAT2 magenta. Selected side chains involved in intermolecular interaction are shown as sticks. Dashed lines represent hydrogen bonds or salt bridges. **(B) (Upper panel)** Amino acid sequence alignment of E27 and homologues from other cytomegaloviruses. E27 residues contacting STAT2 are indicated by red dots. E27 secondary structure elements are drawn above the alignment. **(Lower panel)** Amino acid sequence alignment of STAT2 and homologues from other species. STAT2 residues contacting E27 are indicated by red dots. STAT2 secondary structure elements are drawn above the alignment. **(C) (Left panel)** Two views of cryo-EM map 2 contoured at threshold level 0.15 by the Chimerax software tool (Pettersen et al., 2021). Density segments corresponding to DDB1_ΔBPB_ are coloured blue, to E27 green, to the STAT2 CCD magenta, to the STAT2 DBD light pink, to the STAT2 LD purple, and to the STAT2 SH2 domain dark pink. **(Right panel)** Cartoon representation of the molecular model fitted in cryo-EM map 2, in the same orientation and colour code as in the left panel. The zinc ion shown as grey sphere.

Conservation analysis of this binding interface revealed that out of 17 involved residues, 12 and 11 are type-conserved in rodent E27 homologues (M27 and R27, respectively), while only 8 are type-conserved in their tupaiid and bat counterparts, and only 6 in macaque or human CMV (Fig. 5B). Additionally, in the more distantly related, non-rodent CMV species, most of the VPVL loop downstream of α4 is deleted, removing a significant portion of the STAT2 CCD-binding site. Thus, it is likely that STAT2 recruitment is restricted to rodent E27 homologues. Variation within this branch might be driven by species-specific adaptation to differences in the respective host STAT2 sequences, e.g. rnSTAT2 positions 169, 198, and 285 (Fig. 5B).

To corroborate the structural data, an E27 R211E/R214E double mutant (E27-RE) was generated by site-directed mutagenesis to disrupt the intricate salt-bridge network between these residues and STAT2 D168 and D171 (Fig. 5A). The E27-RE variant, alongside WT E27, was subjected to *in vitro* reconstitution with STAT2 or DDB1, followed by gel filtration analysis. The results demonstrate that E27-RE had lost the ability to form a stable complex with STAT2 (Fig. S9B), while it retains DDB1-binding activity (Fig. S9C), thus validating the molecular model of STAT2 CCD recruitment.

Moreover, further 3D classification of the cryo-EM particle images yielded a second map at 5 Å resolution, exhibiting additional density features in the E27 and STAT2 regions (map 2, Fig. 5C, S3, Table S1). This allowed the placement of a STAT2 AlphaFold2 model lacking the N-terminal and TAD domains, and enabled fitting of the terminal E27 α/β-domain, revealing that at least a subset of the particles observed in the cryo-EM analysis exhibit a bipartite E27/STAT2-binding interface (Fig. 5C). It comprises the primary interaction between the STAT2 coiled-coil domain (CCD) and the E27 α-domain, which has also been observed in cryo-EM map 1 (Fig. 3, 5A), as well as a second, auxiliary contact region involving the STAT2 DNA-binding domain (DBD) and the E27 α/β-domain (Fig. 5C, S10A, B). Thirty-nine out of 96 heteromeric cross-links could be mapped on this E27/STAT2 complex, of which 22 lie within the 25 Å distance threshold and 9 are slight outliers (25 Å - 35 Å), supporting the structural data. Four severely overlength cross-links are apparent (>40 Å), all of which involve the E27 α/β-domain or the STAT2 SH2 domain, indicating flexibility of these distal components (Fig. S10C), rationalising the low local resolution of the corresponding cryo-EM map regions (Fig. S3).

The auxiliary E27/STAT2 interaction area involves parts of the E27 α/β-domain, located between strand βA and helix αA, between αB and βC, and a C-terminal stretch upstream of βD (Fig. S10A). These regions form a shallow surface which contacts STAT2 DBD protrusions comprising amino acid stretches 377-383 and 406-416. However, due to the low local resolution of the cryo-EM map, we refrained from modelling individual side chain interactions.

### E27 positions STAT2 appropriately for CRL4-catalysed ubiquitin transfer

The structure of the DDB1_ΔBPB_/E27/STAT2 complex enabled modelling of STAT2 in the context of a complete CRL4 assembly (Fig. 6). The model shows that STAT2 is placed centrally in the ubiquitylation zone created by rotation of the CUL4/RBX1 scaffold arm. This rotation is enabled by a flexible hinge connecting DDB1 BPB, where CUL4/RBX1 is attached, to DDB1 BPC, which together with the BPA forms the binding platform for substrate receptors. Furthermore, the model also provides a rationale for the existence of the second, auxiliary E27/STAT2 DBD-binding interface, which nudges the STAT2 molecule into the ubiquitylation zone. In such a way, surface-exposed lysines on all STAT2 domains are within reach of ubiquitin-charged E2, which is positioned for ubiquitin transfer by a catalytic assembly comprising the neddylated distal end of CUL4 and RBX1 (Baek et al., 2020), guaranteeing efficient STAT2 ubiquitylation and subsequent degradation.

**Figure 6:**
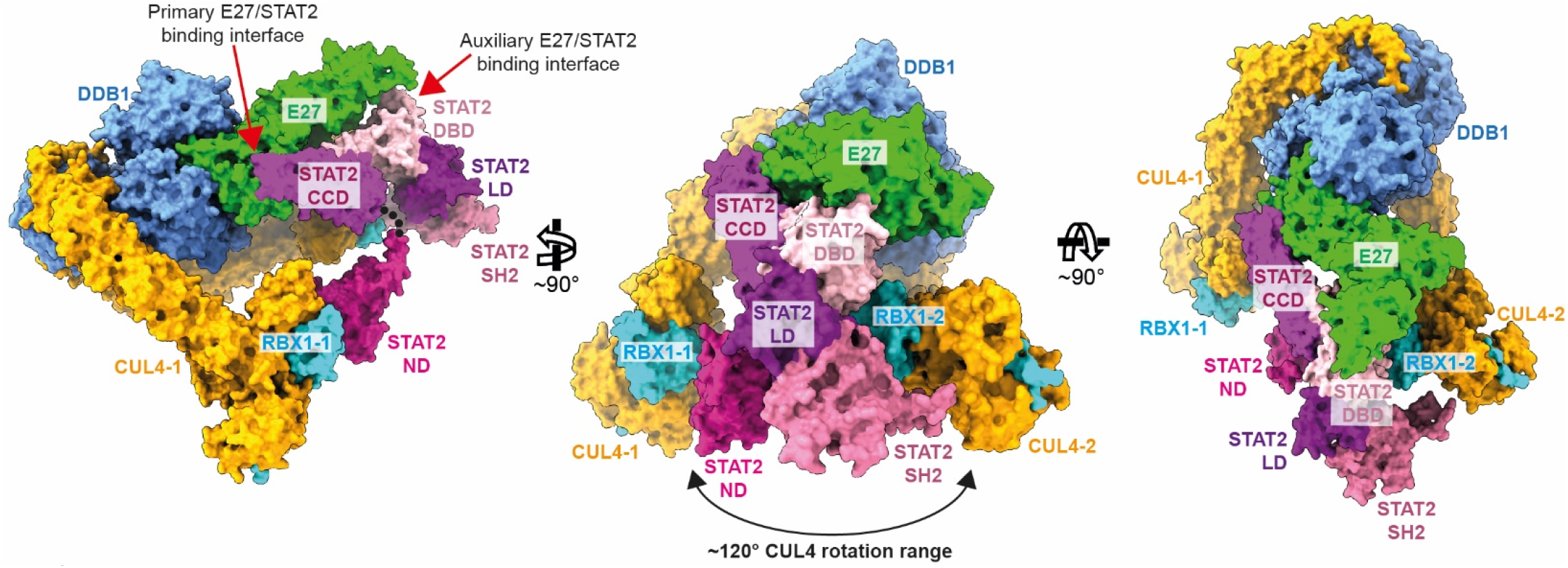
Model of the complete CRL4^E27^/STAT2 assembly. A model of how E27/STAT2 is positioned relative to the remainder of CRL4 was assembled by superposition of DDB1_ΔBPB_/E27/STAT2 with DDB1 BPA/BPC from two CRL4^DCAF1^ cryo-EM structures, which represent the two outermost positions of the CUL4 scaffold (CUL4-1, CUL4-2), thus visualising the rotation range of CUL4/RBX1 with respect to DDB1 (Banchenko et al., 2021). RBX1 (cyan), CUL4 (orange), DDB1 (blue), E27 (green) and STAT2 (coloured as in Fig. 5) are shown in surface representation. The STAT2 N-terminal domain (magenta cartoon) was modelled in an arbitrary position, since it is connected to the STAT2 CCD via a flexible linker (indicated by the dotted line).

### The E27 structure reveals a conserved zinc-binding motif of functional importance

The cryo-EM density (map 1) revealed a cryptic zinc-binding motif in E27 involving three cysteines (C273, C275, C278) and H332 from a loop upstream of helix α9 (Fig. 7A, S5C), anchoring the protrusion, which inserts into the binding cleft on DDB1, to the E27 α-domain bundle. The sequence conservation of this motif argues strongly in favour of its functional relevance (Fig. 7B), which prompted us to investigate its function in closely related cytomegaloviruses. MCMV M27, the closest E27 homologue, exhibits 56% and 76% amino acid sequence identity and similarity, respectively (Fig. S10A). E27 and M27 AlphaFold2 structure predictions are almost indistinguishable (RMSD of 1.195 Å, 6,652 aligned atoms; Fig. S11), and structural alignment of these predictions demonstrated conservation of side chain positions involved in zinc-binding (Fig. 7A). Thus, M27 represents a valid model to study functional consequences of interference with the zinc coordination motif through site-directed mutagenesis, and to test the relevance of our E27 structure model for the MCMV system.

**Figure 7:**
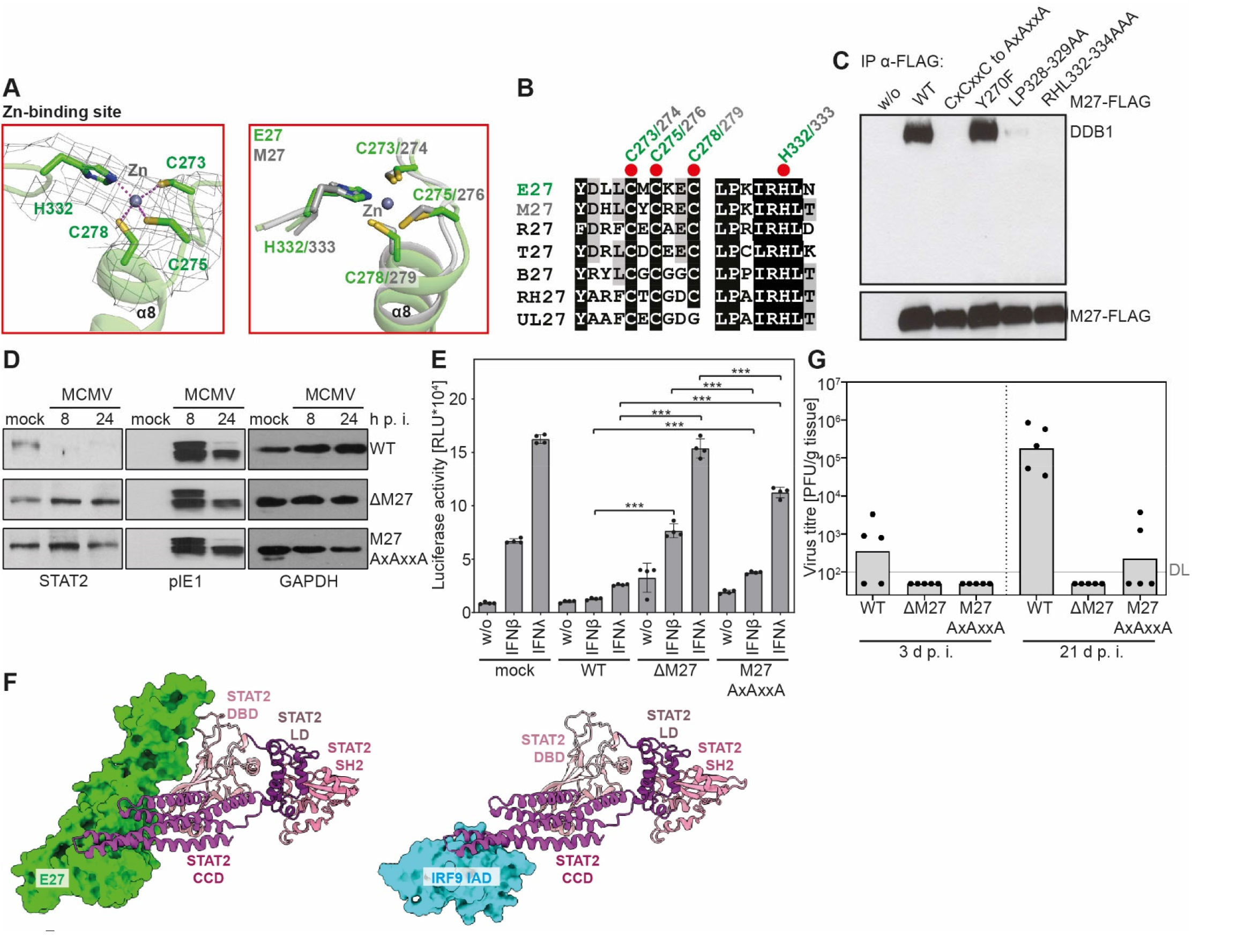
E27 structure reveals a conserved zinc-binding motif of functional importance *in vivo*. **(A)** Detailed view of the zinc-binding motif. E27 is shown as green cartoon, residues involved in zinc-binding as sticks. Zinc coordination bonds are indicated as dashed lines. Cryo-EM density, contoured at 6s, is shown as grey mesh. The right panel shows a superposition of the E27 (green) and M27 (grey) zinc-binding sites. **(B)** Amino acid sequence alignment of E27 and homologues from other CMV species. Residues involved in zinc coordination are marked by red dots. **(C)** 293T cells were transfected with M27-FLAG or the indicated M27-FLAG variants. 24 h after transfection, cells were lysed and M27-FLAG was precipitated using a FLAG-specific antibody. Precipitates were separated by SDS-PAGE, blotted and probed using the indicated antibodies. **(D)** Mouse fibroblasts were infected with MCMV or the indicated MCMV mutants. At the indicated time points, cells were lysed, proteins were separated by SDS-PAGE and blotted. Blots were probed using the indicated antibodies. pIE1 and GAPDH were used as infection and loading controls. **(E)** NIH3T3 reporter cells stably expressing ISRE-Luc were infected with the indicated MCMV strains, treated with IFNβ or IFNλ, and luciferase activity was measured. *** – P ≤ 0.001 (ANOVA). **(F)** Comparison of STAT2-binding to E27 and IRF9 (PDB 5oen) (Rengachari et al., 2018). STAT2 is shown as cartoon and coloured as in Fig. 5. E27 (green) and the IRF9 IRF-associated domain (IAD, cyan) are shown in surface representation. **(G)** BALB/c mice were infected i.p. with WT-MCMV and indicated MCMV mutants. At 3 and 21 d p.i., salivary glands were collected and frozen. Virus titres were determined from organ homogenates by plaque titration. Titrations were performed in quadruplicates. Bars depict the geometric mean, dots show titres of individual mice. DL – detection limit.

First, in FLAG-tagged M27 constructs, individual residues and combinations in the zinc-binding site were mutated to alanine. The constructs were expressed in 293T cells and their ability to bind endogenous DDB1 was assessed by co-immunoprecipitation. While WT-M27 efficiently co-precipitated DDB1, mutation of the three zinc-coordinating cysteine residues abrogated DDB1-binding (Fig. 7C). Likewise, a mutation of the loop containing the zinc ligand H333 and of an adjacent LP motif interfered with DDB1-interaction. Serving as an additional control, the Y270F mutation upstream of the cysteine triad had no effect on DDB1-binding. Thus, the integrity of the zinc-binding motif is necessary for interaction with DDB1.

Next, MCMV mutants lacking M27 (“ΔM27”) or carrying the AxAxxA triple cysteine mutation (M27_AxAxxA_) were generated. Immortalised mouse embryonic fibroblasts were infected with these viruses alongside WT-MCMV as a control. STAT2 protein levels of cell lysates were probed by western blot. As expected, the WT-MCMV infection depleted STAT2. In contrast, the M27_AxAxxA_-MCMV failed to reduce STAT2 protein levels, phenocopying the ΔM27 virus (Fig. 7D). In addition, to test the impact of these MCMV variants on ISG induction, NIH3T3 cells stably expressing an interferon-stimulated response element (ISRE)-dependent luciferase reporter were infected with WT, ΔM27, and M27_AxAxxA_ viruses. Cells were treated with IFN, and luciferase activity was quantified as reporter for IFN-driven gene expression. In accordance with the STAT2 degradation, IFN signalling was efficiently supressed following WT-MCMV infection. Conversely, ΔM27-MCMV failed to counteract IFN-induced ISRE signalling. Intriguingly, the M27_AxAxxA_-MCMV infection exhibited strongly attenuated but incomplete IFN inhibition (Fig. 7E). These data demonstrate that the disruption of the M27 zinc-binding site interferes with M27-dependent STAT2 degradation, and alleviates but not fully abrogates suppression of IFN responses.

The residual IFN inhibitory capacity of M27_AxAxxA_-MCMV prompted us to find a molecular explanation. A comparison of the E27/STAT2 CCD binding mode discovered here to the IRF9/STAT2 CCD complex (Rengachari et al., 2018) demonstrates overlap of the binding interfaces (Fig. 7F). Consequently, the STAT2/E27 interaction is sterically incompatible with IRF9-binding, prohibiting formation of an active ISGF3 signalling complex. This might provide a second layer of STAT2 inhibition, beyond induction of its proteasomal degradation, and might rationalise the observed significant residual ISG-suppression in cells infected with the DDB1-binding-deficient M27_AxAxxA_ MCMV mutant (Fig. 7E).

Lastly, to assess the relevance of these observations *in vivo*, mice were infected with WT-, ΔM27-, or M27_AxAxxA_-MCMVs, and virus titres in salivary glands were determined. Virus titre in ΔM27- and M27_AxAxxA_-MCMV infection were reduced strongly in comparison to WT-MCMV (Fig. 7G). This highlights the importance of M27 for MCMV replication and shows that the M27_AxAxxA_ mutation severely impairs the IFN inhibitory capacity in the mouse model. In conclusion, the disruption of the M27 zinc-binding motif causes a loss of DDB1 binding and STAT2 degradation, an impairment of IFN antagonism, and ultimately a virus attenuation *in vivo*. This establishes the predictive power of our atomic E27 model also for the MCMV infection model by connecting it with the MCMV *in vivo* replication capacity.

## Discussion

We identified STAT2, the central hub driving the antiviral IFN response, as one of the most drastically down-regulated host factors during RCMV-E infection and elucidated E27 as causative factor. The structure determination of E27 uncovered a zinc-binding motif close to the DDB1-binding site, highly conserved in homologous proteins from related cytomegaloviruses. Disruption of this motif in the closely related E27 homologue M27 resulted in impairment of DDB1 binding, STAT2 degradation, and ISG suppression, and finally to inhibition of virus replication *in vivo*. Similarly, the integrity of conserved zinc-coordinating motifs has been shown to be crucial for CRL4 association and function of SV5 V (Lin et al., 1998) and the retroviral accessory proteins Vpx and Vpr (Wang et al., 2017; Yamamoto et al., 2017). However, HBx side chains involved in zinc coordination seem to be dispensable for CRL4 interaction, but critical for substrate recruitment (Ramakrishnan et al., 2019).

With UL27, HCMV encodes a positional and sequence homologue of M27 and E27. HCMV reduces STAT2 levels (Le et al., 2008; Nightingale et al., 2018) and - like MCMV - exploits the UPS and CRLs to this end (Le-Trilling et al., 2016; Le et al., 2008). However, UL27 is neither essential nor sufficient for the HCMV-induced STAT2 degradation (Le et al., 2008), and at best interacts very weakly with DDB1 (Landsberg et al., 2018; Reitsma et al., 2011; Trilling et al., 2011). Our structure-based sequence alignment shows exchange of the third zinc-coordinating cysteine to a glycine in UL27, alongside considerable sequence divergence in the critical DDB1-binding helix α8, thereby rationalising these observations. In addition, only few of the E27 residues mediating the critical interaction with the STAT2 CCD are conserved in UL27. In accordance with these observations, HCMV UL27 induces the degradation of the unrelated host Tip60 acetyltransferase, eventually via direct interaction with proteasome components (Reitsma et al., 2011).

The HCMV-encoded STAT2 antagonist was recently identified as UL145 (Le-Trilling et al., 2020). Like M27, UL145 interacts with DDB1 and CRL4 (Nightingale et al., 2018) and induces proteasomal degradation of STAT2 (Le-Trilling et al., 2020). Accordingly, an unbiased single cell sequencing approach combined with a genome-wide CRISPR screen confirmed the central importance of the IFNAR2/STAT2/IRF9 signalling axis as well as neddylation and Cullin-RING E3s in defining HCMV infection propensity (Hein and Weissman, 2021). Sequence alignments from previous and from our present study indicate the presence of an H-box motif in the N-terminus of UL145 (residues 25-36, Fig. 4D), suggesting that this UL145 region is positioned similarly in the DDB1-binding cleft as endogenous DCAF substrate receptors and the C-terminal half of E27 helix α8. Indeed, site-directed mutagenesis of several putative H-box residues in UL145 abrogated DDB1 interaction in co-immunoprecipitation experiments (Le-Trilling et al., 2020), and a crystal structure of the UL145 H-box peptide in complex with DDB1 very recently supplied the ultimate proof of this hypothesis (Wick et al., 2022). However, future structural studies are necessary to clarify how the whole of UL145 integrates into CRL4, in order to pinpoint additional DDB1-interacting motifs, and to decipher the mechanisms of STAT2 recruitment. These will be highly informative to fully understand the different mechanistic principles pursued by HCMV and rodent CMV for STAT2 antagonism, both converging on CRL4.

The comparison of the DDB1-binding modes of E27 and other CRL4 substrate receptors revealed that E27 engages similar surface regions in the DDB1 binding cleft as DCAF-like receptors. However, structure and sequence determinants in E27 and other viral and cellular DCAFs mediating DDB1-binding are highly divergent. In fact, the only common element in these DDB1 binders with structural homology, and at least a certain degree of sequence conservation is the short H-box motif (Fig. 4C, D) (Li et al., 2010), interacting with the bottom of the binding cleft formed by DDB1 BPA and BPC. The physico-chemical properties of this DDB1 cleft seem to select for helical secondary structure, for an L or V side chain at position 7 of the sequence alignment in Fig. 4D (albeit with exception of the viral DCAFs HBx and SV5V), and with less stringency for R at position 10, L/I/V at position 12, and G at position 13. The latter G residue, having a very low propensity to be located in α-helices (Pace and Scholtz, 1998), might be necessary as helix-breaking residue to lead over into the loop segment exiting the binding cleft.

Our biochemical analysis indicated that E27 replaces the endogenous DCAF1 substrate receptor on DDB1. This is in agreement with recent results demonstrating that H-box peptides of the vDCAFs UL145 and HBx exhibit significantly higher DDB1-binding affinities than the corresponding H-box peptides derived from endogenous DCAF1 and DDB2 receptors (Wick et al., 2022). Accordingly, it is tempting to speculate that vDCAFs, in addition to marking antiviral substrates for degradation, might block a plethora of cellular DCAF-dependent ubiquitylation processes. Further research is needed to delineate if and how virus replication would benefit from these putative ubiquitylation inhibition events.

Interestingly, another CRL4 substrate receptor, CRBN, possesses a unique DDB1-binding mechanism involving three α-helices, which do not structurally align to the H-box motif at all (Chamberlain et al., 2014; Fischer et al., 2014). This pronounced plasticity of the DDB1 substrate receptor binding cleft impedes sequence-based prediction, and suggests the existence of further, yet unidentified viral and endogenous DDB1 binders. In line with these observations, recent HCMV interactome studies discovered several more DDB1- and CRL4-associated HCMV factors (Nightingale et al., 2022; Nobre et al., 2019). Accordingly, future studies of DDB1-binding mechanisms, together with proteomics and novel *in silico* structure prediction methods (Humphreys et al., 2021; Jumper et al., 2021), might greatly expand the spectrum of known DDB1-binders and extend our understanding of CRL4 biology and its viral exploitation.

Our study has limitations. We employed human DDB1 for *in vitro* reconstitution and structure determination of the complex with the RCMV factor E27 and rat STAT2. It is important to note that DDB1 is highly conserved across eukaryotes (Tang and Chu, 2002), because it is critical for development and cell viability as essential component of the nucleotide excision DNA repair machinery (Cang et al., 2006; Wakasugi et al., 2007). As a consequence, out of 1140 DDB1 amino acids, there are only 12 positions exchanged between human and rat DDB1, of which four are type-conserved (Fig. S12A). We have mapped these exchanges on a model of full length human DDB1 bound to E27 and the rat STAT2 CCD, to show that none of these are involved in intermolecular interaction, demonstrating the validity of our approach (Fig. S12B, C). In support of these observations, there is precedence in studying inter-species DDB1 protein complexes, e.g., the SV5V, WHX, SIV Vpx and Vpr, zebrafish DDB2 and chicken CRBN proteins have been *in vitro* reconstituted with human DDB1 for structural characterisation, and valid biological conclusions have been drawn from these studies (Angers et al., 2006; Banchenko et al., 2021; Fischer et al., 2014; Fischer et al., 2011; Li et al., 2006; Li et al., 2010; Schwefel et al., 2015; Schwefel et al., 2014; Wu et al., 2015).

We used a DDB1 mutant lacking the entire BPB for XL-MS and structure determination, while we employed full length DDB1 for analytical GF experiments. This had technical reasons, since protein complex preparations including full length DDB1, E27 and STAT2 did not yield cryo-EM reconstructions at appropriate resolution for model building, most likely due to the flexibility of the DDB1 BPB, impeding proper alignment of the cryo-EM particle images. In addition, the structure model including full length DDB1 (Fig. S12B) clearly demonstrates that the BPB is located on the opposite side of the E27 binding interface on DDB1, where it is situated to flexibly connect to the CUL4 scaffold to create the ubiquitination zone around immobilised substrates (Fig. 6). This rules out any involvement of DDB1 BPB in E27- and/or STAT2-binding processes, and thus validating the use of DDB1_ΔBPB_ in experiments aimed at studying E27- and/or STAT2-binding mechanisms. Along these lines, several previous studies have employed DDB1_ΔBPB_ to facilitate structure determination, and have successfully applied the resulting structural models for functional follow-up experiments in the context of complete CRL4 assemblies (Bussiere et al., 2020; Petzold et al., 2016; Slabicki et al., 2020).

Lastly, it should be noted, that we used a cross-linked protein complex sample for cryo-EM structure determination. Again, this had technical causes, since the equivalent non-cross-linked sample failed to produce cryo-EM specimens suitable for single particle analysis, likely because of aggregation and/or denaturation of the particles in the plunge-freezing process. Even though cross-linking is a common procedure to optimise cryo-EM samples (Drulyte et al., 2018; Stark, 2010), and XL-MS has been proven to be a suitable tool to inform modelling of native structures (O’Reilly and Rappsilber, 2018; Piersimoni et al., 2022; Schneider et al., 2018), we cannot exclude that the cross-linking procedure stabilised a rare or non-native conformation of the protein sample.

Our data, to our knowledge for the first time, reveal that RCMV (like MCMV) exploits host CRL4 complexes to induce proteasomal degradation of STAT2. With UL145, HCMV relies on an analogous, but evolutionary unrelated factor, also exploiting DDB1 and CRL4 for STAT2 degradation and evasion from the antiviral activity of IFNs. In clear contrast to HCMV, RCMV and MCMV are both amenable to *in vivo* experiments in small animal models. Over 40 years ago, HCMV has been called the troll of transplantation (Tx) due to its grim impact on immunosuppressed individuals after Tx (Balfour, 1979). Despite tremendous efforts, HCMV still harms and kills graft recipients. While MCMV allows various experiments regarding general principles of cytomegaloviral pathogenesis and antiviral immunity, one shortcoming is that mice are rather small animals in which chirurgical and solid organ transplantation (SOT) procedures are difficult. In clear contrast, various SOT procedures that are indispensable for human medicine can be recapitulated in bigger rat models. We hope that our work provides the molecular foundation for future studies addressing the relevance of STAT2-dependent IFN signalling, CMV-induced STAT2 degradation, and viral DDB1-CRL4 exploitation in rat SOT models.

## Acknowledgments

This research was supported by the German Research Foundation (DFG) Emmy Noether Programme SCHW1851/1-1 (D.S.), by an EMBO Advanced grant aALTF-1650 (D.S.), and by an iNEXT grant 5419 (D.S.). M.T. receives funding from the DFG through RTG1949, TR1208/1-1 and TR1208/2-1. We thank the MPI-MG for granting access to the TEM instruments of the microscopy and cryo-EM service group; Dominik A. Megger for initial mass spectrometry analyses; Birgit Zülch and Kristin Fuchs for technical support.

## Author Contributions

V.T.K.L.-T., S.B., A.G., J.B., T.B., B.S., T.M., J.R., C.M.T.S., S.V., M.T., D.S. planned experiments. V.T.K.L.-T., D.P., S.B., L.B., A.G., C.G., J.B., T.B., B.S., S.V., M.T., D.S. performed experiments. V.T.K.L.-T., D.P., S.B., S.L., A.G., T.B., B.S., J.R., S.V., M.T., D.S. analysed data. S.V., M.T., D.S. wrote the initial draft, all authors reviewed and edited the manuscript.

## Declaration of Interests

The authors declare no competing interests.

## Methods

### Animals

BALB/c mice were obtained from Harlan and bred and housed in the animal facility of the Institute for Virology of the University Hospital Essen. Mice were maintained under selective pathogen-free conditions. Age (8-16 weeks) and sex-matched (male and female) groups of BALB/c mice were infected intraperitoneally with 2× 10^5^ PFU of WT-MCMV or MCMV mutants per mouse. All procedures were performed in accordance with the guidelines of the University Hospital Essen, Germany, the national animal protection law (Tierschutzgesetz [TierSchG]) and the recommendations of the Federation of European Laboratory Animal Science Association (FELASA). The study was approved by the Northrhine-Westphalia State Office for Nature, Environment and Consumer Protection (LANUV NRW, Düsseldorf, Northrhine-Westphalia, Germany; permit #84-02.04.2016.A441).

### Microbe strains

*E. coli* NovaBlue and Rosetta™ 2(DE3) were cultivated in LB medium at 37°C in a shaker incubator (150 rpm) (Table S3).

### Cell lines

Immortalised mouse fibroblasts were generated from primary C57BL/6 and BALB/c MEF by crisis immortalisation (Rattay et al., 2015). NIH3T3-ISREluc:IFNLR cells were described in (Le-Trilling et al., 2018). Mouse fibroblasts, REF and HEK293T (ATCC CRL-11268, female) cells were grown in Dulbecco`s minimal essential medium (DMEM) supplemented with 10% fetal calf serum (FCS), penicillin, streptomycin, and 2 mM glutamine at 37°C in 5% CO2. Sf9 cells (originally derived from the ovaries of *Spodoptera frugiperda*) were cultivated in Insect-XPRESS^™^ medium at 28°C in a shaker incubator (180 rpm) (Table S3).

### Primary cultures

Organs of infected mice were harvested, snap frozen in liquid nitrogen, and stored at −80°C until titrations were performed. Plaque titrations were conducted by use of primary mouse fibroblasts (mouse embryonic fibroblasts [MEF] and mouse newborn cells [MNC]) which were isolated from mouse embryos and newborns, respectively, according to described protocols (Le-Trilling and Trilling, 2017). Fibroblasts were grown in DMEM supplemented with 10% (v/v) FCS, 100 µg/ml streptomycin, 100 U/ml penicillin, and 2 mM glutamine. All cell culture media and supplements were obtained from Gibco/Life technologies.

### Viruses

WT-RCMV-E has previously been described (Ettinger et al., 2012). ΔE27-RCMV-E was generated by CRISPR/Cas9-mediated genome editing. REF were transduced with a mixture of four CRISPR/Cas9 lentiviral vectors carrying sgRNAs that target viral genome up- and downstream of the E27 gene (see Table S3). Successfully transduced cells were selected with 2.5 *μ*g/mL puromycin prior to infection with WT-RCMV-E. This resulted in an E27-knockout virus (ΔE27) with a complete deletion of the 1970 bp *E27* gene. E27 deletion was verified by PCR and sequencing. Mutant virus was screened by limiting dilution and was double plaque-purified. WT-MCMV was reconstituted from the BAC described in (Jordan et al., 2011). ΔM27-MCMV has been previously described (Le-Trilling et al., 2018). The M27_AxAxxA_ mutant was generated according to previously published procedures (Tischer et al., 2006; Wagner et al., 2002) using the primers MCMV-M27-QC1-3_1 and MCMV-M27-QC1-3_2 (Table S4). Viral titers were determined by standard plaque titration on primary MEF, REF or MNC.

### Plasmids

For CRISPR/Cas9 viral genome editing, crRNA sequences targeting the viral genome up- and downstream of the E27 gene were designed manually or using the CRISPOR website (http://crispor.tefor.net); crRNA sequences are presented in Table S4 and were cloned as sgRNA downstream a human U6 promoter in the pSicoR-CRISPR-PuroR vector (van Diemen et al., 2016). The pHisSUMO-E27 and pHisSUMO-*Rattus norvegicus* (rn)STAT2 expression plasmids were prepared using standard restriction enzyme cloning. Full-length open reading frames were PCR-amplified from cDNA templates using oligonucleotide primers XmaI-E27-fw and SacI-E27-rev or XmaI-STAT2-fw and NotI-STAT2-rev, and ligated in the pHisSUMO plasmid, to generate N-terminal 6xHis-SUMO-fusion proteins. The rnSTAT2 cDNA template was ordered as *E. coli* codon-optimised Strings DNA fragment from ThermoFisher. To generate pAcGHLT-B-3C-*Homo sapiens* (hs)DDB1, the thrombin protease site between the GST purification tag and DDB1 in pAcGHLT-B-hsDDB1 was changed to a 3C protease site using insertion/deletion PCR according to (Hansson et al., 2008), using the oligonucleotide primers 3C-DDB1-A and 3C-DDB1-B. To generate pAcGHLT-B-3C-hsDDB1 ΔBPB (amino acid residues 396-705 replaced with a GNGNSG sequence), and HisSUMO-E27Δ (residues 284-305 replaced with a GNGNSG linker), the same method was used, with the oligonucleotide primers ΔBPB-A and ΔBPB-B or E27Δ-A and E27 E27Δ-B, and pAcGHLT-B-3C-hsDDB1 or pHisSUMO-E27 as template. Point mutations were introduced by site-directed mutagenesis using the primers E27-R211E/R214E-fw and E27-R211E/R214E-rev, and pHisSUMO-E27 as template. The M27-FLAG expression plasmid was described previously (Trilling et al., 2011). Site-directed mutations were introduced using the M27-QC primers and the QuikChangeII kit (Agilent) according to manufacturer’s instructions. For primer sequences, see Table S4. All constructs were verified by Sanger sequencing.

### LC-MS/MS Analyses

Liquid chromatography couple to mass spectrometry analyses were performed as described previously (Megger et al., 2017). Briefly, cells were harvested and dissolved in 50 mM ammoniumbicarbonat containing 0.1% (w/v) Rapigest SF Surfactant (Waters, Boston, MA, USA). Four µg proteins per sample were reduced with dithiothreitol for 30 min at 60°C and thereafter alkylated with 15 mM iodoacetamide for 30 min at room temperature. Subsequently, the samples were digested using trypsin (SERVA Electrophoresis, Heidelberg, Germany) in a ratio of 1:20 (w/w) for 16 h at 37 °C. The reaction was stopped by adding trifluoroacetic acid (TFA) to final 1% (v/v) and the precipitated surfactant was removed by centrifugation at 16.000 x *g* for 10 min. The supernatants were dried by vacuum centrifugation and dissolved in 0.1 % TFA. An amount of 300 ng tryptic peptides per sample was used for LC-MS/MS analysis and injected into an Ultimate 3000 RSLC nanoLC online coupled to an Orbitrap Elite mass spectrometer (both Thermo Scientific). The peptides were concentrated for 7 min on the trap column (Acclaim PepMap 100, 75 µm × 2 cm, C18, 5 µm particle size, 100 Å pore size) using a flow rate of 30 µL/min. Subsequently, the samples were separated on the analytical column (Acclaim^®^ PepMap RSLC, 75 µm × 50 cm, nano Viper, C18, 5 µm particle size, 100 Å pore size) by applying a gradient from 5 to 40% solvent B over 98 min (solvent A: 0.1% (v/v) formic acid; solvent B: 0.1% (v/v) formic acid, 84% acetonitrile) at a flow rate of 400 nL/min. The column oven temperature was 60°C. Full scan MS spectra were acquired in the Orbitrap analyzer at a resolution of 60,000 (400 m/z) with a mass range from 350 to 2000 m/z. Tandem mass spectra of the 20 highest abundant precursors were acquired in the linear ion trap after fragmentation by collision-induced dissociation.

Peptide identification was carried out using Proteome Discoverer 1.4 (Thermo Scientific, Bremen, Germany). The acquired spectra were searched against the UniProt reference proteome for *Rattus norvegicus* (UP000002494, release 2015_11, 29880 entries) and UniProt/TrEMBL sequences for *Rat cytomegalovirus* (RCMV-E, isolate England; release 2015_11, 208 entries) using Mascot (v.2.5; Matrix Science, London, UK). Trypsin was set as digesting enzyme and one missed cleavage site was considered as well as dynamic oxidation of methionine and static carbamidomethylation of cysteine residues. The mass tolerance was set to 5 ppm for precursor ions and 0.4 Da for fragment ions. The false discovery rate (FDR) was calculated using the percolator function and only identifications passing an FDR threshold of 1% were considered. Label-free quantification was carried out using Progenesis QI for Proteomics (ver.2.0.5, Nonlinear Dynamics, Newcastle upon Tyne, UK) and the peptide spectrum matches derived from Proteome Discoverer were used for protein identification. Normalised protein abundancies were exported and further processed using R (R Foundation for Statistical Computing, Vienna, Austria). Statistical significance was calculated using acrsinh-transformed data by ANOVA followed by Tukey’s honest significant difference method. Ratios of means were calculated using non-transformed data. Proteins passing an ANOVA p_FDR_ value ≤ 0.05 (adjusted according to Benjamini-Hochberg), a post hoc test p value ≤ 0.05 and an absolute ratio of means ≥ 2 were considered as significantly differentially abundant.

### Immunoblotting

Immunoblotting was performed according to standard procedures. Briefly, cells were lysed as described (Trilling et al., 2009) and equal amounts of protein were subjected to SDS-PAGE and transferred to nitrocellulose membranes. Immunoblot analysis was performed using the specific antibodies recognizing the indicated proteins (Table S3). Proteins were visualised using peroxidase-coupled secondary antibodies (Dianova) and the ECL chemiluminescence system (Cell Signaling).

### Co-immunoprecipitation

Transfected HEK293T cells were washed with ice cold PBS and subsequently lysed as described (Le et al., 2011; Trilling et al., 2011) by incubation for 1 h on ice in the following buffer: 0.1 mM EDTA, 150 mM NaCl, 10 mM KCl, 10 mM MgCl2, 10% (v/v) glycerol, 20 mM HEPES (pH 7.4), 0.5% (v/v) NP-40, 0.1 mM PMSF, 1 mM DTT, 10 μM pepstatin A, 5 μM leupeptin, 0.1 mM Na-vanadate, Complete protease inhibitor (Roche). Lysates were cleared by centrifugation for 30 min at 4°C and 16,000 g. The IP antibody (Table S3) was added to the supernatant and incubated overnight at 4°C in a rotating device. Immune complexes were precipitated with Protein-G-Sepharose (GE Healthcare) for 1 h at 4°C in a rotating device and washed with 150, 250, and 400 mM NaCl-containing lysis buffer. Samples were separated by SDS-PAGE for immunoblot analysis of indicated proteins.

### Luciferase assays

NIH3T3 reporter cells were infected with MCMV, treated with IFNβ or IFNλ for 6 h, and lysed for quantification of reporter gene expression (Le-Trilling et al., 2018). Luciferase activity was measured according to the manufacturer’s instructions (pjk) using a microplate multireader (Mithras2 LB 943; Berthold Technologies). Quantification of luminescence was performed by use of the microplate multireader Mithras2 LB 943 and MicroWin software (Berthold Technologies). The resulting data were analyzed using GraphPad Prism software. The values are reported as Mean ± standard deviation (SD) as described in the figure legends.

### Protein production and purification

Constructs in the pHisSUMO vector, containing an N-terminal HisSUMO-tag, followed by a 3C protease cleavage site, were expressed in *Escherichia coli* Rosetta^™^ 2(DE3). In a typical expression, 2 L LB medium containing 100 µg/mL Ampicillin and 34 µg/mL Chloramphenicol were inoculated with 20 mL of an overnight culture and grown under shaking in an Multitron HT incubator (Infors) at 37°C, 150 rpm, until OD_600_=0.7. Then, temperature was lowered to 18°C and protein expression was induced by addition of 0.2 mM Isopropyl β-d-1-thiogalactopyranoside (IPTG). Cultures were grown for further 20 h. Bacteria were pelleted by centrifugation at 4000 rpm, 4°C for 15 min using a JLA-9.1000 centrifuge rotor (Beckman). The pellet was resuspended in 70 mL buffer containing 50 mM Tris-HCl, pH 7.8, 500 mM NaCl, 4 mM MgCl_2_, 0.5 mM tris-(2-carboxyethyl)-phosphine (TCEP), mini-complete protease inhibitors (1 tablet per 50 mL) (Merck), and 30 mM imidazole. 5 µL Benzonase (Merck) was added and the suspension was stirred thoroughly. Cells were lysed by passing the suspension at least twice through a Microfluidiser (Microfluidics). The lysate was clarified by centrifugation at 48000×g for 45 min at 4°C using a JA-25.50 rotor (Beckman), and applied on a XK 16/20 chromatography column (Cytiva) containing 10 mL Ni Sepharose® High Performance beads (Cytiva), using an Äkta pure FPLC (Cytiva) at 4°C. The column was washed with 250 ml buffer containing 50 mM Tris-HCl, pH 7.8, 500 mM NaCl, 4 mM MgCl_2_, 0.5 mM TCEP, and 30 mM imidazole. Protein was eluted with the same buffer supplemented with 0.3 M imidazole. Eluent fractions were analysed using SDS-PAGE, appropriate fractions were pooled and concentrated to 5 mL with centrifugal filter devices (Vivaspin 20, Sartorius). In pHisSUMO-rnSTAT2 preparations, 30 µg 3C protease per mg total protein was added and the sample was incubated for 12 h on ice to cleave off the HisSUMO-tag. In pHisSUMO-E27 purifications, the tag was not removed, because tag-free E27 precipitated out of solution at higher concentration in the absence of additional binding partners. Subsequently, gel filtration chromatography (GF) was performed on an Äkta prime plus FPLC (Cytiva), using a Superdex 200 16/600 column (Cytiva), equilibrated in 10 mM Tris-HCl pH 7.8, 150 mM NaCl, 4 mM MgCl_2_, 0.5 mM TCEP buffer, at a flow rate of 1 mL/min. GF fractions were analysed by SDS-PAGE, appropriate fractions were pooled and concentrated to approx. 20 mg/mL, flash-frozen in liquid nitrogen in small aliquots and stored at −80°C. Protein concentration was determined with a NanoDrop spectrophotometer (ND 1000, Peqlab), using theoretical absorption coefficients calculated based upon the amino acid sequence by ProtParam on the ExPASy webserver (Wilkins et al., 1999).

Constructs in the pAcGHLT-B vector, encoding an N-terminal GST-His-tag, followed by a 3C protease cleavage site, were expressed by infecting Sf9 cells with recombinant baculoviruses (*Autographa californica nucleopolyhedrovirus* clone C6), which were generated as described previously (Zhao et al., 2003). Sf9 cells were cultivated in Insect-XPRESS medium (Lonza) at 28°C, 180 rpm in an Innova 42R incubator shaker (New Brunswick). 1 L Sf9 cells at a cell density of 3×10^6^ cells/mL were infected with 0.5 mL of high titre recombinant baculovirus for 72 h. Cells were pelleted by centrifugation at 1000 rpm, 4°C for 30 min using a JLA-9.1000 centrifuge rotor (Beckman). The pellet was resuspended in 500 mL hypotonic lysis buffer containing 10 mM Tris-HCl pH 7.8, 4 mM MgCl_2_, 30 mM imidazole, 0.5 mM TCEP, mini-complete protease inhibitors (10 tablets) (Merck) and stirred at 4°C for 30 min. The lysate was cleared by centrifugation at 4000×g, 4°C for 30 min. Subsequent purification steps were performed as described above for proteins expressed in *E. coli*.

### Analytical gel filtration (GF)

10 µM of each protein component was incubated with 30 µg 3C protease in a volume of 150 µL buffer containing 50 mM Tris-HCl, pH 7.8, 500 mM NaCl, 4 mM MgCl_2_, and 0.5 mM TCEP, for 12 h on ice. To remove the cleaved purification tags and GST-tagged 3C protease, 20 μL GSH-Sepharose FF beads (Cytiva) were added and the sample was rotated at 4 °C for 1 h. GSH-Sepharose beads were removed by centrifugation at 4°C, 4000 rpm for 5 min, and 120 µL of the supernatant was loaded on an analytical GF column (Superdex 200 Increase 10/300 GL, Cytiva), equilibrated in 10 mM Tris-HCl pH 7.8, 150 mM NaCl, 4 mM MgCl_2_, and 0.5 mM TCEP, at a flow rate of 0.5 mL/min, using an Äkta pure FPLC (Cytiva). 1 mL fractions were collected and analysed by SDS-PAGE.

### Cryo-EM sample preparation

7 µM HisSUMO-E27, 7 µM GST-DDB1_ΔBPB_, and 7 µM rnSTAT2 were incubated with 0.5 mg 3C protease in a volume of 1 mL buffer containing 10 mM HEPES-NaOH pH 7.8, 150 mM NaCl, 4 mM MgCl_2_, 0.5 mM TCEP, for 12 h on ice. Subsequently, the sample was subjected to GF chromatography on an Äkta prime plus FPLC (Cytiva), using a Superdex 200 16/600 column (Cytiva), connected in line to a 1 mL Glutathione Sepharose® 4 Fast Flow column (Cytiva), at a flow rate of 1 mL/min, using the same buffer as eluent. GF fractions were analysed by SDS-PAGE. Fractions corresponding to the elution peak containing E27, DDB1_ΔBPB_ and rnSTAT2 were pooled and concentrated to 70 µL (protein concentration 4.1 mg/mL).

Suberic acid bis(N-hydroxysuccinimide ester) (DSS) cross-linker (Merck) was added 1:1 (w/w) from a 100 µg/mL stock solution in DMSO, and the sample volume was adjusted to 400 µL with buffer containing 10 mM HEPES-NaOH pH 7.8, 150 mM NaCl, 4 mM MgCl_2_, 0.5 mM TCEP. The sample was incubated for 1 h at 22°C, and the cross-linking reaction was quenched by the addition of 20 µL 1 M Tris-HCl pH 7.8 for 10 min at 22°C. Precipitate was removed by centrifugation at 16.000×g, 22°C for 5 min. The supernatant was loaded on a Superdex 200 Increase 10/300 GL GF column, equilibrated in 10 mM Tris-HCl pH 7.8, 150 mM NaCl, 4 mM MgCl_2_, and 0.5 mM TCEP, at a flow rate of 0.5 mL/min, using an Äkta pure FPLC (Cytiva). 0.5 mL fractions were collected and analysed by SDS-PAGE. Fractions containing the cross-linked DDB1_ΔBPB_/E27/STAT2 complex were pooled, concentrated to 1.3 mg/mL, flash-frozen in 5 µL aliquots in liquid nitrogen and stored at −80°C.

R1.2/1.3 400 mesh Cu holey carbon grids (Quantifoil) were glow-discharged for 30 s using a Harrick plasma cleaner with technical air at 0.3 mbar and 7 W. 3.5 µL solution containing 0.5 µM (0.13 mg/mL) cross-linked DDB1_ΔBPB_/E27/STAT2 complex were applied to the grid, incubated for 45 s, blotted with a Vitrobot Mark IV device (Thermo Fisher) for 1-2 s at 4°C and 99% humidity, and plunged in liquid ethane. Grids were stored in liquid nitrogen until imaging.

### Cryo-EM data collection and processing

Data was collected on a 300 kV Tecnai Polara cryo-EM (Thermo Fisher) equipped with a K2 Summit direct electron detector (Gatan) in super-resolution mode, at a nominal magnification of 31000×, with a pixel size of 0.63 Å/px and the following parameters: defocus range of −1 to −2.5 µm, 50 frames per movie, 200 ms exposure per frame, electron dose of 6.2 e/Å^2^/s, leading to a total dose of 62 e/Å^2^ per movie. In four independent sessions, data from four grids were collected using Leginon (Suloway et al., 2005) yielding 8.069 movie stacks. Movies were aligned and dose-weighted using MotionCor2 (Zheng et al., 2017). Initial estimation of the contrast transfer function (CTF) was performed with the CTFFind4 package via the Relion 3.0.7 GUI (Mindell and Grigorieff, 2003; Scheres, 2012; Zivanov et al., 2018). Power spectra were manually inspected to remove ice contamination and astigmatic, weak, or poorly defined spectra. Particles were picked using Gautomatch 0.53 (https://www2.mrc-lmb.cam.ac.uk/research/locally-developed-software/zhang-software/) or the Relion 3.0.7 autopicker as indicated in Fig. S3. All subsequent image processing procedures were performed using Relion 3.0.7 or cryoSPARC 3.1 (Punjani and Fleet, 2021; Punjani et al., 2017; Punjani et al., 2020), and are shown in Fig. S3.

### Model building and refinement

Guided by the cryo-EM density map 1 (Fig. S3) and XL-MS distance restraints (Fig. 2C), a partial molecular model of E27 was generated *de novo* using the program Coot 0.9.5 (Emsley and Cowtan, 2004; Emsley et al., 2010). In addition, *in silico* AlphaFold2 structure prediction of E27 was performed (Jumper et al., 2021) using the ColabFold web server (https://colab.research.google.com/github/sokrypton/ColabFold/blob/main/AlphaFold2.ipynb) (Mirdita et al., 2021). The AlphaFold2 output coordinates were in very good agreement with the experimentally determined E27 structure (RMSD of 1.489 Å, 2764 aligned atoms). Accordingly, the model was completed by merging the initial model with the AlphaFold2 template and adjusting backbone and side chain positions where necessary using Coot 0.9.5. DDB1 coordinates were taken from a previously solved DDB1/DCAF1-CtD complex structure (Banchenko et al., 2021), rigid body-fitted and adjusted manually using Coot 0.9.5. The coordinates of the rnSTAT2 CCD were extracted from the AlphaFold protein structure database (https://alphafold.ebi.ac.uk/entry/Q5XI26) (Varadi et al., 2022), rigid body-fitted and adjusted manually using Coot 0.9.5. As final step, automated real space refinement was performed using the program Phenix 1.19.2-4158 (Liebschner et al., 2019). The refined DDB1/E27/STAT2 CCD structure was then fitted as rigid body in cryo-EM density map 2 (Fig. S3). Full-length rnSTAT2 coordinates (https://alphafold.ebi.ac.uk/entry/Q5XI26) were placed by structural superposition of the CCDs. The STAT2 NTD and TAD were removed, and flexible loops in the SH2 domain were trimmed, because they could not be assigned to the experimental density, indicating that they are flexibly attached and mobile relative to the other STAT2 domains. Lastly, a chain break was introduced between STAT2 residues 315 and 317, and the STAT2 portion encompassing residues 317-701 (DBD, LD, SH2) was once more separately fitted as rigid body, to account for a small movement relative to the CCD. Subsequently, the model was subjected to rigid body and grouped b-factor real space refinement using Phenix 1.19.2-4158, followed by molecular dynamics flexible fitting using the Namdinator server (Kidmose et al., 2019), applying default parameters.

### Cross-linking mass spectrometry

#### Photo-Crosslinking

The cross-linker sulfo-SDA (sulfosuccinimidyl 4,4′-azipentanoate) (Thermo Scientific) was dissolved in cross-linking buffer (10 mM HEPES-NaOH, pH 7.8, 150 mM NaCl, 4 mM MgCl, 0.5 mM TCEP) to 100 mM before use. The labelling step was performed by incubating 23 μg aliquots of the DDB1_ΔBPB_/E27/DDA1/STAT2 complex at 1 mg/mL with 0.2, 0.3, 0.6 and 0.8 mM sulfo-SDA, added, respectively, for an hour. The samples were then irradiated with UV light at 365 nm using a Luxigen LZ1 LED emitter (Osram Sylvania Inc.), to form cross-links, for 10 seconds and quenched with 50 mM Tris.HCl pH 7.5 for 20 min. All steps were performed on ice. Reaction products were separated on a Novex Bis-Tris 4–12% SDS−PAGE gel (Life Technologies) in order to identify suitable cross-linker concentrations. Samples corresponding to 0.2 and 0.3 mM SDA concentrations were precipitated in 80% acetone at −20 °C. All subsequent steps were carried out at room temperature. Protein pellets were resuspended in denaturing buffer (50 mM NH_4_HCO_3_, 8 M urea), reduced with 2 mM DTT for 30 minutes and alkylated with 5 mM iodoacetamide for 30 minutes. Urea concentration was then reduced to 1.5 M by dilution with 50 mM NH_4_HCO_3_ and samples were digested with trypsin (Thermo Scientific Pierce) (Shevchenko et al., 2006) overnight. The resulting tryptic peptides were extracted, desalted using C18 StageTips (Rappsilber et al., 2003) and pooled. Eluted peptides were fractionated on a Superdex Peptide 3.2/300 increase column (GE Healthcare) at a flow rate of 10 µL/min using 30% (v/v) acetonitrile and 0.1 % (v/v) trifluoroacetic acid as mobile phase. 50 μL fractions were collected and vacuum-dried.

#### XL-MS acquisition

Samples for analysis were resuspended in 0.1% (v/v) formic acid, 3.2% (v/v) acetonitrile. LC-MS/MS analysis was performed on an Orbitrap Fusion Lumos Tribrid mass spectrometer (Thermo Fisher) coupled on-line with an Ultimate 3000 RSLCnano HPLC system (Dionex, Thermo Fisher). Samples were separated on a 50 cm EASY-Spray column (Thermo Fisher). Mobile phase A consisted of 0.1% (v/v) formic acid and mobile phase B of 80% (v/v) acetonitrile with 0.1% (v/v) formic acid. Flow rates were 0.3 μL/min using gradients optimised for each chromatographic fraction from offline fractionation, ranging from 2% mobile phase B to 55% mobile phase B over 90 min. MS data were acquired in data-dependent mode using the top-speed setting with a 2.5 s cycle time. For every cycle, the full scan mass spectrum was recorded using the Orbitrap at a resolution of 120,000 in the range of 400 to 1,500 m/z. Ions with a precursor charge state between 3+ and 7+ were isolated and fragmented. Analyte fragmentation was achieved by stepped Higher-Energy Collisional Dissociation (HCD) and a decision tree strategy (Kolbowski et al., 2017). Fragmentation spectra were then recorded in the Orbitrap with a resolution of 60,000. Dynamic exclusion was enabled with single repeat count and 60 s exclusion duration.

#### XL-MS processing

A recalibration of the precursor m/z was conducted based on high-confidence (<1% false discovery rate (FDR)) linear peptide identifications. The re-calibrated peak lists were searched against the sequences and the reversed sequences (as decoys) of cross-linked peptides using the Xi software suite (v.1.7.6.3) for identification (Mendes et al., 2019). Final crosslink lists were compiled using the identified candidates filtered to <2% FDR on link level with xiFDR v.2.1.5.5 (Fischer and Rappsilber, 2017) imposing a minimum of 5 observed fragments per peptide and at least one crosslinker-containing fragment. FDR settings were optimised by boosting heteromeric crosslinks based on delta score, minimum fragments and minimum sequence coverage, and lower result levels.

#### DisVis

Center of mass position of the DDA1 C-terminal region relative to the complex was determined using DisVis (van Zundert and Bonvin, 2015; van Zundert et al., 2017). The DDB1/STAT2/E27 complex structure from this study was selected as the fixed chain and residues 49-76 of DDA1 from PDB 6ue5 (Bussiere et al., 2020) were selected as the scanning chain. 46 SDA crosslinking restraints were entered from Cα to Cα with allowed distance between 2.5 Å and 22 Å. The rotational sampling was set to 15°.

#### Sequence alignments

Sequence alignments were calculated using the EBI ClustalOmega server (Madeira et al., 2019) (https://www.ebi.ac.uk/Tools/msa/clustalo/), and adjusted manually using the program GeneDoc (Nicholas, 1997).

#### Virus taxonomy IDs (TaxID)

HCMV, 10359; MuHV-1, 10366; MuHV-2, 79700; MuHV-8, 1261657

#### Amino acid sequences used for multiple sequence alignments

E27, NCBI YP_007016432; M27, NCBI AWV68118; R27, NCBI NP_064132; T27, NCBI NP_116370; B27, NCBI AFK83839; rhesus macaque UL27 (RH27), NCBI AAZ80544; UL27, NCBI ACN63126; DCAF1, UniProt Q9Y4B6; DCAF2, UniProt Q9NZJ0; DCAF4, UniProt Q8WV16; DCAF5, UniProt Q96JK2; DCAF6, UniProt Q58WW2; DCAF8, UniProt Q5TAQ9; DCAF9, UniProt Q8N5D0; DCAF10, UniProt Q5QP82; DCAF11, UniProt Q8TEB1; DCAF12, UniProt Q5T6F0; DCAF14, UniProt Q8WWQ0; DCAF15, UniProt Q66K64; DCAF17, UniProt Q5H9S7; AMBRA1, UniProt Q9C0C7; CSA, UniProt Q13216; DDB2, UniProt Q92466; HBX, NCBI AMH41022; WHX, NCBI AAA46776; SV5V, UniProt P11207; UL145, UniProt F5HF44; rat STAT2, UniProt Q5XI26; mouse STAT2, UniProt Q9WVL2; human STAT2, UniProt P52630; rhesus macaque STAT2, UniProt F6R1P6; pig STAT2, UniProt O02799; chicken STAT2, UniProt A0A1D5PIV4; frog STAT2, UniProt A0A1L8HHX5

### Data availability

LC-MS data have been deposited at ProteomeXchange via the PRIDE partner repository (Perez-Riverol et al., 2019) with the dataset identifier PXD033639 (reviewer account username, password: KsIRcYHZ). XL-MS data have been deposited at the JPOST ProteomeXChange (Deutsch et al., 2020) with accession number JPST001566 (ProteomeXChange PXD033366, reviewer details: https://repository.jpostdb.org/preview/110121477262651feae5f48, access key: 3943). The EM maps have been deposited at the Electron Microscopy Data Bank (EMDB) with the accession numbers EMD-14802 (map 1) and EMD-14812 (map 2). Atom coordinates have been deposited at the Protein Data Bank (PDB) with the accession numbers 7zn7 (DDB1_ΔBPB_/E27/STAT2 CCD) and 7znn (DDB1_ΔBPB_/E27/STAT2 FL).

## Supplementary Information

**Figure S1: Proteomic analyses of protein abundance changes upon RCMV-E infection, related to Figure 1**

**(A)** Mass spectrometric analyses of protein abundance changes upon RCMV-E infection, compared to mock-infected samples, at indicated time points. Significantly down-or upregulated proteins are marked in red.

**(B)** Relative abundance changes of selected proteins involved in IFN signalling upon RCMV-E infection compared to mock-infected samples at indicated time points.

**Figure S2: Cryo-EM sample preparation, related to Figures 3-5**

**(A)** GF chromatogram of the disuccinimidyl suberate (DSS) cross-linked DDB1_ΔBPB_/DDA1/E27/STAT2 protein complex. GF fractions that have been pooled for cryo-EM analysis are indicated.

**(B)** SDS-PAGE analysis of the cross-linking reaction and GF fractions.

**Figure S3: Cryo-EM data processing flowchart, related to Figures 3-5**

If not stated otherwise, all computational procedures have been performed using the program Relion 3.0.7 (Zivanov et al., 2018). A crystal structure of hexameric ArnA, PDB 4wkg (Fischer et al., 2015; Yang et al., 2019), was fitted in the contamination density (red box).

**Figure S4: Molecular modelling of DDA1, related to Figure 3**

**(A)** Superposition of a DDB1_ΔBPB_/DDA1/DCAF15 complex structure (PDB 6q0v) (Faust et al., 2020) with DDB1_ΔBPB_/E27 complex coordinates. Proteins are shown as cartoons, DDB1 is coloured in blue, E27 in green, and DDA1 in red. DCAF15 has been removed for clarity, and only DDB1 from the DDB1/E27 structure is shown. DDA1 termini and the C-terminal helix are indicated. Experimental cryo-EM map 1 is shown as semi-transparent surface.

**(B)** Visualisation (grey semi-transparent surface) of the space accessible to the centre of mass of the DDA1 C-terminal helix, calculated based on cross-link distance restraints, consistent with at least 15 of the observed cross-links involving DDA1, using the DisVis software tool (van Zundert and Bonvin, 2015; van Zundert et al., 2017). Proteins are represented as in **A**. This analysis implies that the C-terminal helix preferentially locates to a relatively large area around the edges of DDB1 BPC and CTD, and does not occupy a defined location in contact with E27, as observed in the DDB1_ΔBPB_/DDA1/DCAF15 complex (indicated by the red arrow).

**Figure S5: Cryo-EM map 1 quality, AlphaFold2 models, model fit, and CLMS analysis, related to Figure 3**

**(A)** Overview of DDB1_ΔBPB_ model-to-map fit. DDB1_ΔBPB_ is shown as blue cartoon, cryo-EM density as grey semi-transparent surface (threshold level 0.555, Chimerax software tool) (Pettersen et al., 2021).

**(B) (Left panel)** Superposition of the AlphaFold2 E27 model (rainbow-coloured from N-to C-terminus) with a partial structure (grey) manually built into cryo-EM map 1 (Fig. S3). Structures are shown as cartoons. Terminal regions of the AlphaFold2 model with very low confidence have been removed for clarity. Red arrows indicate regions with significant structural differences between predicted and experimental model. The black arrow and the red box indicate the zinc-binding motif. The zinc ion is shown as grey sphere. **(Middle panel)** Cartoon representation of the E27 AlphaFold2 model, coloured according to model confidence, i.e., the model’s prediction of its score on the local Distance Difference Test (lDDT-Cα) (Jumper et al., 2021). Terminal regions with very low confidence have been removed from the model for clarity. **(Right panel)** Overview of E27 model-to-map fit. E27 is shown as green cartoon, cryo-EM density as grey semi-transparent surface (threshold level 0.5, Chimerax software tool) (Pettersen et al., 2021).

**(C)** Close-up view of the zinc-binding site marked as red box in **B**. E27 is shown as green cartoon, with side chains involved in zinc coordination in stick representation. The zinc ion is shown as grey sphere. Coordination bond lengths (in Å) are indicated. Cryo-EM density is shown as grey semi-transparent surface (threshold level 0.5, Chimerax software tool) (Pettersen et al., 2021). Note that the tetrahedral coordination geometry, the ligands (Cys, His), and the coordination bond lengths are typical for structural zinc-binding sites, as demonstrated by analysis of almost 25,000 zinc-binding motifs available in the PDB (Ireland and Martin, 2019; Laitaoja et al., 2013).

**(D) (Left panel)** Superposition of the AlphaFold2 STAT2 model (rainbow-coloured from N-to C-terminus) with the portion of the STAT2 CCD (grey) fitted into cryo-EM map 1 (Fig. S3). Structures are shown as cartoons. ND and TAD have been removed for clarity. **(Middle panel)** Cartoon representation of the STAT2 AlphaFold2 model, coloured according to model confidence, i.e., the model’s prediction of its score on the local Distance Difference Test (lDDT-Cα) (Jumper et al., 2021). **(Right panel)** Overview of STAT2 CCD model to map fit. STAT2 is shown as magenta cartoon, cryo-EM density as grey semi-transparent surface (threshold level 0.35, Chimerax software tool) (Pettersen et al., 2021).

**(E)** Intermolecular sulfo-SDA cross-links were mapped on the DDB1_ΔBPB_/E27/STAT2 CCD structure (Fig. 3A), colour-coded according to their length as indicated.

**Figure S6: CLMS analysis of DDB1/E27/STAT2 models derived from cryo-EM map 1 and map 2, related to Figures 3 and 5**

**(A)** Left panel: sulfo-SDA cross-links mapped onto the DDB1/E27/STAT2 model built in map 1. Both within protein and heteromeric links are displayed. Satisfied cross-links (shorter than 25 Å Cα-Cα) are coloured in blue, borderline violations (25-35 Å) in orange, and violated (> 35 Å) cross-links in red. Right panel: circle view of the links between proteins coloured by cross-link satisfaction. Grey cross-links lie between regions not modelled in the structure. Solid colounalogred bars indicate structural coverage.

**(B)** As in **A**, but for the model built in map 2. Violated cross-links between E27 α/β-domain, STAT2 and DDB1 may indicate a swivel motion of the top of the complex relative to DDB1, which is consistent with the high mobility of this module relative to the core of the complex observed in cryo-EM analysis.

**Figure S7: Side chain densities and mutational analysis of the DDB1**_**ΔBPB**_**/E27 interaction, related to Figure 3**

**(A)** Model-to-map fit of the DDB1_ΔBPB_/E27 interaction regions (Fig. 3B). DDB1_ΔBPB_ is shown as blue cartoon, with side chains involved in E27 interaction in stick representation. E27 is shown as green cartoon, with side chains mediating DDB1_ΔBPB_ interaction in stick representation. Cryo-EM density map 1 (Fig. S3) is shown as blue mesh, contoured at 5σ, using the the PyMOL Molecular Graphics System, Version 2.0 Schrödinger, LLC.

**(B)** *In vitro* reconstitution, followed by analytical gel filtration (GF) analysis, of protein complexes containing DDB1 and E27 or an E27 variant, where residues E284-R305 have been replaced by a GNGNSG-linker (E27Δ). SDS-PAGE analyses of GF fractions are shown next to the chromatogram. *-contaminant from E27 preparations.

**(C)** *In vitro* reconstitution, followed by GF analysis, of protein complexes containing STAT2 and E27 or E27Δ. SDS-PAGE analyses of GF fractions are shown as in **D**. ** - contaminant from STAT2 preparation.

**Figure S8: AlphaFold2 prediction of HBx in complex with DDB1**_**ΔBPB**_, **related to Figure 4**

HBx is shown in cartoon representation. DDB1 has been removed for clarity. On the left panel, the model is coloured according to model confidence, i.e., the model’s prediction of its score on the local Distance Difference Test (lDDT-Cα) (Jumper et al., 2021). On the right panel, the model is rainbow-coloured from N-to C-terminus, and side chains involved in zinc coordination (Ramakrishnan et al., 2019) are shown as sticks.

**Figure S9: Side chain densities and mutational analysis of the E27/STAT2 interaction, related to Figure 5**

**(A)** Model-to-map fit of the E27/STAT2 interaction (Fig. 5A). E27 is shown as green cartoon, with selected side chains shown as sticks, and side chains involved in STAT2 interaction marked in light green. STAT2 is shown as magenta cartoon, with selected side chains shown as sticks, and side chains involved in E27 interaction marked in orange. Cryo-EM density map 1 (Fig. S3) is shown as blue mesh, contoured at 4σ, using the the PyMOL Molecular Graphics System, Version 2.0 Schrödinger, LLC.

**(B)** *In vitro* reconstitution, followed by GF analysis, of protein complexes containing STAT2 and E27 or the E27 R211E/R214E double mutant (E27-RE). SDS-PAGE analyses of fractions collected during the GF runs are shown next to the chromatogram. * - contaminant from E27 preparations, ** - contaminant from STAT2 preparations.

**(C)** *In vitro* reconstitution, followed by GF analysis, of protein complexes containing DDB1 and E27 or E27-RE. SDS-PAGE analyses of fractions collected during the GF runs are shown next to the chromatogram as in **B**.

**Figure S10: E27 multiple sequence alignment, cryo-EM map 2 model fit, and CLMS analysis, related to Figure 5**

**(A)** Multiple sequence alignment of E27 and its homologs from other CMV species. >80% type-conserved residues are highlighted black, >60% type-conserved residues grey. Secondary structure elements are drawn schematically above the alignment. Dashed lines indicate flexible regions not included in the molecular model. Residues involved in DDB1-binding and in the main STAT2-binding interface are indicated by blue and magenta dots above the alignment, respectively. Residue stretches involved in the auxiliary E27 α/β-domain/STAT2 DBD binding interface are indicated by magenta lines above the alignment. Side chains involved in zinc-coordination are marked by grey dots.

**(B)** Overview of DDB1_ΔBPB_/E27/STAT2 model fit into cryo-EM map 2 (Fig. S3). DDB1_ΔBPB_ is shown as blue cartoon, E27 as green cartoon, and STAT2 as magenta cartoon. Cryo-EM density is shown as grey semi-transparent surface (threshold level 0.15, Chimerax software tool) (Pettersen et al., 2021).

**(C)** Intermolecular sulfo-SDA cross-links were mapped on the DDB1_ΔBPB_/E27/STAT2 structure (Fig. 5C), colour-coded according to their length as indicated.

**Figure S11: Comparison of E27 and M27 AlphaFold2 predictions, related to Figure 7**

Cartoon representation of AlphaFold2 models of E27 (left) and M27 (right), rainbow-coloured from N-to C-terminus. Terminal regions with very low confidence have been removed. Termini and secondary structure elements are labelled.

**Figure S12: Multiple sequence alignment of human, rat and mouse DDB1, visualisation of non-conserved residues, related to Discussion**

**(A)** Multiple sequence alignment of human (hs), rat (rn) and mouse (mm) DDB1. Non-synonymous but type-conserved amino acid exchanges between the species are highlighted in green, non-conserved exchanges in red. Residues involved in the DDB1_ΔBPB_/E27 interaction interface are highlighted in blue.

**(B)** Amino acid exchanges from (A) have been mapped on a structural model of the DDB1/E27/STAT2 CCD complex. This model has been generated by superposition of a full-length DDB1 structure (PDB 2b5m (Li et al., 2006)) with DDB1_ΔBPB_ from the complex structure presented here (Fig. 3). Only the coordinates of full-length DDB1 are shown for clarity. DDB1 is shown as semi-transparent blue cartoon, with the BPs and the CTD indicated. Non-conserved residues from **A** are shown as spheres using the same colour code as in **A**.

**(C)** Close-up view of the E27-binding cavity as shown in **B**. The closest distance of an amino acid exchange between human and rat DDB1 to an E27 side chain, DDB1 L738T to E27 R290, has been measured as 6.2 Å, more than twice the length of a typical hydrogen bond, strongly indicating that amino acid exchanges between human and rat DDB1 residues do not affect E27 binding in any way.

**Table S1: Cryo-EM data collection and refinement**

**Table S2: Crosslink violation analysis**

**Table S3: Key resources**

**Table S4: Oligonucleotide primer sequences**

## Notes

### Competing Interest Statement

The authors have declared no competing interest.

### Summary of Updates

Figs. 2, 3, 5 and 7 have been revised and re-arranged. The discussion has been extended. The SI has been extended and now includes imporved visualisation and descriptors of structure quality and model-to-map fit.

## References

Andersen, K.R., Leksa, N.C., and Schwartz, T.U. (2013). Optimized E. coli expression strain LOBSTR eliminates common contaminants from His-tag purification. Proteins 81, 1857–1861.

Angers, S., Li, T., Yi, X., MacCoss, M.J., Moon, R.T., and Zheng, N. (2006). Molecular architecture and assembly of the DDB1-CUL4A ubiquitin ligase machinery. Nature 443, 590–593.

Ashour, J., Laurent-Rolle, M., Shi, P.Y., and Garcia-Sastre, A. (2009). NS5 of dengue virus mediates STAT2 binding and degradation. Journal of virology 83, 5408–5418.

Ashour, J., Morrison, J., Laurent-Rolle, M., Belicha-Villanueva, A., Plumlee, C.R., Bernal-Rubio, D., Williams, K.L., Harris, E., Fernandez-Sesma, A., Schindler, C., et al. (2010). Mouse STAT2 restricts early dengue virus replication. Cell host & microbe 8, 410–421.

Baek, K., Krist, D.T., Prabu, J.R., Hill, S., Klugel, M., Neumaier, L.M., von Gronau, S., Kleiger, G., and Schulman, B.A. (2020). NEDD8 nucleates a multivalent cullin-RING-UBE2D ubiquitin ligation assembly. Nature 578, 461–466.

Balfour, H.H., Jr. (1979). Cytomegalovirus: the troll of transplantation. Arch Intern Med 139, 279–280.

Banchenko, S., Krupp, F., Gotthold, C., Burger, J., Graziadei, A., O’Reilly, F.J., Sinn, L., Ruda, O., Rappsilber, J., Spahn, C.M.T., et al. (2021). Structural insights into Cullin4-RING ubiquitin ligase remodelling by Vpr from simian immunodeficiency viruses. PLoS pathogens 17, e1009775.

Baranek, T., Manh, T.P., Alexandre, Y., Maqbool, M.A., Cabeza, J.Z., Tomasello, E., Crozat, K., Bessou, G., Zucchini, N., Robbins, S.H., et al. (2012). Differential responses of immune cells to type I interferon contribute to host resistance to viral infection. Cell host & microbe 12, 571–584.

Barik, S. (2022). Mechanisms of Viral Degradation of Cellular Signal Transducer and Activator of Transcription 2. Int J Mol Sci 23.

Becker, S.D., Bennett, M., Stewart, J.P., and Hurst, J.L. (2007). Serological survey of virus infection among wild house mice (Mus domesticus) in the UK. Lab Anim 41, 229–238.

Becker, T., Le-Trilling, V.T.K., and Trilling, M. (2019). Cellular Cullin RING Ubiquitin Ligases: Druggable Host Dependency Factors of Cytomegaloviruses. Int J Mol Sci 20.

Blaszczyk, K., Olejnik, A., Nowicka, H., Ozgyin, L., Chen, Y.L., Chmielewski, S., Kostyrko, K., Wesoly, J., Balint, B.L., Lee, C.K., et al. (2015). STAT2/IRF9 directs a prolonged ISGF3-like transcriptional response and antiviral activity in the absence of STAT1. Biochem J 466, 511–524.

Brizic, I., Lisnic, B., Brune, W., Hengel, H., and Jonjic, S. (2018). Cytomegalovirus Infection: Mouse Model. Curr Protoc Immunol 122, e51.

Bussiere, D.E., Xie, L., Srinivas, H., Shu, W., Burke, A., Be, C., Zhao, J., Godbole, A., King, D., Karki, R.G., et al. (2020). Structural basis of indisulam-mediated RBM39 recruitment to DCAF15 E3 ligase complex. Nat Chem Biol 16, 15–23.

Cang, Y., Zhang, J., Nicholas, S.A., Bastien, J., Li, B., Zhou, P., and Goff, S.P. (2006). Deletion of DDB1 in mouse brain and lens leads to p53-dependent elimination of proliferating cells. Cell 127, 929–940.

Cavadini, S., Fischer, E.S., Bunker, R.D., Potenza, A., Lingaraju, G.M., Goldie, K.N., Mohamed, W.I., Faty, M., Petzold, G., Beckwith, R.E., et al. (2016). Cullin-RING ubiquitin E3 ligase regulation by the COP9 signalosome. Nature 531, 598–603.

Chamberlain, P.P., Lopez-Girona, A., Miller, K., Carmel, G., Pagarigan, B., Chie-Leon, B., Rychak, E., Corral, L.G., Ren, Y.J., Wang, M., et al. (2014). Structure of the human Cereblon-DDB1-lenalidomide complex reveals basis for responsiveness to thalidomide analogs. Nature structural & molecular biology 21, 803–809.

Darnell, J.E., Jr., Kerr, I.M., and Stark, G.R. (1994). Jak-STAT pathways and transcriptional activation in response to IFNs and other extracellular signaling proteins. Science 264, 1415–1421.

Decorsiere, A., Mueller, H., van Breugel, P.C., Abdul, F., Gerossier, L., Beran, R.K., Livingston, C.M., Niu, C., Fletcher, S.P., Hantz, O., et al. (2016). Hepatitis B virus X protein identifies the Smc5/6 complex as a host restriction factor. Nature 531, 386–389.

Deutsch, E.W., Bandeira, N., Sharma, V., Perez-Riverol, Y., Carver, J.J., Kundu, D.J., Garcia-Seisdedos, D., Jarnuczak, A.F., Hewapathirana, S., Pullman, B.S., et al. (2020). The ProteomeXchange consortium in 2020: enabling ‘big data’ approaches in proteomics. Nucleic acids research 48, D1145–D1152.

Drulyte, I., Johnson, R.M., Hesketh, E.L., Hurdiss, D.L., Scarff, C.A., Porav, S.A., Ranson, N.A., Muench, S.P., and Thompson, R.F. (2018). Approaches to altering particle distributions in cryo-electron microscopy sample preparation. Acta Crystallogr D Struct Biol 74, 560–571.

Du, X., Volkov, O.A., Czerwinski, R.M., Tan, H., Huerta, C., Morton, E.R., Rizzi, J.P., Wehn, P.M., Xu, R., Nijhawan, D., et al. (2019). Structural Basis and Kinetic Pathway of RBM39 Recruitment to DCAF15 by a Sulfonamide Molecular Glue E7820. Structure 27, 1625–1633 e1623.

Emsley, P., and Cowtan, K. (2004). Coot: model-building tools for molecular graphics. Acta crystallographica Section D, Biological crystallography 60, 2126–2132.

Emsley, P., Lohkamp, B., Scott, W.G., and Cowtan, K. (2010). Features and development of Coot. Acta crystallographica Section D, Biological crystallography 66, 486–501.

Ettinger, J., Geyer, H., Nitsche, A., Zimmermann, A., Brune, W., Sandford, G.R., Hayward, G.S., and Voigt, S. (2012). Complete genome sequence of the english isolate of rat cytomegalovirus (Murid herpesvirus 8). Journal of virology 86, 13838.

Faust, T.B., Yoon, H., Nowak, R.P., Donovan, K.A., Li, Z., Cai, Q., Eleuteri, N.A., Zhang, T., Gray, N.S., and Fischer, E.S. (2020). Structural complementarity facilitates E7820-mediated degradation of RBM39 by DCAF15. Nat Chem Biol 16, 7–14.

Fischer, E.S., Bohm, K., Lydeard, J.R., Yang, H., Stadler, M.B., Cavadini, S., Nagel, J., Serluca, F., Acker, V., Lingaraju, G.M., et al. (2014). Structure of the DDB1-CRBN E3 ubiquitin ligase in complex with thalidomide. Nature 512, 49–53.

Fischer, E.S., Scrima, A., Bohm, K., Matsumoto, S., Lingaraju, G.M., Faty, M., Yasuda, T., Cavadini, S., Wakasugi, M., Hanaoka, F., et al. (2011). The molecular basis of CRL4DDB2/CSA ubiquitin ligase architecture, targeting, and activation. Cell 147, 1024–1039.

Fischer, U., Hertlein, S., and Grimm, C. (2015). The structure of apo ArnA features an unexpected central binding pocket and provides an explanation for enzymatic cooperativity. Acta crystallographica Section D, Biological crystallography 71, 687–696.

Geyer, H., Ettinger, J., Moller, L., Schmolz, E., Nitsche, A., Brune, W., Heaggans, S., Sandford, G.R., Hayward, G.S., and Voigt, S. (2015). Rat cytomegalovirus (RCMV) English isolate and a newly identified Berlin isolate share similarities with but are separate as an anciently diverged clade from Mouse CMV and the Maastricht isolate of RCMV. The Journal of general virology 96, 1873–1870.

Goldenberg, S.J., Cascio, T.C., Shumway, S.D., Garbutt, K.C., Liu, J., Xiong, Y., and Zheng, N. (2004). Structure of the Cand1-Cul1-Roc1 complex reveals regulatory mechanisms for the assembly of the multisubunit cullin-dependent ubiquitin ligases. Cell 119, 517–528.

Gowen, B.B., Westover, J.B., Miao, J., Van Wettere, A.J., Rigas, J.D., Hickerson, B.T., Jung, K.H., Li, R., Conrad, B.L., Nielson, S., et al. (2017). Modeling Severe Fever with Thrombocytopenia Syndrome Virus Infection in Golden Syrian Hamsters: Importance of STAT2 in Preventing Disease and Effective Treatment with Favipiravir. Journal of virology 91.

Grant, A., Ponia, S.S., Tripathi, S., Balasubramaniam, V., Miorin, L., Sourisseau, M., Schwarz, M.C., Sanchez-Seco, M.P., Evans, M.J., Best, S.M., et al. (2016). Zika Virus Targets Human STAT2 to Inhibit Type I Interferon Signaling. Cell host & microbe 19, 882–890.

Greenwood, E.J.D., Williamson, J.C., Sienkiewicz, A., Naamati, A., Matheson, N.J., and Lehner, P.J. (2019). Promiscuous Targeting of Cellular Proteins by Vpr Drives Systems-Level Proteomic Remodeling in HIV-1 Infection. Cell reports 27, 1579–1596 e1577.

Griffiths, P., and Reeves, M. (2021). Pathogenesis of human cytomegalovirus in the immunocompromised host. Nat Rev Microbiol 19, 759–773.

Hambleton, S., Goodbourn, S., Young, D.F., Dickinson, P., Mohamad, S.M., Valappil, M., McGovern, N., Cant, A.J., Hackett, S.J., Ghazal, P., et al. (2013). STAT2 deficiency and susceptibility to viral illness in humans. Proceedings of the National Academy of Sciences of the United States of America 110, 3053–3058.

Hansson, M.D., Rzeznicka, K., Rosenback, M., Hansson, M., and Sirijovski, N. (2008). PCR-mediated deletion of plasmid DNA. Anal Biochem 375, 373–375.

Harper, J.W., and Schulman, B.A. (2021). Cullin-RING Ubiquitin Ligase Regulatory Circuits: A Quarter Century Beyond the F-Box Hypothesis. Annual review of biochemistry 90, 403–429.

Hein, M.Y., and Weissman, J.S. (2021). Functional single-cell genomics of human cytomegalovirus infection. bioRxiv, 775080.

Ho, J., Pelzel, C., Begitt, A., Mee, M., Elsheikha, H.M., Scott, D.J., and Vinkemeier, U. (2016). STAT2 Is a Pervasive Cytokine Regulator due to Its Inhibition of STAT1 in Multiple Signaling Pathways. PLoS Biol 14, e2000117.

Horvath, C.M., Stark, G.R., Kerr, I.M., and Darnell, J.E., Jr. (1996). Interactions between STAT and non-STAT proteins in the interferon-stimulated gene factor 3 transcription complex. Molecular and cellular biology 16, 6957–6964.

Humphreys, I.R., Pei, J., Baek, M., Krishnakumar, A., Anishchenko, I., Ovchinnikov, S., Zhang, J., Ness, T.J., Banjade, S., Bagde, S.R., et al. (2021). Computed structures of core eukaryotic protein complexes. Science 374, eabm4805.

Ireland, S.M., and Martin, A.C.R. (2019). ZincBind-the database of zinc binding sites. Database (Oxford) 2019.

Isaacs, A., and Lindenmann, J. (1957). Virus interference. I. The interferon. Proc R Soc Lond B Biol Sci 147, 258–267.

Ivashkiv, L.B., and Donlin, L.T. (2014). Regulation of type I interferon responses. Nat Rev Immunol 14, 36–49.

Jones, M., Davidson, A., Hibbert, L., Gruenwald, P., Schlaak, J., Ball, S., Foster, G.R., and Jacobs, M. (2005). Dengue virus inhibits alpha interferon signaling by reducing STAT2 expression. Journal of virology 79, 5414–5420.

Jordan, S., Krause, J., Prager, A., Mitrovic, M., Jonjic, S., Koszinowski, U.H., and Adler, B. (2011). Virus progeny of murine cytomegalovirus bacterial artificial chromosome pSM3fr show reduced growth in salivary Glands due to a fixed mutation of MCK-2. Journal of virology 85, 10346–10353.

Jumper, J., Evans, R., Pritzel, A., Green, T., Figurnov, M., Ronneberger, O., Tunyasuvunakool, K., Bates, R., Zidek, A., Potapenko, A., et al. (2021). Highly accurate protein structure prediction with AlphaFold. Nature 596, 583–589.

Kidmose, R.T., Juhl, J., Nissen, P., Boesen, T., Karlsen, J.L., and Pedersen, B.P. (2019). Namdinator - automatic molecular dynamics flexible fitting of structural models into cryo-EM and crystallography experimental maps. IUCrJ 6, 526–531.

Kotenko, S.V., Gallagher, G., Baurin, V.V., Lewis-Antes, A., Shen, M., Shah, N.K., Langer, J.A., Sheikh, F., Dickensheets, H., and Donnelly, R.P. (2003). IFN-lambdas mediate antiviral protection through a distinct class II cytokine receptor complex. Nature immunology 4, 69–77.

Laitaoja, M., Valjakka, J., and Janis, J. (2013). Zinc coordination spheres in protein structures. Inorg Chem 52, 10983–10991.

Landsberg, C.D., Megger, D.A., Hotter, D., Ruckborn, M.U., Eilbrecht, M., Rashidi-Alavijeh, J., Howe, S., Heinrichs, S., Sauter, D., Sitek, B., et al. (2018). A Mass Spectrometry-Based Profiling of Interactomes of Viral DDB1- and Cullin Ubiquitin Ligase-Binding Proteins Reveals NF-kappaB Inhibitory Activity of the HIV-2-Encoded Vpx. Front Immunol 9, 2978.

Le-Trilling, V.T., Megger, D.A., Katschinski, B., Landsberg, C.D., Ruckborn, M.U., Tao, S., Krawczyk, A., Bayer, W., Drexler, I., Tenbusch, M., et al. (2016). Broad and potent antiviral activity of the NAE inhibitor MLN4924. Scientific reports 6, 19977.

Le-Trilling, V.T., and Trilling, M. (2017). Mouse newborn cells allow highly productive mouse cytomegalovirus replication, constituting a novel convenient primary cell culture system. PloS one 12, e0174695.

Le-Trilling, V.T.K., Becker, T., Nachshon, A., Stern-Ginossar, N., Scholer, L., Voigt, S., Hengel, H., and Trilling, M. (2020). The Human Cytomegalovirus pUL145 Isoforms Act as Viral DDB1-Cullin-Associated Factors to Instruct Host Protein Degradation to Impede Innate Immunity. Cell reports 30, 2248–2260 e2245.

Le-Trilling, V.T.K., and Trilling, M. (2020). Ub to no good: How cytomegaloviruses exploit the ubiquitin proteasome system. Virus Res 281, 197938.

Le-Trilling, V.T.K., Wohlgemuth, K., Ruckborn, M.U., Jagnjic, A., Maassen, F., Timmer, L., Katschinski, B., and Trilling, M. (2018). STAT2-Dependent Immune Responses Ensure Host Survival despite the Presence of a Potent Viral Antagonist. Journal of virology 92.

Le, V.T., Trilling, M., and Hengel, H. (2011). The cytomegaloviral protein pUL138 acts as potentiator of tumor necrosis factor (TNF) receptor 1 surface density to enhance ULb’-encoded modulation of TNF-alpha signaling. Journal of virology 85, 13260–13270.

Le, V.T.K., Trilling, M., Wilborn, M., Hengel, H., and Zimmermann, A. (2008). Human cytomegalovirus interferes with signal transducer and activator of transcription (STAT) 2 protein stability and tyrosine phosphorylation. The Journal of general virology 89, 2416–2426.

Levy, D.E., and Darnell, J.E., Jr. (2002). Stats: transcriptional control and biological impact. Nature reviews Molecular cell biology 3, 651–662.

Li, T., Chen, X., Garbutt, K.C., Zhou, P., and Zheng, N. (2006). Structure of DDB1 in complex with a paramyxovirus V protein: viral hijack of a propeller cluster in ubiquitin ligase. Cell 124, 105–117.

Li, T., Robert, E.I., van Breugel, P.C., Strubin, M., and Zheng, N. (2010). A promiscuous alpha-helical motif anchors viral hijackers and substrate receptors to the CUL4-DDB1 ubiquitin ligase machinery. Nature structural & molecular biology 17, 105–111.

Liebschner, D., Afonine, P.V., Baker, M.L., Bunkoczi, G., Chen, V.B., Croll, T.I., Hintze, B., Hung, L.W., Jain, S., McCoy, A.J., et al. (2019). Macromolecular structure determination using X-rays, neutrons and electrons: recent developments in Phenix. Acta Crystallogr D Struct Biol 75, 861–877.

Lin, G.Y., Paterson, R.G., Richardson, C.D., and Lamb, R.A. (1998). The V protein of the paramyxovirus SV5 interacts with damage-specific DNA binding protein. Virology 249, 189–200.

Madeira, F., Park, Y.M., Lee, J., Buso, N., Gur, T., Madhusoodanan, N., Basutkar, P., Tivey, A.R.N., Potter, S.C., Finn, R.D., et al. (2019). The EMBL-EBI search and sequence analysis tools APIs in 2019. Nucleic acids research 47, W636–W641.

Matsumoto, M., Tanaka, N., Harada, H., Kimura, T., Yokochi, T., Kitagawa, M., Schindler, C., and Taniguchi, T. (1999). Activation of the transcription factor ISGF3 by interferon-gamma. Biol Chem 380, 699–703.

Megger, D.A., Philipp, J., Le-Trilling, V.T.K., Sitek, B., and Trilling, M. (2017). Deciphering of the Human Interferon-Regulated Proteome by Mass Spectrometry-Based Quantitative Analysis Reveals Extent and Dynamics of Protein Induction and Repression. Front Immunol 8, 1139.

Mindell, J.A., and Grigorieff, N. (2003). Accurate determination of local defocus and specimen tilt in electron microscopy. J Struct Biol 142, 334–347.

Mirdita, M., Ovchinnikov, S., and Steinegger, M. (2021). ColabFold - Making protein folding accessible to all. bioRxiv, 2021.2008.2015.456425.

Nicholas, K.B., Nicholas Jr., H. B., Deerfield II., D. W. (1997). GeneDoc: Analysis and Visualization of Genetic Variation. embnetnews 4, 1–4.

Nightingale, K., Lin, K.M., Ravenhill, B.J., Davies, C., Nobre, L., Fielding, C.A., Ruckova, E., Fletcher-Etherington, A., Soday, L., Nichols, H., et al. (2018). High-Definition Analysis of Host Protein Stability during Human Cytomegalovirus Infection Reveals Antiviral Factors and Viral Evasion Mechanisms. Cell host & microbe 24, 447–460 e411.

Nightingale, K., Potts, M., Hunter, L.M., Fielding, C.A., Zerbe, C.M., Fletcher-Etherington, A., Nobre, L., Wang, E.C.Y., Strang, B.L., Houghton, J.W., et al. (2022). Human cytomegalovirus protein RL1 degrades the antiviral factor SLFN11 via recruitment of the CRL4 E3 ubiquitin ligase complex. Proceedings of the National Academy of Sciences of the United States of America 119.

Nobre, L.V., Nightingale, K., Ravenhill, B.J., Antrobus, R., Soday, L., Nichols, J., Davies, J.A., Seirafian, S., Wang, E.C., Davison, A.J., et al. (2019). Human cytomegalovirus interactome analysis identifies degradation hubs, domain associations and viral protein functions. Elife 8.

O’Reilly, F.J., and Rappsilber, J. (2018). Cross-linking mass spectrometry: methods and applications in structural, molecular and systems biology. Nature structural & molecular biology 25, 1000–1008.

Olma, M.H., Roy, M., Le Bihan, T., Sumara, I., Maerki, S., Larsen, B., Quadroni, M., Peter, M., Tyers, M., and Pintard, L. (2009). An interaction network of the mammalian COP9 signalosome identifies Dda1 as a core subunit of multiple Cul4-based E3 ligases. Journal of cell science 122, 1035–1044.

Pace, C.N., and Scholtz, J.M. (1998). A helix propensity scale based on experimental studies of peptides and proteins. Biophys J 75, 422–427.

Panne, D., Maniatis, T., and Harrison, S.C. (2007). An atomic model of the interferon-beta enhanceosome. Cell 129, 1111–1123.

Parisien, J.P., Lau, J.F., and Horvath, C.M. (2002a). STAT2 acts as a host range determinant for species-specific paramyxovirus interferon antagonism and simian virus 5 replication. Journal of virology 76, 6435–6441.

Parisien, J.P., Lau, J.F., Rodriguez, J.J., Sullivan, B.M., Moscona, A., Parks, G.D., Lamb, R.A., and Horvath, C.M. (2001). The V protein of human parainfluenza virus 2 antagonizes type I interferon responses by destabilizing signal transducer and activator of transcription 2. Virology 283, 230–239.

Parisien, J.P., Lau, J.F., Rodriguez, J.J., Ulane, C.M., and Horvath, C.M. (2002b). Selective STAT protein degradation induced by paramyxoviruses requires both STAT1 and STAT2 but is independent of alpha/beta interferon signal transduction. Journal of virology 76, 4190–4198.

Park, C., Li, S., Cha, E., and Schindler, C. (2000). Immune response in Stat2 knockout mice. Immunity 13, 795–804.

Perez-Riverol, Y., Csordas, A., Bai, J., Bernal-Llinares, M., Hewapathirana, S., Kundu, D.J., Inuganti, A., Griss, J., Mayer, G., Eisenacher, M., et al. (2019). The PRIDE database and related tools and resources in 2019: improving support for quantification data. Nucleic acids research 47, D442–D450.

Pettersen, E.F., Goddard, T.D., Huang, C.C., Meng, E.C., Couch, G.S., Croll, T.I., Morris, J.H., and Ferrin, T.E. (2021). UCSF ChimeraX: Structure visualization for researchers, educators, and developers. Protein Sci 30, 70–82.

Petzold, G., Fischer, E.S., and Thoma, N.H. (2016). Structural basis of lenalidomide-induced CK1alpha degradation by the CRL4(CRBN) ubiquitin ligase. Nature 532, 127–130.

Piersimoni, L., Kastritis, P.L., Arlt, C., and Sinz, A. (2022). Cross-Linking Mass Spectrometry for Investigating Protein Conformations and Protein-Protein Interactions horizontal line A Method for All Seasons. Chem Rev 122, 7500–7531.

Platanitis, E., Demiroz, D., Schneller, A., Fischer, K., Capelle, C., Hartl, M., Gossenreiter, T., Muller, M., Novatchkova, M., and Decker, T. (2019). A molecular switch from STAT2-IRF9 to ISGF3 underlies interferon-induced gene transcription. Nature communications 10, 2921.

Precious, B., Childs, K., Fitzpatrick-Swallow, V., Goodbourn, S., and Randall, R.E. (2005). Simian virus 5 V protein acts as an adaptor, linking DDB1 to STAT2, to facilitate the ubiquitination of STAT1. Journal of virology 79, 13434–13441.

Punjani, A., and Fleet, D.J. (2021). 3D variability analysis: Resolving continuous flexibility and discrete heterogeneity from single particle cryo-EM. J Struct Biol 213, 107702.

Punjani, A., Rubinstein, J.L., Fleet, D.J., and Brubaker, M.A. (2017). cryoSPARC: algorithms for rapid unsupervised cryo-EM structure determination. Nat Methods 14, 290–296.

Punjani, A., Zhang, H., and Fleet, D.J. (2020). Non-uniform refinement: adaptive regularization improves single-particle cryo-EM reconstruction. Nat Methods 17, 1214–1221.

Qureshi, S.A., Salditt-Georgieff, M., and Darnell, J.E., Jr. (1995). Tyrosine-phosphorylated Stat1 and Stat2 plus a 48-kDa protein all contact DNA in forming interferon-stimulated-gene factor 3. Proceedings of the National Academy of Sciences of the United States of America 92, 3829–3833.

Ramakrishnan, D., Xing, W., Beran, R.K., Chemuru, S., Rohrs, H., Niedziela-Majka, A., Marchand, B., Mehra, U., Zabransky, A., Dolezal, M., et al. (2019). Hepatitis B Virus X Protein Function Requires Zinc Binding. Journal of virology 93.

Rappsilber, J., Ishihama, Y., and Mann, M. (2003). Stop and go extraction tips for matrix-assisted laser desorption/ionization, nanoelectrospray, and LC/MS sample pretreatment in proteomics. Anal Chem 75, 663–670.

Rattay, S., Trilling, M., Megger, D.A., Sitek, B., Meyer, H.E., Hengel, H., and Le-Trilling, V.T. (2015). The Canonical Immediate Early 3 Gene Product pIE611 of Mouse Cytomegalovirus Is Dispensable for Viral Replication but Mediates Transcriptional and Posttranscriptional Regulation of Viral Gene Products. Journal of virology 89, 8590–8598.

Reddehase, M.J., and Lemmermann, N.A.W. (2018). Mouse Model of Cytomegalovirus Disease and Immunotherapy in the Immunocompromised Host: Predictions for Medical Translation that Survived the “Test of Time”. Viruses 10.

Reichermeier, K.M., Straube, R., Reitsma, J.M., Sweredoski, M.J., Rose, C.M., Moradian, A., den Besten, W., Hinkle, T., Verschueren, E., Petzold, G., et al. (2020). PIKES Analysis Reveals Response to Degraders and Key Regulatory Mechanisms of the CRL4 Network. Molecular cell 77, 1092–1106 e1099.

Reitsma, J.M., Savaryn, J.P., Faust, K., Sato, H., Halligan, B.D., and Terhune, S.S. (2011). Antiviral inhibition targeting the HCMV kinase pUL97 requires pUL27-dependent degradation of Tip60 acetyltransferase and cell-cycle arrest. Cell host & microbe 9, 103–114.

Rengachari, S., Groiss, S., Devos, J.M., Caron, E., Grandvaux, N., and Panne, D. (2018). Structural basis of STAT2 recognition by IRF9 reveals molecular insights into ISGF3 function. Proceedings of the National Academy of Sciences of the United States of America 115, E601–E609.

Salsman, J., Jagannathan, M., Paladino, P., Chan, P.K., Dellaire, G., Raught, B., and Frappier, L. (2012). Proteomic profiling of the human cytomegalovirus UL35 gene products reveals a role for UL35 in the DNA repair response. Journal of virology 86, 806–820.

Scheres, S.H. (2012). RELION: implementation of a Bayesian approach to cryo-EM structure determination. J Struct Biol 180, 519–530.

Schneider, K., Loewendorf, A., De Trez, C., Fulton, J., Rhode, A., Shumway, H., Ha, S., Patterson, G., Pfeffer, K., Nedospasov, S.A., et al. (2008). Lymphotoxin-mediated crosstalk between B cells and splenic stroma promotes the initial type I interferon response to cytomegalovirus. Cell host & microbe 3, 67–76.

Schneider, M., Belsom, A., and Rappsilber, J. (2018). Protein Tertiary Structure by Crosslinking/Mass Spectrometry. Trends in biochemical sciences 43, 157–169.

Schwefel, D., Boucherit, V.C., Christodoulou, E., Walker, P.A., Stoye, J.P., Bishop, K.N., and Taylor, I.A. (2015). Molecular Determinants for Recognition of Divergent SAMHD1 Proteins by the Lentiviral Accessory Protein Vpx. Cell host & microbe 17, 489–499.

Schwefel, D., Groom, H.C., Boucherit, V.C., Christodoulou, E., Walker, P.A., Stoye, J.P., Bishop, K.N., and Taylor, I.A. (2014). Structural basis of lentiviral subversion of a cellular protein degradation pathway. Nature 505, 234–238.

Shabek, N., Ruble, J., Waston, C.J., Garbutt, K.C., Hinds, T.R., Li, T., and Zheng, N. (2018). Structural insights into DDA1 function as a core component of the CRL4-DDB1 ubiquitin ligase. Cell Discov 4, 67.

Shevchenko, A., Tomas, H., Havlis, J., Olsen, J.V., and Mann, M. (2006). In-gel digestion for mass spectrometric characterization of proteins and proteomes. Nature protocols 1, 2856–2860.

Slabicki, M., Kozicka, Z., Petzold, G., Li, Y.D., Manojkumar, M., Bunker, R.D., Donovan, K.A., Sievers, Q.L., Koeppel, J., Suchyta, D., et al. (2020). The CDK inhibitor CR8 acts as a molecular glue degrader that depletes cyclin K. Nature 585, 293–297.

Stark, H. (2010). GraFix: stabilization of fragile macromolecular complexes for single particle cryo-EM. Methods Enzymol 481, 109–126.

Suloway, C., Pulokas, J., Fellmann, D., Cheng, A., Guerra, F., Quispe, J., Stagg, S., Potter, C.S., and Carragher, B. (2005). Automated molecular microscopy: the new Leginon system. J Struct Biol 151, 41–60.

Tang, J., and Chu, G. (2002). Xeroderma pigmentosum complementation group E and UV-damaged DNA-binding protein. DNA Repair (Amst) 1, 601–616.

Tischer, B.K., von Einem, J., Kaufer, B., and Osterrieder, N. (2006). Two-step red-mediated recombination for versatile high-efficiency markerless DNA manipulation in Escherichia coli. Biotechniques 40, 191–197.

Trilling, M., Bellora, N., Rutkowski, A.J., de Graaf, M., Dickinson, P., Robertson, K., Prazeres da Costa, O., Ghazal, P., Friedel, C.C., Alba, M.M., et al. (2013). Deciphering the modulation of gene expression by type I and II interferons combining 4sU-tagging, translational arrest and in silico promoter analysis. Nucleic acids research 41, 8107–8125.

Trilling, M., Le, V.T., Fiedler, M., Zimmermann, A., Bleifuss, E., and Hengel, H. (2011). Identification of DNA-damage DNA-binding protein 1 as a conditional essential factor for cytomegalovirus replication in interferon-gamma-stimulated cells. PLoS pathogens 7, e1002069.

Trilling, M., Le, V.T., Zimmermann, A., Ludwig, H., Pfeffer, K., Sutter, G., Smith, G.L., and Hengel, H. (2009). Gamma interferon-induced interferon regulatory factor 1-dependent antiviral response inhibits vaccinia virus replication in mouse but not human fibroblasts. Journal of virology 83, 3684–3695.

Ulane, C.M., and Horvath, C.M. (2002). Paramyxoviruses SV5 and HPIV2 assemble STAT protein ubiquitin ligase complexes from cellular components. Virology 304, 160–166.

Ulane, C.M., Kentsis, A., Cruz, C.D., Parisien, J.P., Schneider, K.L., and Horvath, C.M. (2005). Composition and assembly of STAT-targeting ubiquitin ligase complexes: paramyxovirus V protein carboxyl terminus is an oligomerization domain. Journal of virology 79, 10180–10189.

van Diemen, F.R., Kruse, E.M., Hooykaas, M.J., Bruggeling, C.E., Schurch, A.C., van Ham, P.M., Imhof, S.M., Nijhuis, M., Wiertz, E.J., and Lebbink, R.J. (2016). CRISPR/Cas9-Mediated Genome Editing of Herpesviruses Limits Productive and Latent Infections. PLoS pathogens 12, e1005701.

van Zundert, G.C., and Bonvin, A.M. (2015). DisVis: quantifying and visualizing accessible interaction space of distance-restrained biomolecular complexes. Bioinformatics 31, 3222–3224.

van Zundert, G.C., Trellet, M., Schaarschmidt, J., Kurkcuoglu, Z., David, M., Verlato, M., Rosato, A., and Bonvin, A.M. (2017). The DisVis and PowerFit Web Servers: Explorative and Integrative Modeling of Biomolecular Complexes. Journal of molecular biology 429, 399–407.

Varadi, M., Anyango, S., Deshpande, M., Nair, S., Natassia, C., Yordanova, G., Yuan, D., Stroe, O., Wood, G., Laydon, A., et al. (2022). AlphaFold Protein Structure Database: massively expanding the structural coverage of protein-sequence space with high-accuracy models. Nucleic acids research 50, D439–D444.

Virgin, H.W., Wherry, E.J., and Ahmed, R. (2009). Redefining chronic viral infection. Cell 138, 30–50.

Voigt, S., Mesci, A., Ettinger, J., Fine, J.H., Chen, P., Chou, W., and Carlyle, J.R. (2007). Cytomegalovirus evasion of innate immunity by subversion of the NKR-P1B:Clr-b missing-self axis. Immunity 26, 617–627.

Wagner, M., Ruzsics, Z., and Koszinowski, U.H. (2002). Herpesvirus genetics has come of age. Trends Microbiol 10, 318–324.

Wakasugi, M., Matsuura, K., Nagasawa, A., Fu, D., Shimizu, H., Yamamoto, K., Takeda, S., and Matsunaga, T. (2007). DDB1 gene disruption causes a severe growth defect and apoptosis in chicken DT40 cells. Biochem Biophys Res Commun 364, 771–777.

Wang, H., Guo, H., Su, J., Rui, Y., Zheng, W., Gao, W., Zhang, W., Li, Z., Liu, G., Markham, R.B., et al. (2017). Inhibition of Vpx-Mediated SAMHD1 and Vpr-Mediated Host Helicase Transcription Factor Degradation by Selective Disruption of Viral CRL4 (DCAF1) E3 Ubiquitin Ligase Assembly. Journal of virology 91.

Wick, E.T., Treadway, C.J., Li, Z., Nicely, N.I., Ren, Z., Baldwin, A.S., Xiong, Y., Harrison, J.S., and Brown, N.G. (2022). Insight into Viral Hijacking of CRL4 Ubiquitin Ligase through Structural Analysis of the pUL145-DDB1 Complex. Journal of virology, e0082622.

Wilkins, M.R., Gasteiger, E., Bairoch, A., Sanchez, J.C., Williams, K.L., Appel, R.D., and Hochstrasser, D.F. (1999). Protein identification and analysis tools in the ExPASy server. Methods Mol Biol 112, 531–552.

Wu, Y., Koharudin, L.M., Mehrens, J., DeLucia, M., Byeon, C.H., Byeon, I.J., Calero, G., Ahn, J., and Gronenborn, A.M. (2015). Structural Basis of Clade-specific Engagement of SAMHD1 (Sterile alpha Motif and Histidine/Aspartate-containing Protein 1) Restriction Factors by Lentiviral Viral Protein X (Vpx) Virulence Factors. The Journal of biological chemistry 290, 17935–17945.

Yamamoto, M., Koga, R., Fujino, H., Shimagaki, K., Ciftci, H.I., Kamo, M., Tateishi, H., Otsuka, M., and Fujita, M. (2017). Zinc-binding site of human immunodeficiency virus 2 Vpx prevents instability and dysfunction of the protein. The Journal of general virology 98, 275–283.

Yang, M., Chen, Y.S., Ichikawa, M., Calles-Garcia, D., Basu, K., Fakih, R., Bui, K.H., and Gehring, K. (2019). Cryo-electron microscopy structures of ArnA, a key enzyme for polymyxin resistance, revealed unexpected oligomerizations and domain movements. J Struct Biol 208, 43–50.

Yoshikawa, R., Sakabe, S., Urata, S., and Yasuda, J. (2019). Species-Specific Pathogenicity of Severe Fever with Thrombocytopenia Syndrome Virus Is Determined by Anti-STAT2 Activity of NSs. Journal of virology 93.

Zhao, Y., Chapman, D.A., and Jones, I.M. (2003). Improving baculovirus recombination. Nucleic acids research 31, E6–6.

Zheng, S.Q., Palovcak, E., Armache, J.P., Verba, K.A., Cheng, Y., and Agard, D.A. (2017). MotionCor2: anisotropic correction of beam-induced motion for improved cryo-electron microscopy. Nat Methods 14, 331–332.

Zimmerman, E.S., Schulman, B.A., and Zheng, N. (2010). Structural assembly of cullin-RING ubiquitin ligase complexes. Current opinion in structural biology 20, 714–721.

Zimmermann, A., Trilling, M., Wagner, M., Wilborn, M., Bubic, I., Jonjic, S., Koszinowski, U., and Hengel, H. (2005). A cytomegaloviral protein reveals a dual role for STAT2 in IFN-{gamma} signaling and antiviral responses. The Journal of experimental medicine 201, 1543–1553.

Zivanov, J., Nakane, T., Forsberg, B.O., Kimanius, D., Hagen, W.J., Lindahl, E., and Scheres, S.H. (2018). New tools for automated high-resolution cryo-EM structure determination in RELION-3. Elife 7.

