## Supplementary Information for "Structure and mechanism of a novel cytomegaloviral DCAF mediating interferon antagonism"

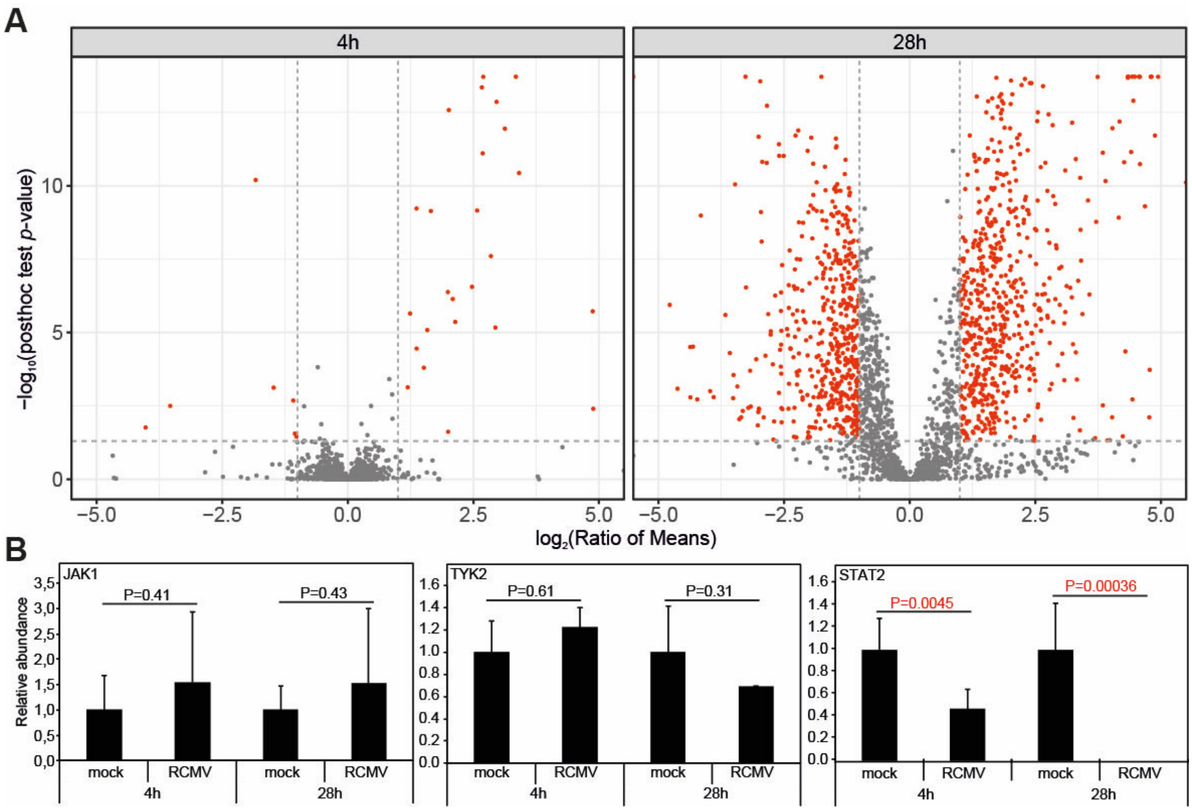


**Figure S1: Proteomic analyses of protein abundance changes upon RCMV-E infection, related to Figure 1**

**(A)** Mass spectrometric analyses of protein abundance changes upon RCMV-E infection, compared to mock-infected samples, at indicated time points. Significantly down- or upregulated proteins are marked in red.

**(B)** Relative abundance changes of selected proteins involved in IFN signalling upon RCMV-E infection compared to mock-infected samples at indicated time points.

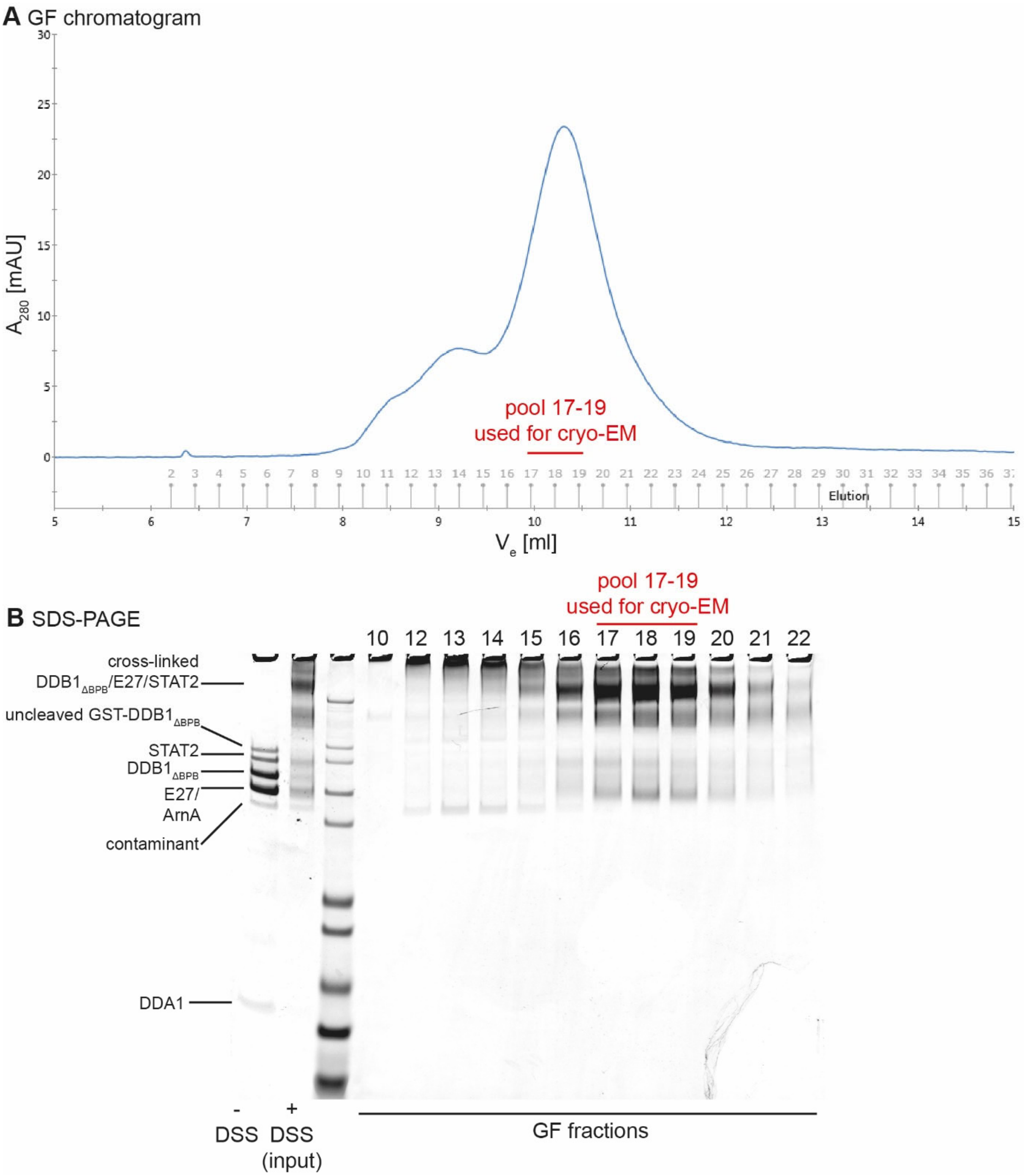


**Figure S2: Cryo-EM sample preparation, related to Figures 3-5**

**(A)** GF chromatogram of the disuccinimidyl suberate (DSS) cross-linked DDB1$_{\Delta BPB}$/DDA1/E27/STAT2 protein complex. GF fractions that have been pooled for cryo-EM analysis are indicated.

**(B)** SDS-PAGE analysis of the cross-linking reaction and GF fractions.

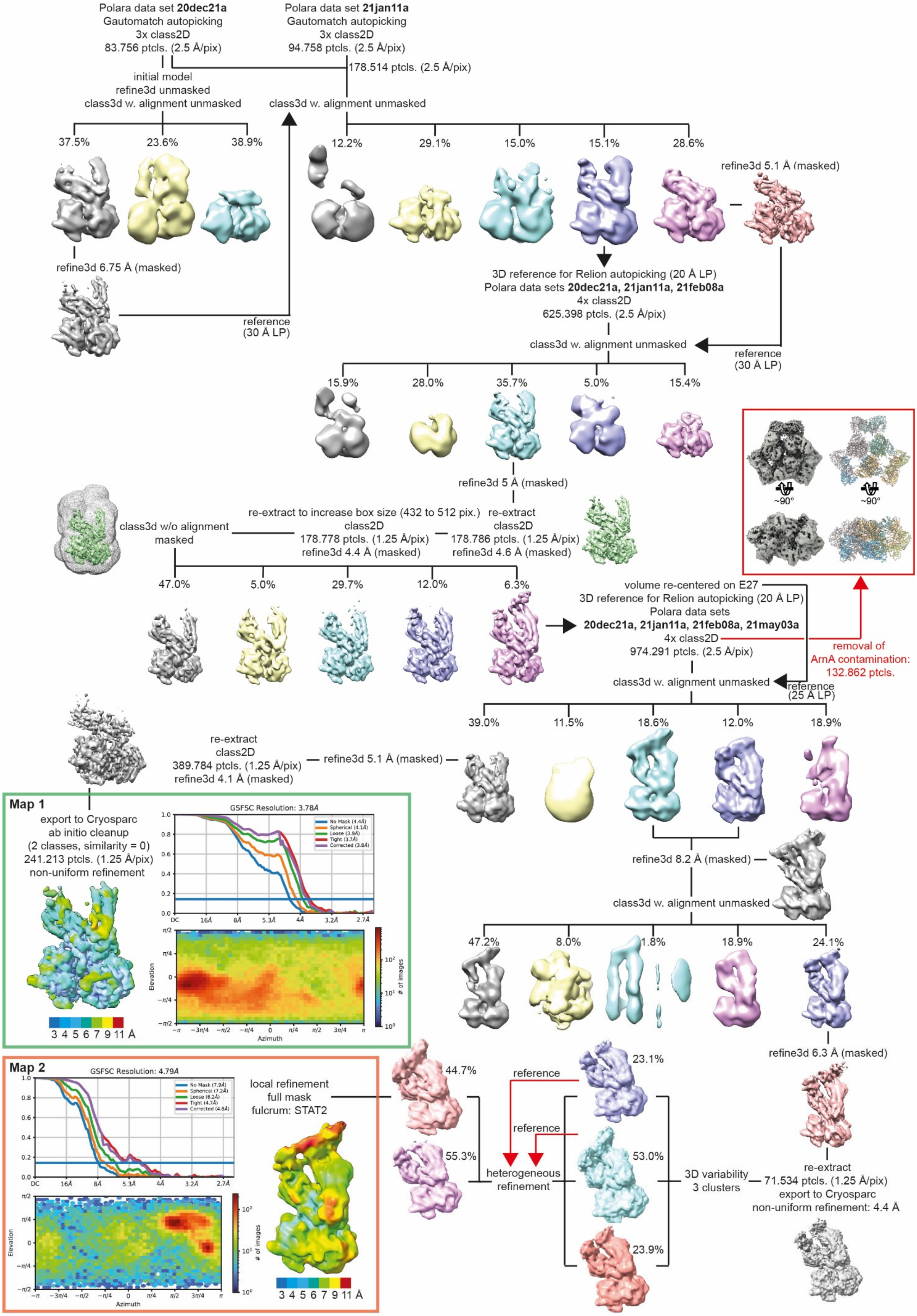

Polara data set 20dec21a
Gautomatch autopicking
3x class2D
83.756 ptcls. (2.5 Å/pix)
Polara data set 21jan11a
Gautomatch autopicking
3x class2D
94.758 ptcls. (2.5 Å/pix)
178.514 ptcls. (2.5 Å/pix)
initial model
refine3d unmasked
class3d w. alignment unmasked
class3d w. alignment unmasked
37.5%
23.6%
38.9%
12.2%
29.1%
15.0%
15.1%
28.6%
refine3d 5.1 Å (masked)
refine3d 6.75 Å (masked)
reference
(30 Å LP)
3D reference for Relion autopicking (20 Å LP)
Polara data sets 20dec21a, 21jan11a, 21feb08a
4x class2D
625.398 ptcls. (2.5 Å/pix)
class3d w. alignment unmasked
reference
(30 Å LP)
15.9%
28.0%
35.7%
5.0%
15.4%
~90°
~90°
refine3d 5 Å (masked)
re-extract to increase box size (432 to 512 pix.)
class2D
178.778 ptcls. (1.25 Å/pix)
refine3d 4.4 Å (masked)
re-extract
class2D
178.786 ptcls. (1.25 Å/pix)
refine3d 4.6 Å (masked)
class3d w/o alignment
masked
47.0%
5.0%
29.7%
12.0%
6.3%
volume re-centered on E27
3D reference for Relion autopicking (20 Å LP)
Polara data sets
20dec21a, 21jan11a, 21feb08a, 21may03a
4x class2D
974.291 ptcls. (2.5 Å/pix)
removal of
ArnA contamination:
132.862 ptcls.
class3d w. alignment unmasked
reference
(25 Å LP)
39.0%
11.5%
18.6%
12.0%
18.9%
re-extract
class2D
389.784 ptcls. (1.25 Å/pix)
refine3d 4.1 Å (masked)
refine3d 5.1 Å (masked)
Map 1
export to Cryosparc
ab initio cleanup
(2 classes, similarity = 0)
241.213 ptcls. (1.25 Å/pix)
non-uniform refinement
GSFSC Resolution: 3.78Å
No Mask (4.4Å)
Spherical (4.1Å)
Loose (3.9Å)
Tight (3.7Å)
Corrected (3.8Å)
DC 16Å 8Å 5.3Å 4Å 3.2Å 2.7Å
Elevation
Azimuth
### of images
3 4 5 6 7 9 11 Å
refine3d 8.2 Å (masked)
class3d w. alignment unmasked
47.2%
8.0%
1.8%
18.9%
24.1%
refine3d 6.3 Å (masked)
Map 2
GSFSC Resolution: 4.79Å
No Mask (7.9Å)
Spherical (7.2Å)
Loose (6.2Å)
Tight (4.7Å)
Corrected (4.8Å)
DC 16Å 8Å 5.3Å 4Å 3.2Å 2.7Å
Elevation
Azimuth
### of images
local refinement
full mask
fulcrum: STAT2
3 4 5 6 7 9 11 Å
44.7%
55.3%
reference
reference
heterogeneous
refinement
23.1%
53.0%
23.9%
3D variability
3 clusters
re-extract
71.534 ptcls. (1.25 Å/pix)
export to Cryosparc
non-uniform refinement: 4.4 Å

**Figure S3: Cryo-EM data processing flowchart, related to Figures 3-5**

If not stated otherwise, all computational procedures have been performed using the program Relion 3.0.7 (Zivanov et al., 2018). A crystal structure of hexameric ArnA, PDB 4wkg (Fischer et al., 2015; Yang et al., 2019), was fitted in the contamination density (red box).

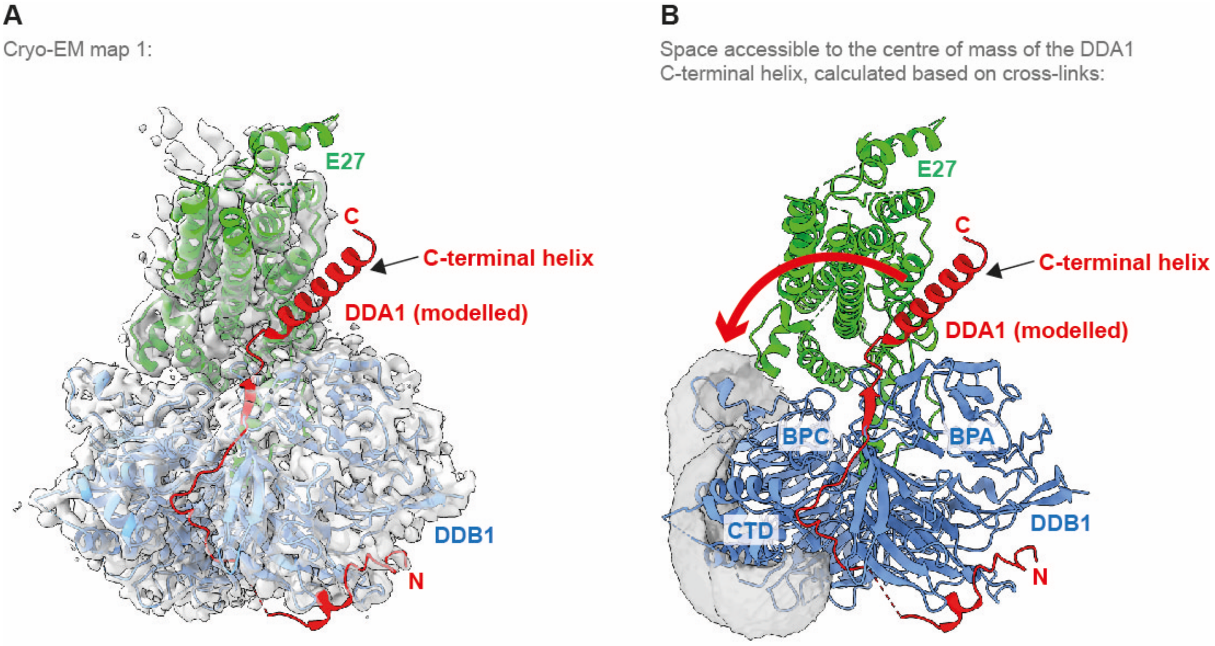


**Figure S4: Molecular modelling of DDA1, related to Figure 3**

**(A)** Superposition of a $DDB1_{\Delta BPB}$/DDA1/DCAF15 complex structure (PDB 6q0v) (Faust et al., 2020) with $DDB1_{\Delta BPB}$/E27 complex coordinates. Proteins are shown as cartoons, DDB1 is coloured in blue, E27 in green, and DDA1 in red. DCAF15 has been removed for clarity, and only DDB1 from the DDB1/E27 structure is shown. DDA1 termini and the C-terminal helix are indicated. Experimental cryo-EM map 1 is shown as semi-transparent surface.

**(B)** Visualisation (grey semi-transparent surface) of the space accessible to the centre of mass of the DDA1 C-terminal helix, calculated based on cross-link distance restraints, consistent with at least 15 of the observed cross-links involving DDA1, using the DisVis software tool (van Zundert and Bonvin, 2015; van Zundert et al., 2017). Proteins are represented as in **A**. This analysis implies that the C-terminal helix preferentially locates to a relatively large area around the edges of DDB1 BPC and CTD, and does not occupy a defined location in contact with E27, as observed in the $DDB1_{\Delta BPB}$/DDA1/DCAF15 complex (indicated by the red arrow).

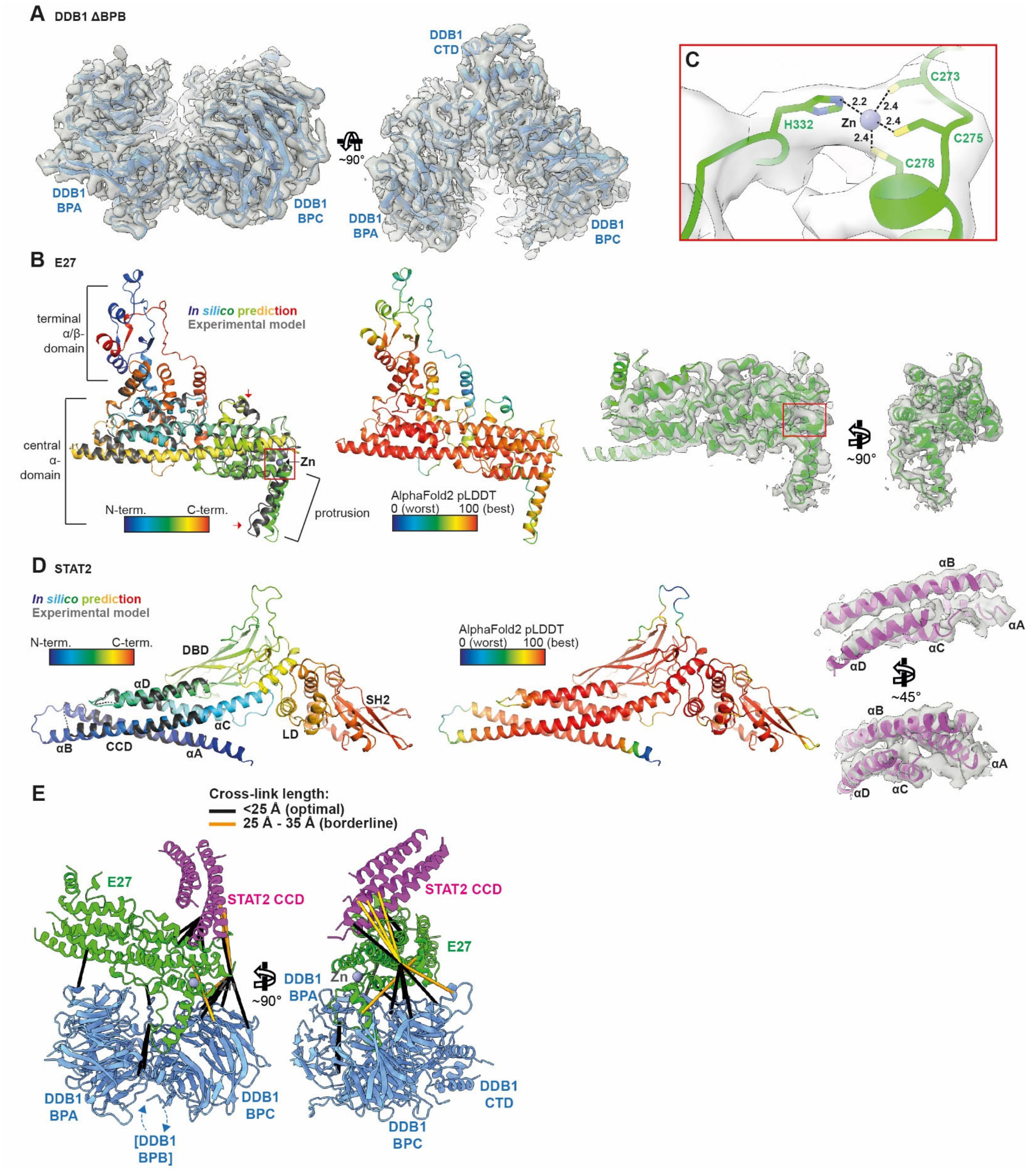


**Figure S5: Cryo-EM map 1 quality, AlphaFold2 models, model fit, and CLMS analysis, related to Figure 3**

**(A)** Overview of DDB1$_{\Delta BPB}$ model-to-map fit. DDB1$_{\Delta BPB}$ is shown as blue cartoon, cryo-EM density as grey semi-transparent surface (threshold level 0.555, Chimerax software tool) (Pettersen et al., 2021).

**(B) (Left panel)** Superposition of the AlphaFold2 E27 model (rainbow-coloured from N- to C-terminus) with a partial structure (grey) manually built into cryo-EM map 1 (Fig. S3). Structures are shown as cartoons. Terminal regions of the AlphaFold2 model with very low confidence have been removed for clarity. Red arrows indicate regions with significant structural differences between predicted and experimental model. The black arrow and the red box indicate the zinc-binding motif. The zinc ion is shown as grey sphere. **(Middle panel)** Cartoon representation of the E27 AlphaFold2 model, coloured according to model confidence, i.e., the model's prediction of its score on the local Distance Difference Test (lDDT-Cα) (Jumper et al., 2021). Terminal regions with very low confidence have been removed from the model for clarity. **(Right panel)** Overview of E27 model-to-map fit. E27 is shown as green cartoon, cryo-EM density as grey semi-transparent surface (threshold level 0.5, Chimerax software tool) (Pettersen et al., 2021).

**(C)** Close-up view of the zinc-binding site marked as red box in **B**. E27 is shown as green cartoon, with side chains involved in zinc coordination in stick representation. The zinc ion is shown as grey sphere. Coordination bond lengths (in Å) are indicated. Cryo-EM density is shown as grey semi-transparent surface (threshold level 0.5, Chimerax software tool) (Pettersen et al., 2021). Note that the tetrahedral coordination geometry, the ligands (Cys, His), and the coordination bond lengths are typical for structural zinc-binding sites, as demonstrated by analysis of almost 25,000 zinc-binding motifs available in the PDB (Ireland and Martin, 2019; Laitaoja et al., 2013).

**(D) (Left panel)** Superposition of the AlphaFold2 STAT2 model (rainbow-coloured from N- to C-terminus) with the portion of the STAT2 CCD (grey) fitted into cryo-EM map 1 (Fig. S3). Structures are shown as cartoons. ND and TAD have been removed for clarity. **(Middle panel)** Cartoon representation of the STAT2 AlphaFold2 model, coloured according to model confidence, i.e., the model's prediction of its score on the local Distance Difference Test (lDDT-Cα) (Jumper et al., 2021). **(Right panel)** Overview of STAT2 CCD model to map fit. STAT2 is shown as magenta cartoon, cryo-EM density as grey semi-transparent surface (threshold level 0.35, Chimerax software tool) (Pettersen et al., 2021).

**(E)** Intermolecular sulfo-SDA cross-links were mapped on the DDB1$_{\Delta BPB}$/E27/STAT2 CCD structure (Fig. 3A), colour-coded according to their length as indicated.

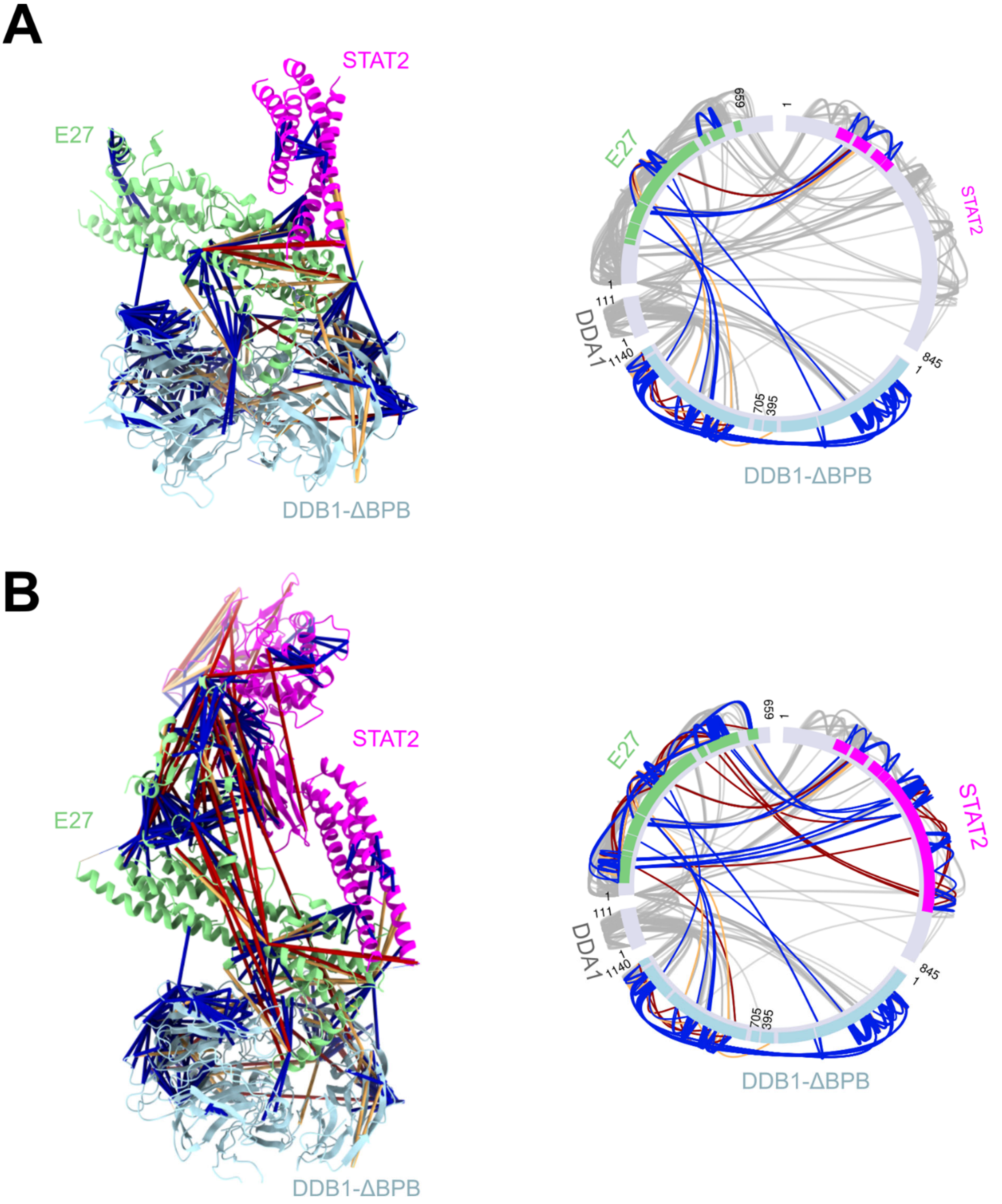


**Figure S6: CLMS analysis of DDB1/E27/STAT2 models derived from cryo-EM map 1 and map 2, related to Figures 3 and 5**

(**A**) Left panel: sulfo-SDA cross-links mapped onto the DDB1/E27/STAT2 model built in map 1. Both within protein and heteromeric links are displayed. Satisfied cross-links (shorter than 25 Å Cα-Cα) are coloured in blue, borderline violations (25-35 Å) in orange, and violated (> 35 Å) cross-links in red. Right panel: circle view of the links between proteins coloured by cross-link satisfaction. Grey cross-links lie between regions not modelled in the structure. Solid coloured bars indicate structural coverage.

(**B**) As in **A**, but for the model built in map 2. Violated cross-links between E27 α/β-domain, STAT2 and DDB1 may indicate a swivel motion of the top of the complex relative to DDB1, which is consistent with the high mobility of this module relative to the core of the complex observed in cryo-EM analysis.

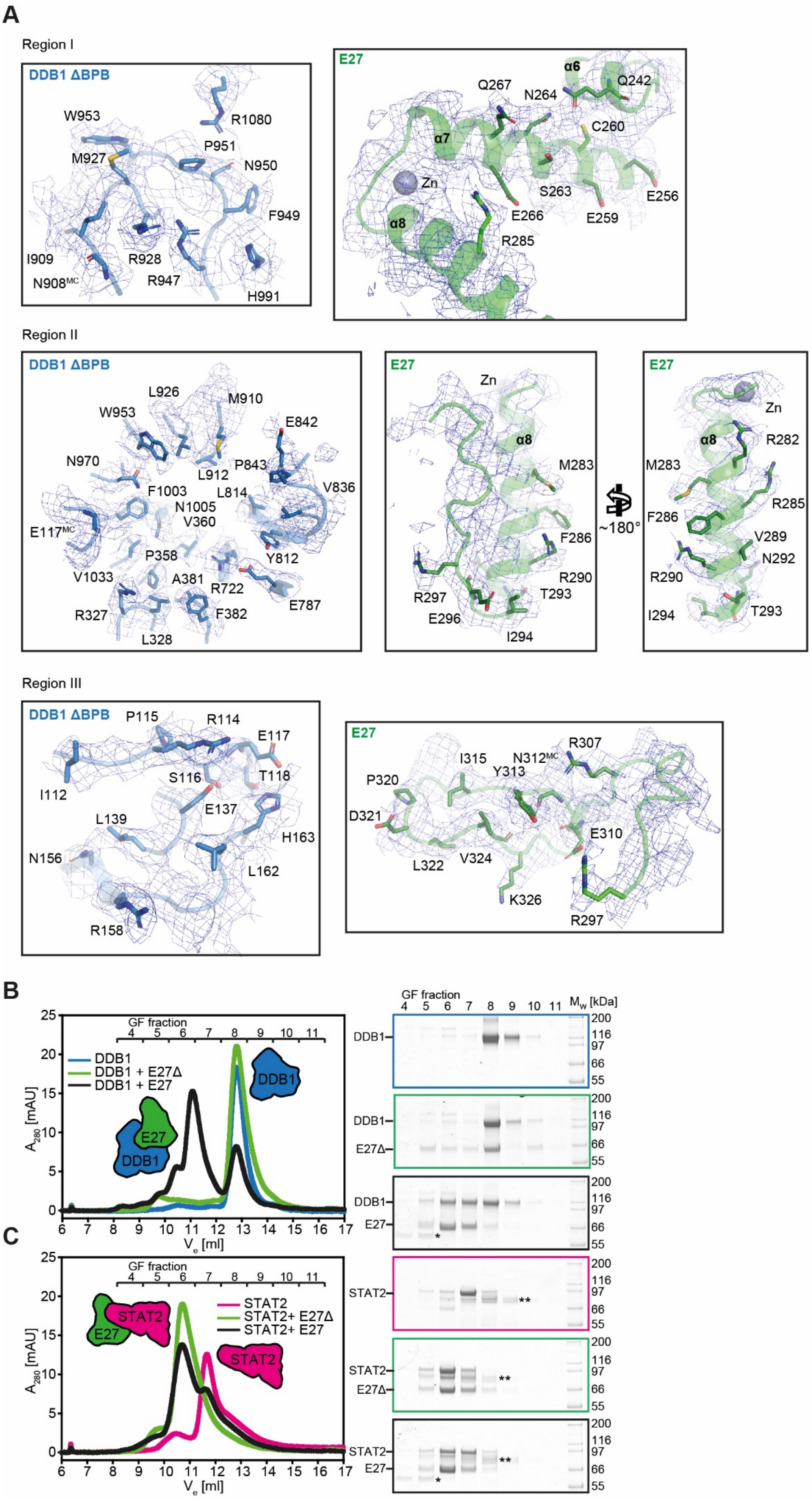
A
Region I
DDB1 ΔBPB
W953
R1080
P951
M927
N950
F949
I909
R928
N908MC
R947
H991
E27
α6
Q267
N264
Q242
C260
α7
Zn
S263
E256
E266
E259
α8
R285
Region II
DDB1 ΔBPB
L926
M910
W953
E842
N970
L912
P843
F1003
L814
V836
N1005
V360
E117MC
P358
Y812
V1033
A381
R722
E787
R327
F382
L328
E27
Zn
α8
M283
F286
R290
R297
T293
E296
I294
~180°
E27
Zn
α8
R282
M283
R285
F286
V289
N292
R290
I294
T293
Region III
DDB1 ΔBPB
P115
R114
E117
S116
T118
I112
E137
L139
H163
N156
L162
R158
E27
R307
I315
N312MC
Y313
P320
D321
E310
L322
V324
K326
R297
B
GF fraction
4 5 6 7 8 9 10 11
DDB1
DDB1 + E27Δ
DDB1 + E27
DDB1
E27
DDB1
A280 [mAU]
Ve [ml]
GF fraction
4 5 6 7 8 9 10 11
Mw [kDa]
DDB1
E27Δ
E27
200
116
97
66
55
*
C
GF fraction
4 5 6 7 8 9 10 11
STAT2
STAT2+ E27Δ
STAT2+ E27
STAT2
E27
STAT2
A280 [mAU]
Ve [ml]
STAT2
E27Δ
E27
**
*

**Figure S7: Side chain densities and mutational analysis of the $DDB1_{\Delta BPB}$/E27 interaction, related to Figure 3**

**(A)** Model-to-map fit of the $DDB1_{\Delta BPB}$/E27 interaction regions (Fig. 3B). $DDB1_{\Delta BPB}$ is shown as blue cartoon, with side chains involved in E27 interaction in stick representation. E27 is shown as green cartoon, with side chains mediating $DDB1_{\Delta BPB}$ interaction in stick representation. Cryo-EM density map 1 (Fig. S3) is shown as blue mesh, contoured at 5σ, using the the PyMOL Molecular Graphics System, Version 2.0 Schrödinger, LLC.

**(B)** *In vitro* reconstitution, followed by analytical gel filtration (GF) analysis, of protein complexes containing DDB1 and E27 or an E27 variant, where residues E284-R305 have been replaced by a GNGNSG-linker (E27Δ). SDS-PAGE analyses of GF fractions are shown next to the chromatogram. * - contaminant from E27 preparations.

**(C)** *In vitro* reconstitution, followed by GF analysis, of protein complexes containing STAT2 and E27 or E27Δ. SDS-PAGE analyses of GF fractions are shown as in **D**. ** - contaminant from STAT2 preparation.

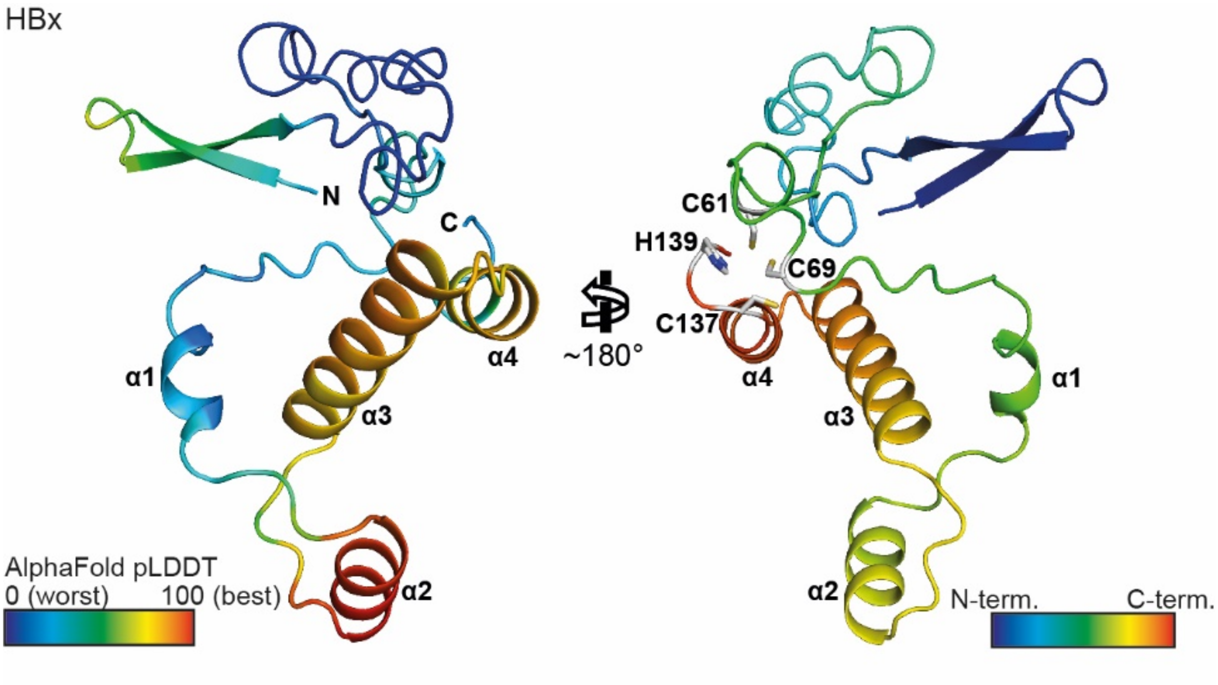


**Figure S8: AlphaFold2 prediction of HBx in complex with DDB1$_{\Delta BPB}$, related to Figure 4**

HBx is shown in cartoon representation. DDB1 has been removed for clarity. On the left panel, the model is coloured according to model confidence, i.e., the model's prediction of its score on the local Distance Difference Test (lDDT-Cα) (Jumper et al., 2021). On the right panel, the model is rainbow-coloured from N- to C-terminus, and side chains involved in zinc coordination (Ramakrishnan et al., 2019) are shown as sticks.

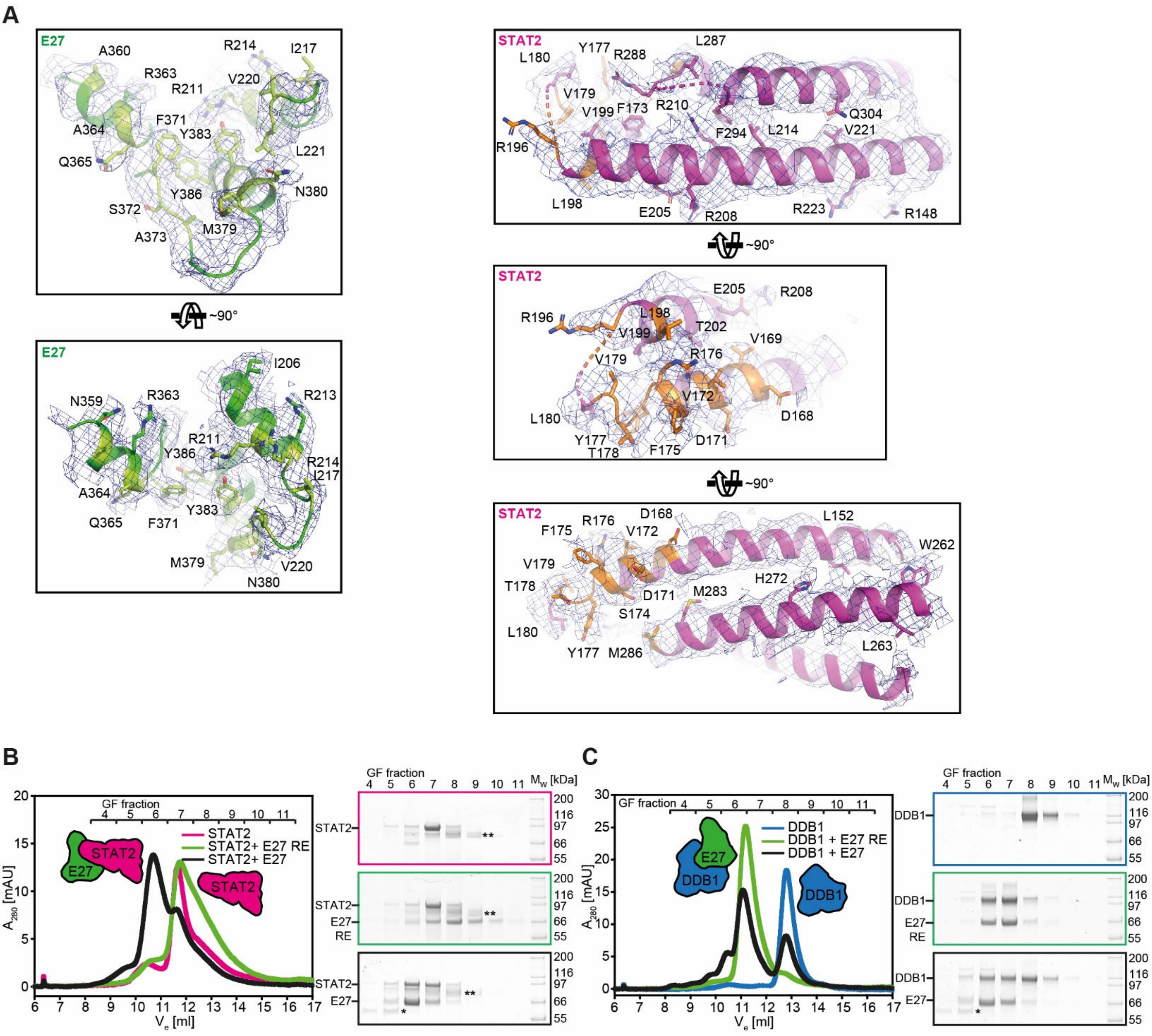


**Figure S9: Side chain densities and mutational analysis of the E27/STAT2 interaction, related to Figure 5**

**(A)** Model-to-map fit of the E27/STAT2 interaction (Fig. 5A). E27 is shown as green cartoon, with selected side chains shown as sticks, and side chains involved in STAT2 interaction marked in light green. STAT2 is shown as magenta cartoon, with selected side chains shown as sticks, and side chains involved in E27 interaction marked in orange. Cryo-EM density map 1 (Fig. S3) is shown as blue mesh, contoured at 4σ, using the the PyMOL Molecular Graphics System, Version 2.0 Schrödinger, LLC.

**(B)** *In vitro* reconstitution, followed by GF analysis, of protein complexes containing STAT2 and E27 or the E27 R211E/R214E double mutant (E27-RE). SDS-PAGE analyses of fractions collected during the GF runs are shown next to the chromatogram. * - contaminant from E27 preparations, ** - contaminant from STAT2 preparations.

**(C)** *In vitro* reconstitution, followed by GF analysis, of protein complexes containing DDB1 and E27 or E27-RE. SDS-PAGE analyses of fractions collected during the GF runs are shown next to the chromatogram as in **B**.

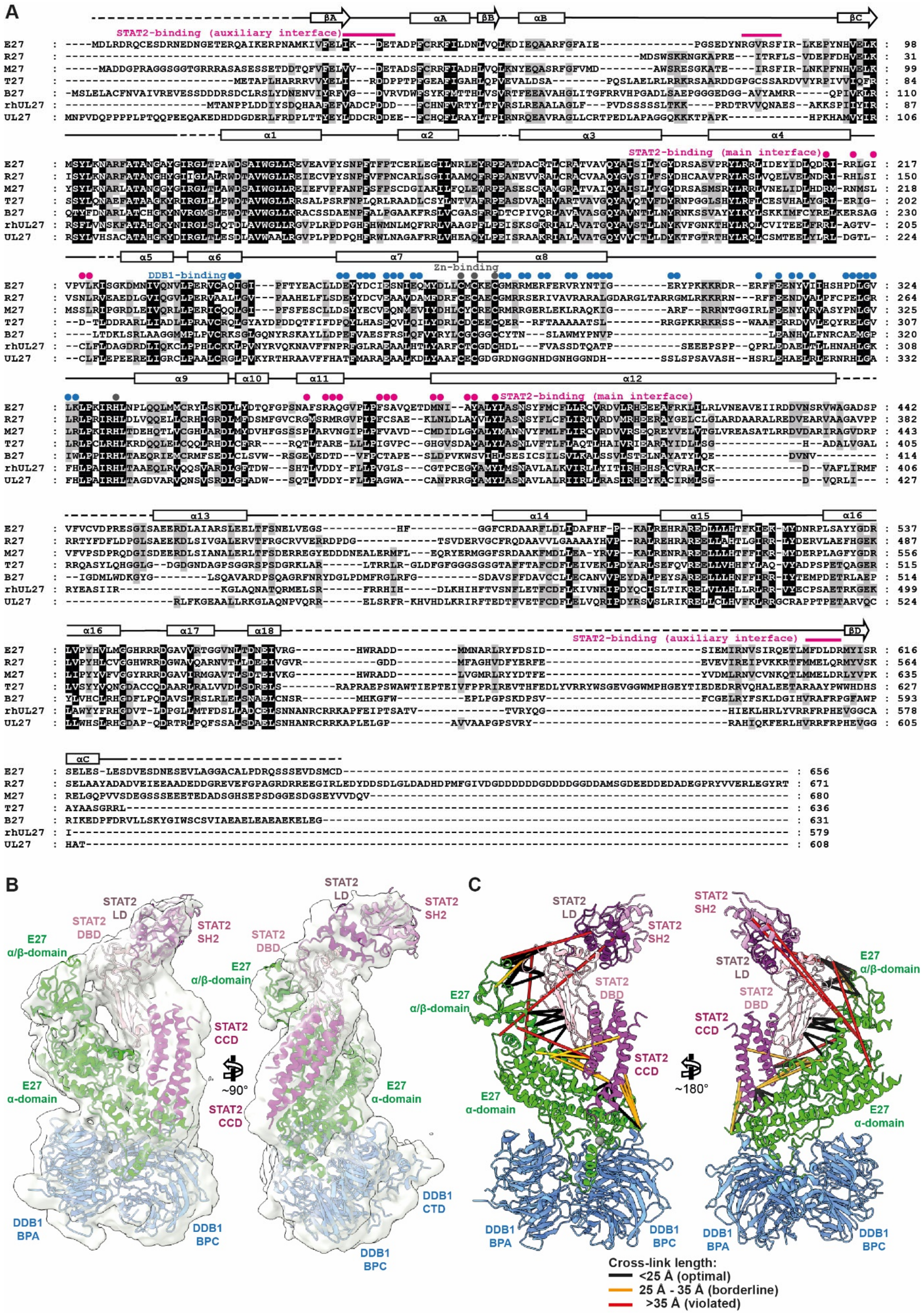
A
STAT2-binding (auxiliary interface)
STAT2-binding (main interface)
DDB1-binding
Zn-binding
E27
R27
M27
T27
B27
rhUL27
UL27
B
STAT2 LD
STAT2 DBD
STAT2 SH2
E27 α/β-domain
STAT2 CCD
E27 α-domain
DDB1 BPA
DDB1 BPC
DDB1 CTD
~90°
C
~180°
Cross-link length:
<25 Å (optimal)
25 Å - 35 Å (borderline)
>35 Å (violated)

**Figure S10: E27 multiple sequence alignment, cryo-EM map 2 model fit, and CLMS analysis, related to Figure 5**

**(A)** Multiple sequence alignment of E27 and its homologs from other CMV species. >80% type-conserved residues are highlighted black, >60% type-conserved residues grey. Secondary structure elements are drawn schematically above the alignment. Dashed lines indicate flexible regions not included in the molecular model. Residues involved in DDB1-binding and in the main STAT2-binding interface are indicated by blue and magenta dots above the alignment, respectively. Residue stretches involved in the auxiliary E27 α/β-domain/STAT2 DBD binding interface are indicated by magenta lines above the alignment. Side chains involved in zinc-coordination are marked by grey dots.

**(B)** Overview of DDB1$_{\Delta BPB}$/E27/STAT2 model fit into cryo-EM map 2 (Fig. S3). DDB1$_{\Delta BPB}$ is shown as blue cartoon, E27 as green cartoon, and STAT2 as magenta cartoon. Cryo-EM density is shown as grey semi-transparent surface (threshold level 0.15, Chimerax software tool) (Pettersen et al., 2021).

**(C)** Intermolecular sulfo-SDA cross-links were mapped on the DDB1$_{\Delta BPB}$/E27/STAT2 structure (Fig. 5C), colour-coded according to their length as indicated.

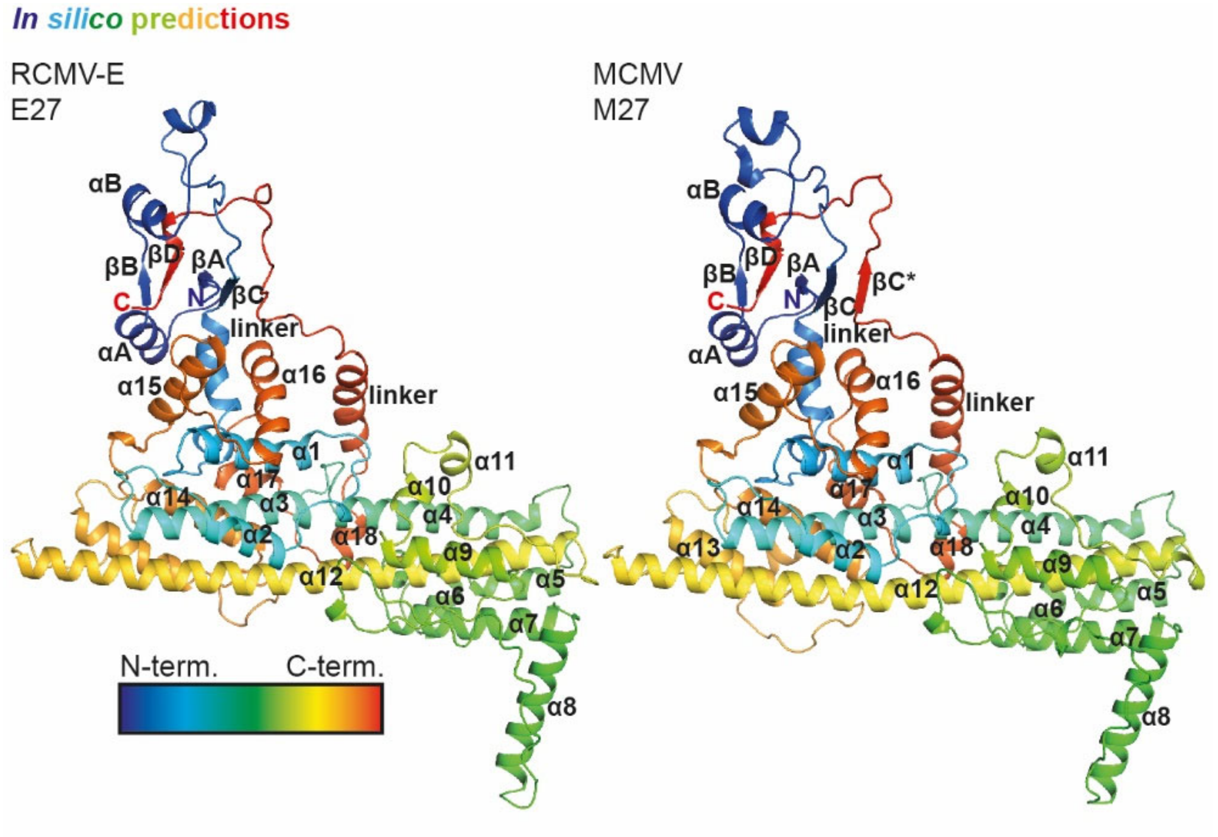


**Figure S11: Comparison of E27 and M27 AlphaFold2 predictions, related to Figure 7**

Cartoon representation of AlphaFold2 models of E27 (left) and M27 (right), rainbow-coloured from N- to C-terminus. Terminal regions with very low confidence have been removed. Termini and secondary structure elements are labelled.

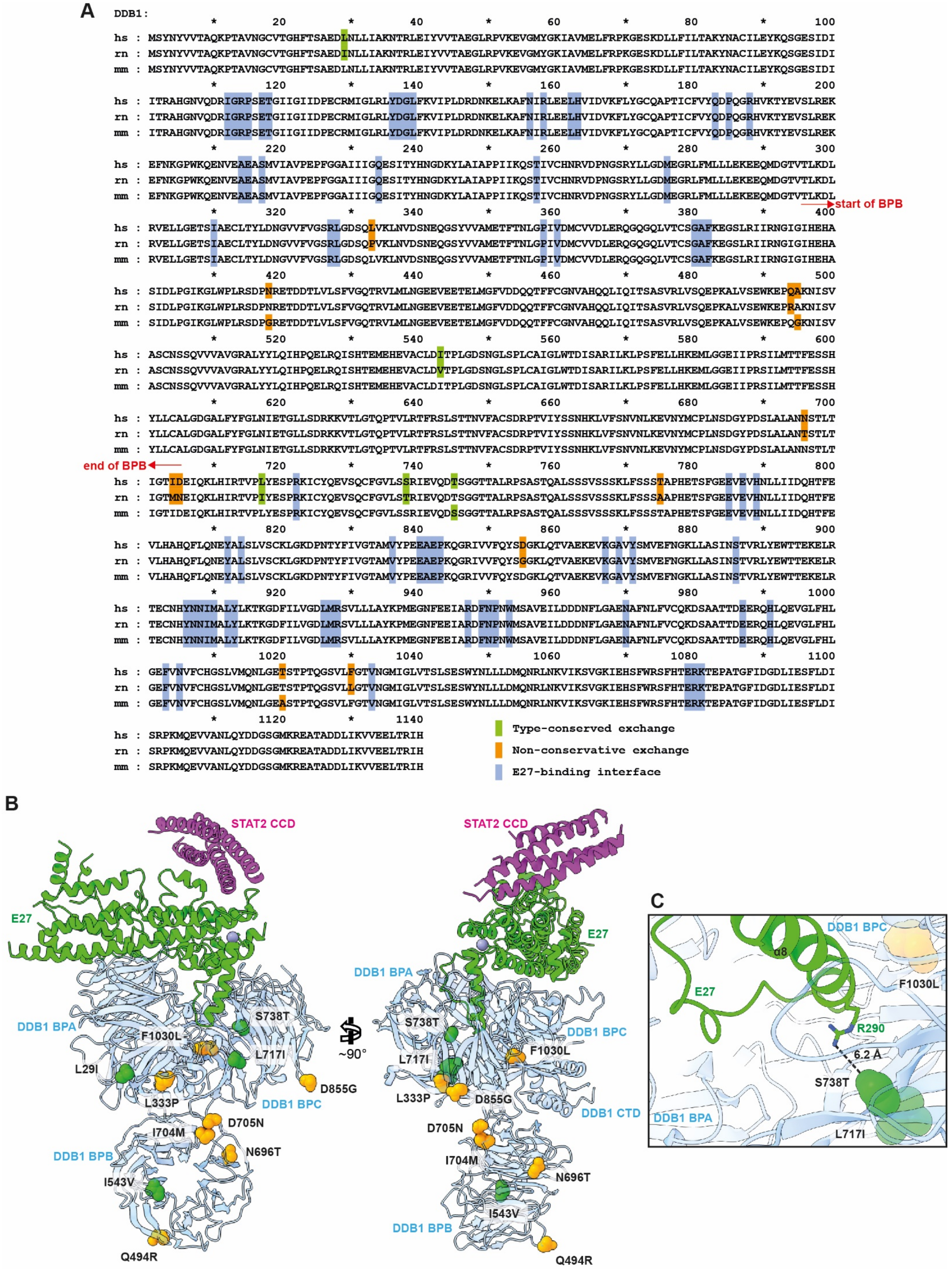

A
DDB1:
hs
rn
mm
start of BPB
end of BPB
Type-conserved exchange
Non-conservative exchange
E27-binding interface
B
STAT2 CCD
E27
DDB1 BPA
DDB1 BPB
DDB1 BPC
DDB1 CTD
S738T
F1030L
L717I
L29I
L333P
D855G
D705N
I704M
N696T
I543V
Q494R
~90°
C
DDB1 BPC
α8
E27
R290
6.2 Å
DDB1 BPA

**Figure S12: Multiple sequence alignment of human, rat and mouse DDB1, visualisation of non-conserved residues, related to Discussion**

**(A)** Multiple sequence alignment of human (hs), rat (rn) and mouse (mm) DDB1. Non-synonymous but type-conserved amino acid exchanges between the species are highlighted in green, non-conserved exchanges in red. Residues involved in the $DDB1_{\Delta BPB}$/E27 interaction interface are highlighted in blue.

**(B)** Amino acid exchanges from (A) have been mapped on a structural model of the DDB1/E27/STAT2 CCD complex. This model has been generated by superposition of a full-length DDB1 structure (PDB 2b5m (Li et al., 2006)) with $DDB1_{\Delta BPB}$ from the complex structure presented here (Fig. 3). Only the coordinates of full-length DDB1 are shown for clarity. DDB1 is shown as semi-transparent blue cartoon, with the BPs and the CTD indicated. Non-conserved residues from **A** are shown as spheres using the same colour code as in **A**.

**(C)** Close-up view of the E27-binding cavity as shown in **B**. The closest distance of an amino acid exchange between human and rat DDB1 to an E27 side chain, DDB1 L738T to E27 R290, has been measured as 6.2 Å, more than twice the length of a typical hydrogen bond, strongly indicating that amino acid exchanges between human and rat DDB1 residues do not affect E27 binding in any way.

**Table S1: Cryo-EM data collection and refinement**

#### Data collection and processing

| | |
|---|---|
| Magnification | 31.000 |
| Voltage | 300 |
| Electron exposure [e/Å$^2$] | 62 |
| Defocus range [µm] | 0.5-2.5 |
| Pixel size [Å] | 0.625 |
| Initial particle no. | 974.291 |
| Final particle no. | 389.784 (map 1); 31.976 (map 2) |
| Map resolution [Å] | 3.8 (map 1); 4.8 (map 2) |
| FSC threshold | 0.143 |
| Map sharpening B factor [Å$^2$] | 165.8 (map 1); 177.4 (map 2) |

#### Refinement

| Model | DDB1$_{\Delta BPB}$/E27 (map 1) | DDB1$_{\Delta BPB}$/E27/STAT2 (map 2) |
|---|---|---|
| Model fit CCs - mask/box/peaks/volume | 0.76/0.75/0.67/0.75 | 0.71/0.87/0.62/0.71 |
| Model resolution - masked/unmasked | 4.0/4.1 (3.7/3.8) | 8.9/9.0 (6.3/6.5) |
| FSC threshold | 0.5 (0.143) | 0.5 (0.143) |
| Model composition | | |
| Nonhydrogen atoms | 10497 | 14734 |
| Ligand atoms | 1 | 1 |
| Average B factors [Å$^2$] | | |
| Protein | 106.71 | 370.21 |
| Ligand | 101.44 | 235.31 |
| R.m.s. deviations | | |
| Bond lengths [Å] | 0.006 | 0.003 |
| Bond angles [°] | 1.572 | 0.776 |
| Validation | | |
| MolProbity score | 1.98 | 1.82 |
| Clashscore | 11.99 | 6.6 |
| Poor rotamers [%] | 0.51 | 0.99 |
| Ramachandran plot | | |
| Favoured [%] | 94.3 | 92.7 |
| Allowed [%] | 5.7 | 7.1 |
| Disallowed [%] | 0 | 0.2 |

**Table S2: Crosslink violation analysis.**

| Model | Crosslink type | Number of crosslinks in structure | Satisfied (<25 Å Cα-Cα) | Borderline (25-35 Å Cα-Cα) | Violated (>35 Å Cα-Cα) | Violation % |
|---|---|---|---|---|---|---|
| Map 1 | Self & heteromeric | 244 | 208 | 26 | 10 | 4.9% |
| Map 1 | Heteromeric only | 21 | 16 | 5 | - | - |
| Map 2 | Self & heteromeric | 432 | 348 | 48 | 36 | 8.3% |
| Map 2 | Heteromeric only | 51 | 33 | 12 | 6 | 11.8% |

The total number of unambiguous cross-linked residue pairs in the dataset is 918, with 219 heteromeric links.

**Table S3: Key resources**

| REAGENT or RESOURCE | SOURCE | IDENTIFIER |
|---|---|---|
| Antibodies | | |
| Mouse monoclonal anti-FLAG (clone M2) | Sigma-Aldrich | Cat# F3165; RRID:AB_259529 |
| Rabbit polyclonal anti-DDB1 | Bethyl Laboratories | Cat# A300-462A |
| Rabbit polyclonal anti-STAT2 | Santa Cruz Biotechnology | Cat# sc-950; RRID:AB_2271322 |
| Mouse monoclonal anti-MCMV-pp89 (CROMA101) | Stipan Jonjic (Rijeka, Croatia) | N/A |
| Rabbit polyclonal anti-GAPDH | Santa Cruz Biotechnology | Cat# sc-25778; RRID:AB_10167668 |
| Rabbit polyclonal β-Tubulin Antibody | Cell Signaling Technology Inc. | Cat# 2146 |
| Rabbit monoclonal Recombinant Anti-STAT2 [Y141] | Abcam plc. | Cat# ab32367 |
| Mouse monoclonal anti-RCMV-E IE1 | Stipan Jonjic (Rijeka, Croatia) | N/A |
| Bacterial and virus strains | | |
| E. coli NovaBlue | Merck KGaA | Cat# 70181 |
| E. coli Rosetta$^{TM}$ 2(DE3) | Merck KGaA | Cat# 71400 |
| Autographa californica nucleopolyhedrovirus clone C6 | (Zhao et al., 2003) | |
| WT-MCMV | (Jordan et al., 2011) | N/A |
| ΔM27-MCMV | (Le-Trilling et al., 2018) | N/A |
| MCMV$_{AxAxxA}$ | This work | N/A |
| WT-RCMV-E | (Ettinger et al., 2012) | RCMV isolated from Rattus norvegicus |
| ΔE27-RCMV-E | This work | CRISPR/Cas9 generated E27 deletion mutant |
| Chemicals, peptides, and recombinant proteins | | |
| Recombinant mouse IFN-β | PBL | Cat# 12405-1 |
| Recombinant mouse IFN-λ2 | Peprotech | Cat# 300-02K |
| MLN-4924 (Pevonedistat) | Active Biochem | Cat# A-1139 |
| MG 132 | Sigma-Aldrich | Cat# 474790 |
| Trypsin Sequencing Grade | SERVA Electrophoresis | Cat# 37283 |
| RapiGest SF Surfactant | Waters | Cat# 186002123 |
| Critical commercial assays | | |
| QuikChange II XL Site Directed Mutagenesis Kit | Agilent | Cat# 200522 |
| QIAprep Spin Miniprep Kit | Qiagen | Cat# 27106X4 |
| Deposited data | | |
| Cryo-EM map 1 | This work | EMD-14802 |
| Cryo-EM map 2 | This work | EMD-14812 |
| DDB1$_{\Delta BPB}$/E27/STAT2 CCD complex coordinates | This work | PDB 7zn7 |
| DDB1$_{\Delta BPB}$/E27/STAT2 FL complex coordinates | This work | PDB 7znn |
| LC-MS data | This work | PRIDE PXD033639 |
| CLMS data | This work | JPOST ProteomeXChange JPST001566 |
| Experimental models: Cell lines | | |
| Sf9 | ThermoFisher | Cat# 11496015 |
| HEK293T | ATCC | Cat# CRL-11268 |
| Immortalised mouse fibroblasts | (Rattay et al., 2015) | N/A |

| | | |
|---|---|---|
| NIH3T3-ISREluc:IFNLR | (Le-Trilling et al., 2018) | N/A |
| Rat embryo fibroblasts (REF) | Laboratory of William Burns | N/A |
| Experimental models: Organisms/strains | | |
| BALB/c mice | Harlan | N/A |
| Recombinant DNA | | |
| pHisSUMO-E27 | This work | |
| pHisSUMO vector | Structural Biology STP, Francis Crick Institute, London, UK | N/A |
| pAcGHLT-B-*Homo sapiens* (hs) DDB1 | Addgene | Cat# 48638 |
| pAcGHLT-B-3C-hsDDB1 | This work | N/A |
| pAcGHLT-B-3C-hsDDB1 ΔBPB | This work | N/A |
| pHisSUMO-*Rattus norvegicus* (rn) STAT2 | This work | N/A |
| pTriEx-6-hsDCAF1 1046-1396 | (Banchenko et al., 2021) | N/A |
| pIRES2-EGFP-M27FLAG | (Trilling et al., 2011) | N/A |
| pIRES2-EGFP-M27FLAG-AxAxxA | (Trilling et al., 2011) | N/A |
| pIRES2-EGFP-M27FLAG-Y270F | This work | N/A |
| pIRES2-EGFP-M27FLAG-LP328/329AA | This work | N/A |
| pIRES2-EGFP-M27FLAG-RHL332-334AA | This work | N/A |
| R-CRISPR-1003 (pSicoR-CRISPR-PuroR vector carrying a TCGGTCCGACTCGCACTGG crRNA) | This work | N/A |
| R-CRISPR-1004 (pSicoR-CRISPR-PuroR vector carrying a AGAGACCGCCAGTGCGAGT crRNA) | This work | N/A |
| R-CRISPR-1005 (pSicoR-CRISPR-PuroR vector carrying a CTGACGAACTCTGTCGATC crRNA) | This work | N/A |
| R-CRISPR-1006 (pSicoR-CRISPR-PuroR vector carrying a TGCGACTGAACAAACGAAT crRNA) | This work | N/A |
| Software and algorithms | | |
| ProtParam | (Wilkins et al., 1999) | N/A |
| Leginon | (Suloway et al., 2005) | N/A |
| Relion 3.0.7 | (Zivanov et al., 2018) | N/A |
| cryoSPARC | (Punjani et al., 2017) | N/A |
| Coot 0.9.5 | (Emsley et al., 2010) | N/A |
| AlphaFold2 | (Jumper et al., 2021) | N/A |
| Phenix 1.19.2-4158 | (Afonine et al., 2018) | N/A |
| GraphPad Prism | GraphPad Software Inc. | N/A |
| MicroWin (Mithras2 LB 943) | Berthold Technologies | N/A |
| Progenesis QI for Proteomics | Nonlinear Dynamics | N/A |
| Proteome Discoverer 1.4 | Thermo Scientific | N/A |
| Other | | |
| LB broth | Carl Roth | Cat# X968.2 |
| Insect-XPRESS$^{TM}$ Protein-free Insect Cell Medium | Lonza | Cat# BELN12-730Q |

**Table S4: Oligonucleotide primer sequences**

| Name | Sequence |
|---|---|
| XmaI-E27-fw | GGCCCCGGGATGGATCTCAGAGACCGCCAGTG |
| SacI-E27-rev | GGCGAGCTCTTATTAGTCGCACATCGAATCGACTTCTG |
| 3C-DDB1-A | CACGTAGTTGTACGACATCCCGGGTCCCTGAAAGAGGA CTTCAAGAATTCCCGCCGCAACACTGGCC |
| 3C-DDB1-B | GTGTTGCGGCGGGAATTCTTGAAGTCCTCTTTCAGGGAC CCGGGATGTCGTACAACTACGTGGTAAC |
| ΔBPB-A | GCAGCTTCTGGATCTCTCCAGAGTTACCGTTTCCTCCAA TTCCATTCCGGATGATCCG |
| ΔBPB-B | GCGGAATGGAATTGGAGGAAACGGTAACTCTGGAGAGA TCCAGAAGCTGCACATTCG |
| XmaI-STAT2-fw | CCCCCCGGGATGGCGCAGTG |
| NotI-STAT2-rev | TAAAGCGGCCGCTTAGTCTTCAG |
| E27Δ-A | CTCAAAAAATCTTTCTCCAGAGTTACCGTTTCCCATTCTGCGCATTCCACATTCTTTAC |
| E27Δ-B | GGAATGCGCAGAATGGGAAACGGTAACTCTGGAGAAAGATTTTTTGAGGAGAACTAC |
| E27-R211E/R214E-fw | CTGCAAGATGAAATACGGGAGCTCGGTATAGTACCCGTATTG |
| E27-R211E/R214E-rev | TATACCGAGCTCCCGTATTTCATCTTGCAGGTCTATATACTC |
| Seq_E27_CRISPR_fwd | CAACCTGCAGCCGTATAG |
| Seq_E27_CRISPR_rev | CGCAACCGACGTAATACAC |
| M27-QC1-3 | CTACGATCAACTCGCGTACGCGCGCGAGGCGCGGATGCGCCGGG |
| M27-QC1-3-rc | CCCGGCGCATCCGCGCCTCGCGCGCGTACGCGAGTTGATCGTAG |
| M27-QC-LP | CAACATCGGCGTGCTGCGCGCGGCCAAGATCCGCCACCTCAGCG |
| M27-QC-LP-rc | CGCTGAGGTGGCGGATCTTGGCCGCGCGCAGCACGCCGATGTTG |
| M27-QC-RHL | GCTGCGCCTGCCCAAGATCGCCGCCGCCAGCGACGAACACCAGACCC |
| M27-QC-RHL | GCTGCGCCTGCCCAAGATCGCCGCCGCCAGCGACGAACACCAGACCC |
| MCMV-M27-QC1-3_1 | GTACGTCGAGATGACCATGGAGGTCATCTACGATCAACTCGCGTACGCGCGCGA GGCGCGGATGCGCCGGGGCGAGCGAGGATGACGACGATAAGTAGGG |
| MCMV-M27-QC1-3_2 | GCGCGCGGAGCCTCTCGAGGCGCTCGCCCCGGCGCATCCGCGCCTCGCGCGC GTACGCGAGTTGATCGTAGATGACCTCAACCAATTAACCAATTCTGATTAG |

#### Supplementary references


Afonine, P.V., Poon, B.K., Read, R.J., Sobolev, O.V., Terwilliger, T.C., Urzhumtsev, A., and Adams, P.D. (2018). Real-space refinement in PHENIX for cryo-EM and crystallography. Acta Crystallogr D Struct Biol *74*, 531-544.
